# Comparison of the Distribution of Fitness Effects Across Primates

**DOI:** 10.64898/2026.03.25.714151

**Authors:** Janek Sendrowski, Bjarke M. Pedersen, Juraj Bergman, Vasili Pankratov, Thomas Bataillon

## Abstract

The distribution of fitness effects (DFE) of mutations is a key determinant of both the efficacy of natural selection and the genetic load of populations. It also provides an indirect summary of the underlying fitness landscape, so the extent to which the DFE is conserved across species is informative about the invariance of those landscapes. Here, we infer the DFE of amino-acid–changing mutations in 38 catarrhine subspecies using site-frequency spectrum (SFS)–based methods. We find that effective population size (*N_e_*) is the dominant axis of cross-species variation in the population-scaled DFE for deleterious mutations, with the apparent clade-level clustering of estimates explained mainly by shared *N_e_* rather than by clade-specific effects. A quantile-based test further shows that the underlying unscaled DFE does not vary significantly across species, so differences are consistent with *N_e_*-rescaling of a single conserved DFE. Consistent with this, the deleterious (non-adaptive) substitution rate *ω_na_*declines significantly with *N_e_*, reflecting more efficient purging of slightly deleterious mutations in larger populations. Turning to adaptive substitution, we find weaker, suggestive evidence that the adaptive rate *ω_a_* increases with *N_e_*. These conclusions are robust to the choice of DFE parametrization, phylogenetic regression framework, the effect of GC-biased gene conversion, and ancestral misidentification. We also extend the DFE estimation procedure to relax the assumption of additive fitness effects, finding that dominance is only weakly identifiable from the SFS but has minimal impact on comparative DFE inference.

**Summary:** The distribution of fitness effects describes how harmful or beneficial new mutations are and shapes how populations evolve, yet how much it varies between species remains unclear. The authors infer this distribution for protein-coding mutations across 38 primate subspecies from genetic variation. A single underlying distribution, rescaled by each species’ effective population size, accounts for much of the variation: larger populations appear to purge harmful mutations more efficiently and show some evidence of higher adaptive substitution. Relaxing the usual assumption of additive effects, they find that dominance has little impact on the comparative conclusions.

## 1 Introduction

The distribution of fitness effects (DFE) of new mutations captures how genotypes map to fitness and, in turn, how single-step mutations generate the heritable variation on which selection acts. While mere knowledge of the DFE does not allow reconstruction of the topology of the fitness landscape, it is an important emergent quantity as it compresses the molecular and phenotypic contingencies of fitness landscapes into a single probability distribution. This distribution determines the expected genetic load and levels of standing variation, and conditions both the efficacy of natural selection and the rate of adaptation (Gillespie, 1984; Orr, 2002; Bataillon et al., 2022). Although DFEs can, in principle, be measured experimentally, such approaches are limited to a small number of model organisms, and most comparative studies infer them from population genomic data—typically by contrasting the site-frequency spectra (SFS) of putatively *neutral* and *selected* sites to attempt to account for and disentangle the confounding effects of demography and selection (Williamson et al., 2005; Eyre-Walker et al., 2006; Tataru and Bataillon, 2020; Galtier, 2016).

In that context, inferring DFEs provides a powerful window into fitness landscape stability across species. If the underlying fitness landscape is broadly conserved, DFEs for comparable genomic regions should be similar across species. This mapping is necessarily indirect, however, because the DFE reflects both the landscape and a population’s current position on it: the same landscape can yield different DFEs depending on where a population sits (Cherry, 1998; Latrille and Lartillot, 2021), and demographic non-equilibrium can further bias DFE-related inferences (Müller et al., 2022). The converse is nonetheless informative: because populations at different positions on a shared landscape would generally display different DFEs, a DFE that is conserved across many independently evolving species points to a conserved fitness landscape—at least within the region that these populations occupy. Systematic shifts in DFEs across taxa or clades, on the other hand, may indicate changes in constraint, environmental conditions or the fitness landscape itself (Chen et al., 2022b).

Comparing DFEs across species raises a conceptual difficulty, however: the units in which the DFE is defined and estimated. Experimental evolution studies can estimate the distribution of selection coefficients (*s*) directly, provided these are properly defined. In contrast, SFS data are often modeled using a Poisson random field framework (Sawyer and Hartl, 1992; Durrett, 2008), in which sites are treated as independent and SFS bin counts are assumed to follow Poisson distributions around their model expectations. Under this framework, the expected counts in the *neutral* and *selected* SFS are determined by population-scaled mutation rates (*N_e_µ*) and selection coefficients (*N_e_s*). This creates a potential issue (see Bailey and Bataillon, 2016) that is often glossed over in comparative genomics: unscaled selection coefficients (*s*) and population-scaled coefficients (*N_e_s*) are not interchangeable, because the same *s* produces different fixation dynamics and SFS patterns in species with different *N_e_*. SFS–based inference is invaluable for understanding polymorphism patterns within species, but conflating the two quantities in between-species comparisons risks mistaking mere differences in the drift regime for differences in the landscape itself.

To what extent the underlying distribution of intrinsic mutational fitness effects depends on *N_e_* is, in turn, largely governed by epistasis. If fitness effects are site-independent (no epistasis), each mutation carries a fixed unscaled coefficient *s*, so increasing *N_e_* simply rescales *S* = 4*N_e_s* and shifts the population-scaled DFE toward stronger selection. Under epistasis, by contrast, the fitness effect of a mutation depends on the genetic background, which is itself shaped by the selection–drift balance and hence by *N_e_*. Consequently, the underlying unscaled DFE may itself vary with *N_e_*. Depending on the form of epistasis and the structure of the fitness landscape, these background-dependent changes in intrinsic fitness effects may either amplify or offset the direct rescaling effect of *N_e_* on *N_e_s* (Cherry, 1998; Goldstein, 2013; Latrille and Lartillot, 2021). This distinction is particularly salient in primates: earlier comparative work among nine great apes (Castellano et al., 2019) found that the shape of the DFE for deleterious mutations is strikingly similar, yet the strength of purifying selection tracks differences in genetic drift, with claims that positive epistasis and recessive load shape departures from simple *N_e_*-scaling. Testing whether these patterns generalize requires extending the comparison beyond great apes to more distantly related primates.

A second axis of uncertainty concerns beneficial substitutions and the rate of adaptive molecular evolution. This is commonly summarized by the proportion of amino-acid substitutions fixed by positive selection, *α*, and by the corresponding rate of adaptive substitution relative to *neutral* divergence, *ω_a_* = *αω*, where *ω* = *dN/dS* is the nonsynonymous substitution rate relative to the neutral expectation (Eyre-Walker et al., 2006; Galtier, 2016). Across animals, larger *N_e_* is associated with higher estimated *α*, whereas the evidence for *ω_a_*is mixed, with a positive association recovered mainly among closely related species (Galtier, 2016; Rousselle et al., 2020). Primates are a particularly informative setting for this question: they span a roughly 20-fold range of *N_e_* (Kuderna et al., 2023) yet show comparatively low estimated rates of adaptive protein evolution, with *α* estimates for humans often close to zero (Eyre-Walker and Keightley, 2009; Galtier, 2016). For *α* in particular, however, any such *N_e_* trend is difficult to interpret: because *α* = *ω_a_/ω* is a ratio, a larger *N_e_* can inflate *α* simply by purging slightly deleterious mutations more efficiently—shrinking the non-adaptive rate *ω_na_*in its denominator—even when the adaptive rate *ω_a_* itself is unchanged (Galtier, 2016; Moutinho et al., 2020; Rousselle et al., 2020). The adaptive rate *ω_a_* itself is not subject to this denominator effect, but isolating it requires resolving the beneficial component of the DFE, which is only weakly identifiable from the SFS alone and better estimated when between-species divergence is modeled jointly with polymorphism (Tataru et al., 2017). Changes in long-term *N_e_*can likewise bias estimates of *α* (Eyre-Walker and Keightley, 2009).

Dominance adds an additional layer of complexity: most deleterious mutations are partially recessive, because one functional copy usually provides enough protein or enzyme activity for near-normal function, masking much of the effect in heterozygotes (Huber et al., 2018). Theoretical and empirical work further suggests that dominance itself may vary with effect size (Keightley, 1996; Manna et al., 2011; Caballero and Keightley, 1994; Di and Lohmueller, 2024). Yet dominance (*h*) and selection coefficients (*s*) are notoriously hard to infer from the SFS (Huber et al., 2018; Kyriazis and Lohmueller, 2024). The population-genetic signal of dominance is largely driven by selection acting on rare alleles, which are almost always found in heterozygous genotypes. Consequently, the fate of alleles is primarily determined by *N_e_hs* (Kyriazis and Lohmueller, 2024). Current methods for inferring the DFE from SFS data often assume additive fitness effects, making it essential to evaluate how such assumptions affect comparative DFE analyses.

Here, we use fastDFE (Sendrowski and Bataillon, 2024), an SFS–based inference framework, together with a large comparative primate population genomics dataset (Pankratov et al., 2026) to address four questions. First, we quantify the extent to which DFEs are conserved across primates and identify which features are most stable. Second, we ask whether apparent differences in the distribution of *N_e_s* estimated from SFS data can be explained by variation in *N_e_* rather than by changes in *s*, which would instead indicate differences in the underlying fitness landscape. Third, we ask whether the adaptive substitution rate *ω_a_* scales with *N_e_* by contrasting polymorphism-only with divergence-informed fits. Fourth, we develop and test an extension of fastDFE that allows the dominance coefficient to be specified or co-estimated alongside selection coefficients, and evaluate how sensitive cross-species conclusions about DFE conservation are to assumptions about dominance. Throughout, we keep the distinction between *N_e_s* and *s* explicit, which helps separate cross-species patterns driven by differences in the underlying landscape from those driven by drift regime or assumed dominance. In addition, we performed simulations to evaluate the robustness of fastDFE inference under demographic distortions, dominance misspecification, and varying SFS sample sizes (Appendix C).

## 2 Materials and Methods

### 2.1 Data

We used the dataset assembled and curated by Pankratov et al. (2026) comprising 3,240 individuals from 269 species. From this dataset we analyzed a subset of 38 subspecies (1,873 individuals) mapped to reference assemblies with sufficient sample size (*n* = 8) and coding-sequence annotation complete enough to assign site degeneracy. In addition, we included 28 French individuals from the Human Genome Diversity Project (Bergström et al., 2020), as the panel contains no humans. The subset spans Catarrhini—apes (Hominoidea) and Old World monkeys (Cercopithecoidea)—and includes 11 great apes, 14 macaques, 6 baboons, 5 leaf-eating monkeys (Colobinae), and 2 additional cheek-pouch monkeys (Cercopithecinae) outside the baboon and macaque clades. These lineages share a most recent common ancestor approximately 30 million years ago (Figure 1).

**Figure 1:**
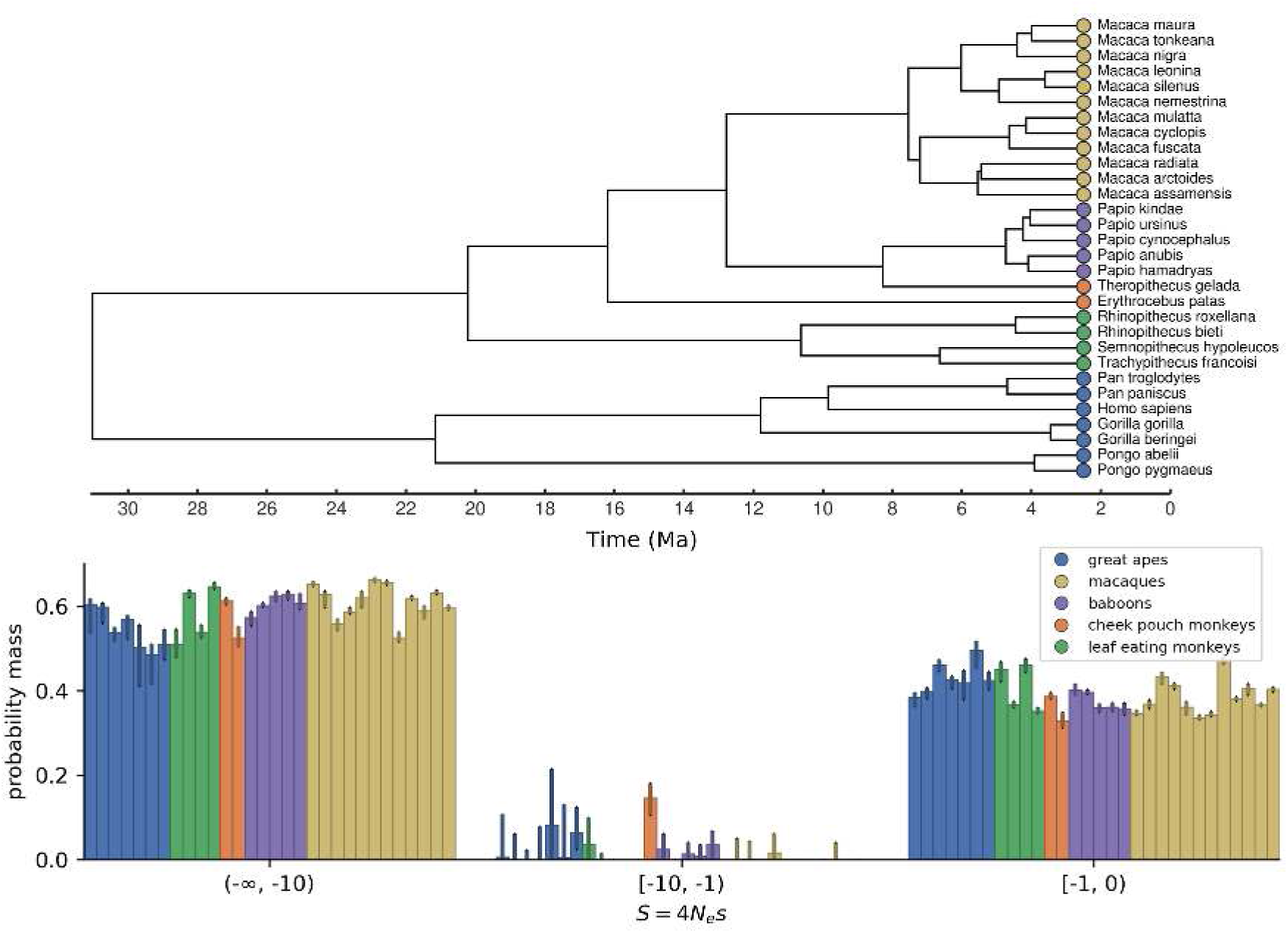
Species-level DFE bar plot and phylogeny. **Top**: Species-level phylogeny. Topology and branch lengths are taken from Kuderna et al. (2023) (supplementary material s4), pruned to the species included in this study. **Bottom**: Discretized DFE for the species in the phylogeny, in the same top-to-bottom order as the tips above. Subspecies-level estimates were averaged per species. DFEs were inferred on unfolded SFS (*n* = 8) using DiscreteParametrization restricted to the deleterious part. Error bars denote 90% confidence intervals, and bar heights correspond to median bootstrap estimates.

Ideally, all species would be mapped to a single reference genome (e.g., the human genome) so that gene annotations and coordinate systems match exactly across species. However, we found that coordinate transfer introduces biases in the SFS for more distantly related species. We therefore retained species-specific reference genomes to preserve within-species accuracy, at the expense of direct comparability across species (see Appendix Table B2). Nevertheless, for ancestral allele inference, outgroup sequences had to be aligned to each ingroup reference. To this end, we generated pairwise whole-genome alignments between ingroup and outgroup reference FASTA files using Minimap2 (Li, 2018), and used these to lift the outgroup reference sequence onto ingroup coordinates, retrieving outgroup alleles at ingroup VCF positions. After generating outgroup-containing VCFs, only biallelic sites were retained. All subsequent steps—degeneracy annotation, ancestral allele inference, SFS construction, and DFE inference—were performed using fastDFE (Sendrowski and Bataillon, 2024).

### 2.2 Degeneracy annotation

DFE inference requires two spectra: one for putatively *neutral* sites and one for *selected* sites. We used fastDFE’s DegeneracyAnnotation utility to classify coding sites as 4-fold degenerate (*neutral*) or 0-fold degenerate (*selected*).

### 2.3 Ancestral allele annotation

Inference of ancestral alleles is required to obtain an unfolded SFS, which records derived allele frequencies relative to their ancestral state. Unfolded spectra are particularly informative about beneficial mutations and are thus necessary for inference of the full DFE, which includes the fraction of beneficial mutations (Schneider et al., 2011; Tataru et al., 2017). Ancestral states for all biallelic SNPs were inferred using fastDFE’s MaximumLikelihoodAncestralAnnotation class, which implements the EST-SFS model (Keightley and Jackson, 2018). Inference proceeds by fitting a fixed phylogenetic tree with branch lengths (shared across sites) and computing, for each site, the probability that each allele is ancestral by comparing the allele states segregating in the ingroup population to the allele state at the homologous position in the outgroup species. Inference was performed separately for each species. Whenever possible, three outgroup species were used, otherwise two (see Appendix Table B1). Sites where more than one outgroup allele was missing were excluded entirely from all downstream analyses. Tree optimization used Kimura’s two-parameter nucleotide substitution model (K2SubstitutionModel), which distinguishes transitions from transversions. The model requires a fixed number of subsampled ingroup haplotypes, set to nine or to the total number available if fewer are present. The method outputs, for each allele, its probability of being ancestral.

### 2.4 SFS parsing

To ensure a sufficiently large sample size for SFS-based inference, we required a minimum of four diploid individuals (eight haplotypes) per subspecies and standardized the SFS sample size to *n* = 8 haplotypes throughout, subsampling species with more individuals to match. This sample size was chosen as a compromise between the number of subspecies retained and the resolution of the SFS (see Figure A1 for the resulting spectra). Both subsampling and polarization were handled probabilistically: subsampling used the expected (hypergeometric-projected) SFS rather than random draws, and each site’s contribution was weighted by its inferred ancestral-allele probabilities rather than fixed to the most likely ancestral state. A further required quantity is the number of mutational target sites, *L*, at which SNPs can potentially occur. We estimated *L* as the per-species coding fraction (computed from the GFF annotation provided with each reference genome) multiplied by the reference genome length, scaled by a factor of 0.8 to account for the exclusion of 2-fold degenerate sites, since only 0-and 4-fold sites were used in the analysis. The factor follows from the standard genetic code: under uniform codon usage, 0-fold and 4-fold sites together account for ∼80% of coding positions. Per-species coding fractions ranged from approximately 1.1% to 1.3% across our reference set. The target-site count *L* directly affects estimates of the effective population size *N_e_*, but influences DFE inference and ancestral allele annotation only weakly, through the expected population-scaled mutation rate used in these models (see below).

Sites in CpG dinucleotide contexts were excluded prior to ancestral allele annotation, as the elevated mutation rate at CpGs violates standard mutation-model assumptions and can bias both ancestral allele annotation and downstream DFE inference. CpG context was determined from the reference sequence, so sites where this context has mutated along the focal lineage may not be recognized as such. The same exclusion was applied when counting target sites, ensuring that *L* counts the same sites used for SNP ascertainment and that estimates of *θ* and downstream quantities are not biased by inconsistent CpG handling. A further potential confounder is GC-biased gene conversion (gBGC), which mimics selection by favoring G/C alleles and can bias DFE inference (Bolívar et al., 2018). As a robustness check, we repeated the analysis with spectra restricted to GC-conservative mutations (A↔T and C↔G), which are immune to gBGC. The same restriction was applied to the target-site count.

### 2.5 Estimating ***N_e_***

The effective population size *N_e_* was estimated as 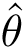/(4*µ*), where *θ* = 4*N_e_µ* is the population-scaled mutation rate and *µ* the mutation rate per site per generation. Species-specific *µ* values were taken from Kuderna et al. (2023), who derived them from fossil-calibrated substitution rates and generation times, so our absolute *N_e_* values inherit the uncertainty of those estimates. 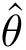 was estimated using Watterson’s estimator (Watterson, 1975), which is unbiased under mutation–drift equilibrium and neutrality. Specifically, 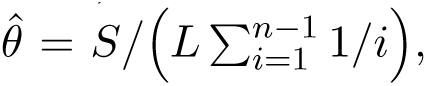 where *S* denotes the number of segregating sites, *n* is the number of subsampled haplotypes, and *L* is the number of mutational target sites. Since this estimator is biased under non-equilibrium demography, we also computed *N_e_* from nucleotide diversity, 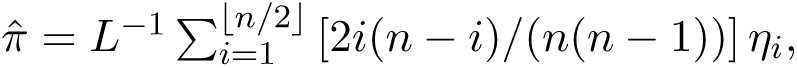 where *η_i_* is the number of sites at which the minor allele is observed *i* times. Both estimators agree closely (Figure A13), and we report Watterson-based estimates throughout. Our *N_e_* estimates are broadly consistent with the coarser, species-level estimates of Kuderna et al. (2023) (Figure A12), while additionally resolving *N_e_* at the subspecies level. Per-subspecies *µ* and *N_e_* values are tabulated in Appendix Table B3.

### 2.6 DFE inference

The DFE was inferred with fastDFE (Sendrowski and Bataillon, 2024) using a *neutral* and a *selected* SFS, where the *neutral* SFS was used to account for demographic and sampling effects via nuisance parameters. Unless stated otherwise, all DFE inference is based on polymorphism alone (the SFS). The estimation that additionally uses divergence counts, which better resolves the beneficial component given the limited sample sizes available here, is described separately (Section 2.7). We performed inference under two alternative DFE parametrizations. The first is a gamma–exponential model (GammaExpParametrization), in which deleterious effects follow a reflected gamma distribution (a gamma distribution over |*S*| mirrored onto the negative axis) and beneficial effects an exponential distribution. Specifically, *S_d_* and *S_b_*denote the population-scaled mean strength of deleterious and beneficial mutations; *b* is the gamma shape parameter; and *p_b_* is the fraction of beneficial mutations. The second parametrization is a discrete DFE (DiscreteParametrization), which assigns weights to predefined intervals of *S* defined by the breakpoints [−∞, −100, −10, −1, 0, 1, ∞]. This approach does not impose a continuous parametric form on the DFE and accommodates species with a large proportion of strongly deleterious mutations, which the gamma–exponential model fits less well. The discrete model has five free parameters, whereas the gamma–exponential model has four. Both can be restricted to model only the deleterious component of the DFE, reducing the number of parameters by two (Appendix Table B4).

We performed 100 local optimization runs per dataset using the default parameter bounds and scales, retaining the best-likelihood run. Convergence was assessed by inspecting the spread of likelihoods across runs, with the top-likelihood plateau reached well before 100 runs in all datasets. Uncertainty was assessed using 100 bootstrap replicates, with 10 optimization runs per bootstrap. SFS bin counts are resampled under a Poisson model, so that the reported confidence intervals quantify uncertainty arising from site-level sampling variation under the fitted model. They therefore do not capture broader sources of uncertainty such as between-gene heterogeneity, biological differences between species, or technical differences in sequencing and variant-calling pipelines. The ancestral misidentification parameter *ε* of fastDFE was fixed to 0, assuming the probabilistic polarization step (see SFS parsing) recovers the ancestral state reliably. Residual misidentification would inflate the high-frequency bins of the unfolded SFS and could bias the inferred beneficial component of the DFE upward. Allowing *ε* to be co-estimated did not improve the fit on our data, so we retained the *ε* = 0 specification. To check the robustness of our findings, we repeated the analyses based on folded spectra, which do not rely on ancestral allele annotation.

From each inferred DFE we computed the predicted nonsynonymous substitution rate relative to neutral, 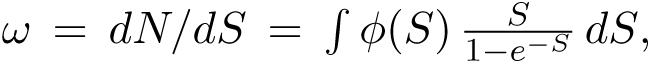 where *ϕ*(*S*) is the DFE density in population-scaled units and *S/*(1 − *e*^−*S*^) is the fixation rate of a mutation with scaled coefficient *S* relative to a neutral mutation. The adaptive component *ω_a_* is defined as the same integral restricted to *S >* 0, the non-adaptive component as *ω_na_* = *ω* − *ω_a_*, and the proportion of adaptive substitutions as *α* = *ω_a_/ω*. All four quantities are functions of the inferred DFE alone and were evaluated for each bootstrap replicate, yielding confidence intervals in the same way as for the other DFE statistics.

### 2.7 Inference from SFS and divergence counts

fastDFE (available from version 1.3.0) was extended to fit the DFE jointly to the within-population polymorphism and the between-lineage divergence. The expected number of fixed substitutions per class is E[*D*_neut_] = *L*_neut_ *r θ* at synonymous sites and E[*D*_sel_] = *L*_sel_ *r θ ω* at nonsynonymous sites, where *L* is the number of mutational target sites, *θ* the population-scaled mutation rate, *ω* the ratio defined in Section 2.6, and *r* = *D*^^^_neut_*/*(*L*_neut_ *θ*) a divergence-scaling factor with *D*^^^_neut_ the observed *neutral* divergence count. The *selected* divergence count *D*^^^_sel_ enters the likelihood as a Poisson term alongside the SFS (Tataru et al., 2017). Allowing *r* to vary as a free parameter, or correcting for polymorphism misattributed as divergence, did not improve recovery of the true DFE in SLiM simulations (not shown), so we retained the model above. The beneficial component of the DFE—and hence *α* and *ω_a_*—is then constrained directly by the observed substitution rate, rather than extrapolated from the SFS alone. Lineage-specific divergence counts were derived per reference genome: at 0-fold (nonsynonymous) and 4-fold (synonymous) coding sites, we counted the fixed differences between each reference genome and the inferred ancestral allele (Section 2.3), reconstructed from outgroup alleles obtained from the pairwise whole-genome alignments (Section 2.1). CpG-context sites were excluded to match the polymorphism SFS, and sites segregating in the focal population were removed so that only fixed differences were retained. The resulting counts were combined with each population’s SFS, and the full DFE was fit with and without divergence (Figure A5). Because divergence is derived from a single reference genome, species mapped to the same reference provide non-independent divergence counts, so the number of independent estimates is set by the number of reference genomes (14) rather than by the number of subspecies (38).

### 2.8 Modeling dominance effects

We extended fastDFE to model and estimate dominance effects (available from version 1.2.0). To this end, we modified the likelihood and numerical integration scheme to incorporate expectations for SFS counts under arbitrary dominance coefficients, based on diffusion approximations (Williamson et al., 2004). In contrast to the semidominant case, for which closed-form expressions exist conditional on fixed selection coefficients, nonadditive effects are modeled in fastDFE by numerical evaluation of a triple integral (over *s*, *h*, and allele frequency). The implementation is numerically stable and was validated against SLiM simulations. Precomputation of integration weights incurs additional upfront cost for non-additive models, but these weights can be reused across likelihood evaluations, mitigating the impact on overall runtime. We further added support for joint inference of the dominance coefficient *h* together with the remaining DFE parameters. This was implemented via a grid over *h* with linear interpolation between grid points. Finally, we extended our framework to support specifying dominance models in which *h* covaries with *S*. This model is motivated by previous theoretical work and empirical studies reporting that slightly deleterious mutations tend to be nearly additive while lethal ones are often very recessive (Keightley, 1996; Manna et al., 2011; Huber et al., 2018; Kyriazis and Lohmueller, 2024). The functional form can be user-defined via a callback, and any hyperparameters it introduces are treated as parameters that may be fixed or optimized. These extensions are described in the fastDFE documentation.

### 2.9 Regression analysis

Associations between DFE properties and log_10_ *N_e_* were tested using ordinary least squares (OLS) and phylogenetic generalized least squares (PGLS). Models were of the form (*x* ∼ log_10_ *N_e_*), where *x* stands for a property of the DFE such as the shape parameter of the gamma distribution used for modeling deleterious mutations, or the proportion of mutations with *S* ∈ (−1, 0]. Models were fitted using the ape (Paradis and Schliep, 2019) and nlme (Pinheiro and Bates, 2000) R packages. Because the phylogeny is available only at species resolution, subspecies-level estimates of DFE properties were averaged per species prior to regression. Species not present in the phylogeny (*Macaca leucogenys*, *M. hecki*, and *Trachypithecus poliocephalus*) were excluded, leaving 30 species for analysis. Phylogenetic covariance was modeled under Brownian motion and Grafen’s branch-length transformation, using the species tree from Kuderna et al. (2023) (supplementary material s4). Throughout the main text and figures, reported *r* and *p* values come from the same Grafen PGLS fit on the species-averaged data: *p* is the two-sided *p*-value of the slope of log_10_ *N_e_*, and 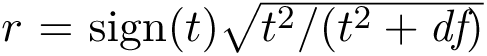 is derived from the slope’s t-statistic and residual degrees of freedom *df*, using the identity 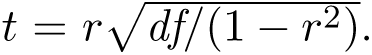 This makes *r* a phylogenetically corrected analogue of Pearson’s correlation. Full OLS, PGLS Brownian, and PGLS Grafen regression tables for the strongly deleterious fraction (*S* ∈ (−∞, −10]) across the four DFE-model variants (including generation time as a second covariate) are provided in Appendix Tables B6–B9.

To test whether the unscaled DFE for deleterious mutations itself varies with *N_e_* (residual epistasis) without relying on the parametrization-sensitive mean *S_d_*, we used a quantile diagnostic. For each population we computed the population-scaled coefficient *S_q_* below which a fixed fraction *f* of the deleterious probability mass lies, and regressed log_10_ |*S_q_*| on log_10_ *N_e_* as above. Because a quantile fixes a cumulative-probability level rather than a point on the effect-size axis, it samples a comparable, observable position in every species and is insensitive to the effectively lethal tail, which the SFS cannot constrain because very strongly deleterious mutations rarely segregate. Under no epistasis it rescales exactly as *S_q_* = 4*N_e_s_q_*, so the expected slope is 1. We verified this calibration on a simulated *N_e_*-rescaled dataset (a fixed unscaled DFE scaled by 4*N_e_*; Figure A15), on which the quantile slope is exactly 1 at every *f*. The *p*-value for the no-epistasis null hypothesis (*H*_0_: slope = 1) was obtained by regressing the unscaled quantile log_10_ |*s_q_*| = log_10_ |*S_q_*|−log_10_(4*N_e_*) on log_10_ *N_e_*, for which the no-epistasis null hypothesis corresponds to a slope of zero, yielding an identical test.

## 3 Results

### 3.1 The DFE across catarrhines

Across the catarrhine phylogeny (Figure 1), great apes exhibit less deleterious population-scaled DFEs than other catarrhines, consistent with their lower effective population sizes. Estimates are based on unfolded SFS (*n* = 8) using DiscreteParametrization restricted to the deleterious part, and DFEs are reported in units of the population-scaled selection coefficient *S* = 4*N_e_s*. Figure A17 shows the same estimates at subspecies resolution, ordered by *N_e_* rather than by phylogenetic relatedness. The estimates appear to cluster by clade, which may reflect a genuine clade-specific effect or simply the tendency of closely related taxa to share similar values of *N_e_*.

To distinguish these possibilities, we plot the mass of the inferred DFE across effect-size intervals directly against *N_e_*, now at subspecies resolution (Figure 2). Under both the DiscreteParametrization and the GammaExpParametrization, *N_e_* correlates positively with the fraction of strongly deleterious mutations *S* ∈ (−∞, −10] (discrete: *r* = 0.82, *p* = 3.6 × 10^−8^; gamma: *r* = 0.83, *p* = 1.6 × 10^−8^) and negatively with the fraction of effectively neutral mutations *S* ∈ (−1, 0] (discrete: *r* = −0.72, *p* = 8.1 × 10^−6^; gamma: *r* = −0.70, *p* = 1.4 × 10^−5^). For the moderately deleterious fraction *S* ∈ (−10, −1], neither parametrization shows a significant trend (discrete: *r* = −0.28, *p* = 0.14; gamma: *r* = −0.30, *p* = 0.11). This is expected, because such mutations segregate at frequencies similar to those of nearly neutral mutations, so the data separate the two classes poorly.

**Figure 2:**
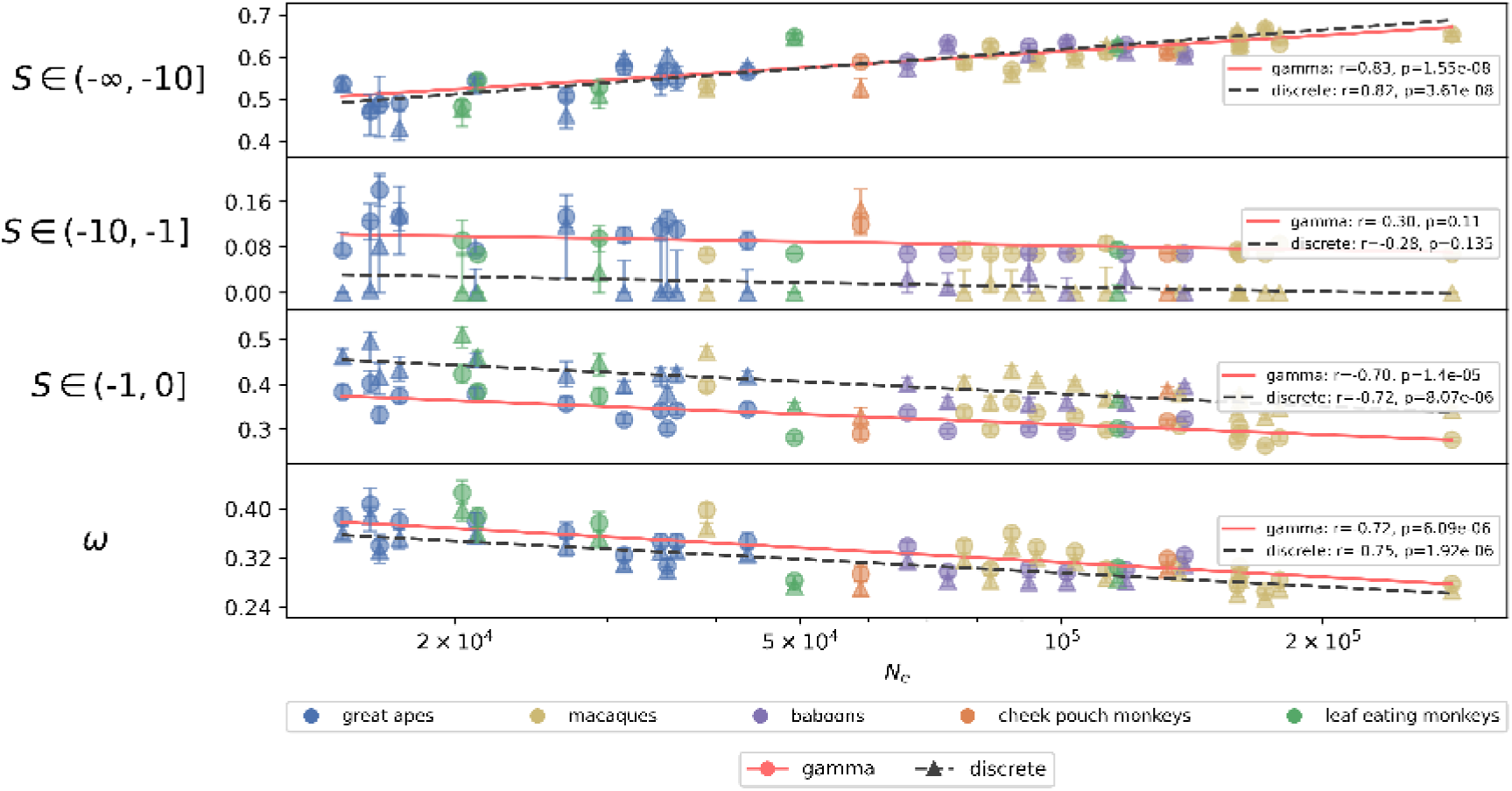
Comparison of the DFE for deleterious mutations across Catarrhini, under both DFE parametrizations. Each point represents a species or subspecies, plotted against its effective population size (*N_e_*); the two colors overlay the deleterious-only DiscreteParametrization and GammaExpParametrization, inferred on unfolded SFS (*n* = 8). The top three subplots show the fraction of mutations in successive intervals of *S*; the bottom panel shows the predicted total nonsynonymous substitution rate *ω* = *dN/dS* relative to the neutral synonymous rate, computed directly from the inferred DFE (from polymorphism alone). Error bars denote 90% confidence intervals, and points correspond to median bootstrap estimates. Dashed lines indicate Grafen PGLS regression fits on species-averaged data; reported *p* and *r* values are the slope *p*-value and a phylogenetically corrected correlation coefficient derived from the slope’s t-statistic (see Methods).

These associations, all inferred from polymorphism-only data, are robust to the choice of DFE parametrization and to whether the SFS is folded: the same qualitative trends appear when the deleterious-only DFE is fit on folded spectra (Figure A4) and when the full DFE (deleterious plus beneficial) is fit under either parametrization (Figure A5).

### 3.2 Model support

Of the four DFE-model variants we considered, DiscreteParametrization restricted to the deleterious part had the most species-level support: it gave the lowest AIC for 14 of 38 subspecies (more than any other variant) and a likelihood-ratio test against the full version was non-significant for most species (Appendix Table B5). The gamma counterpart, by contrast, fits poorly for species with a sizable fraction of mutations under strong purifying selection, because it cannot place substantial probability mass in the strongly deleterious (*S* ∈ (−∞, −10]) and effectively neutral (*S* ∈ (−1, 0]) categories simultaneously. The deleterious-only GammaExpParametrization therefore either saturates at the parameter bound |*S_d_*| = 10^5^ for many large-*N_e_* species, or fits the heavy tail at the cost of the effectively neutral bins. The discrete parametrization does not face this constraint, as it assigns probability mass to each *S* interval independently, including separate effectively neutral deleterious and effectively neutral beneficial classes.

### 3.3 Can variation in ***N_e_*** explain the observed differences?

Because the DFE is reported in units of *S* = 4*N_e_s*, it necessarily depends on *N_e_*. The relative fractions of mutations assigned to each DFE interval are, however, determined by the shape of the SFS and the proportion of *neutral* to *selected* polymorphism, not by the total number of segregating sites: the inference is invariant to uniform rescaling of the *neutral* and *selected* SFS counts (Figure A16), and so does not depend on the absolute level of *θ*. This rules out trivial artifacts driven by larger-*N_e_* species simply contributing more polymorphism, but does not by itself establish invariance to biological variation in *N_e_*, which can additionally reshape the SFS through differences in the efficacy of selection. Instead, the observed negative association between *N_e_* and the fraction of effectively neutral mutations (*S* ∈ (−1, 0]; Figure 2) is consistent with the nearly neutral theory (Ohta, 1992). The corresponding positive correlation between *N_e_* and the fraction of mutations in the most deleterious category (−∞, −10] follows compositionally: a lower fraction of effectively neutral mutations necessarily redistributes mass into the more strongly deleterious bins. More directly, simulating a fixed unscaled DFE and rescaling it by 4*N_e_* across the empirical *N_e_* range reproduces the same bin-vs-*N_e_* trends observed in our scaled DFEs (Figure A15), consistent with the interpretation that the scaled-DFE shifts we report are driven by variation in *N_e_*.

To assess whether intrinsic fitness effects differ across species, we converted the scaled DFEs to unscaled selection coefficients *s*by dividing by 4*N_e_*. The resulting unscaled DFEs are largely similar across species, with comparable probability mass across intervals of *s* (Figure A8). This is a graphical check only. To test for residual structure formally, we regressed the tail-robust quantile *S_q_* of the DFE for deleterious mutations on *N_e_* (Methods). Unlike the mean |*S_d_*|, this quantile is insensitive to the effectively lethal tail that the SFS cannot constrain, and under no epistasis it rescales as *S_q_*= 4*N_e_s_q_*, giving an expected slope of 1 for log_10_ |*S_q_*| on log_10_ *N_e_*. Across three quantile levels (*f* = 0.4, 0.5, 0.6) and all four DFE-model variants, the estimated slope is statistically indistinguishable from the no-epistasis value of 1 (all *p >* 0.05; for example the deleterious-only GammaExpParametrization gives a slope of 1.20, *p* = 0.20, at *f* = 0.6), and the estimated slopes scatter around 1 with deviations of no consistent sign (Figure A9; Table 1). We therefore find no evidence for epistasis on the DFE for deleterious mutations: at the resolution of these data the unscaled DFE is consistent with being conserved across the ∼20-fold range of *N_e_* spanned by catarrhines.

**Table 1:** Tail-robust quantile test for epistasis on the DFE for deleterious mutations. PGLS slope of the regression of log_10_ |*S_q_*| on log_10_ *N_e_* across catarrhine species (unfolded SFS, *n* = 8), with the phylogenetically corrected correlation *r* and the *p*-value for the no-epistasis null hypothesis *H*_0_: slope = 1, across the four DFE-model variants (gamma/discrete × del/full) and three quantile levels *f*. Here *S_q_* is the population-scaled coefficient below which a fraction *f* of the deleterious mass lies. Under no epistasis a fixed quantile rescales as *S_q_* = 4*N_e_s_q_*, so the expected slope is 1. Across all quantile levels and parametrizations the slope is statistically indistinguishable from 1 (all *p >* 0.05), with deviations of no consistent sign, indicating no detectable epistasis. The discrete parametrization has a step CDF and hence coarse quantiles.

| $f$ | | gamma del | gamma full | discrete del | discrete full |
| --- | --- | --- | --- | --- | --- |
| 0.4 | slope | 1.13 | 0.17 | 0.14 | 0.40 |
| | $r$ | 0.09 | -0.26 | -0.33 | -0.18 |
| | $p$ | 0.63 | 0.16 | 0.08 | 0.34 |
| 0.5 | slope | 1.30 | 0.62 | 0.61 | 0.12 |
| | $r$ | 0.29 | -0.11 | -0.10 | -0.22 |
| | $p$ | 0.12 | 0.56 | 0.61 | 0.24 |
| 0.6 | slope | 1.20 | 1.11 | 2.14 | 1.69 |
| | $r$ | 0.24 | 0.03 | 0.34 | 0.17 |
| | $p$ | 0.20 | 0.89 | 0.07 | 0.38 |

We also compared the DFE-based results with the relationship between *N_e_* and *π_N_ /π_S_*across species. Here, *π_N_* and *π_S_* denote pairwise nucleotide diversity at nonsynonymous and synonymous sites, respectively, so their ratio provides a model-free proxy for the strength of purifying selection acting in a population (James et al., 2017). Because larger *N_e_*increases the efficacy of selection against mildly deleterious mutations, we expect an inverse relationship between *N_e_* and *π_N_ /π_S_* (Welch et al., 2008; Bastian et al., 2026). Under the nearly neutral theory with a gamma-distributed DFE for deleterious mutations with shape parameter *b*, this relationship is expected to take the power-law form *π_N_*/*π_S_* ∝ *N_e_^-b^*, so that a regression of log_10_(*π_N_ /π_S_*) on log_10_ *N_e_* has slope −*b* (Welch et al., 2008; James et al., 2017). Consistent with this expectation, we observe a strong negative association (Figure A14). In addition, *π_N_ /π_S_* exhibits clear taxonomic clustering across primate clades, similar to that observed for the DFE estimates. To assess how much information about the inferred DFE for deleterious mutations is already captured by *π_N_ /π_S_*, we refitted the DFE with an SFS sample size of two (Figure A7), which relies only on the total and polymorphic site counts in a pairwise sample and so uses the same information as *π_N_ /π_S_*. The qualitative patterns match those obtained at larger sample sizes, with correlations in the same direction but weaker and substantially more uncertain, indicating that this ratio captures part of that information, consistent with the findings of Chen et al. (2022a).

### 3.4 Substitution rates and adaptive evolution

Under both parametrizations restricted to the deleterious part, the total nonsynonymous substitution rate *ω* = *dN/dS* declines significantly with *N_e_* (discrete: *r* = −0.75, *p* = 1.9 × 10^−6^; gamma: *r* = −0.72, *p* = 6.1 × 10^−6^), as expected under the nearly neutral theory (bottom panel of Figure 2). The same decline is recovered when fitting the gamma deleterious-only model on the folded SFS (*r* = −0.89, *p* = 5.6 ×10^−11^; Figure A4). Adding a free beneficial component (the full DFE), still fit to polymorphism alone, gives a more nuanced picture. Under the decomposition *ω* = *ω_a_* +*ω_na_*, the non-adaptive component *ω_na_*keeps declining with *N_e_*, as expected from more efficient purging. The total *ω*, however, is flatter, because the fits now place some mass on beneficial substitutions (*ω_a_ >* 0), and its decline with *N_e_* is mostly insignificant under the full models (Figure A5). However, neither *ω_a_* nor the adaptive fraction *α* = *ω_a_/ω* shows a significant association with *N_e_*. For *ω_a_* we obtain *r* = 0.16, *p* = 0.41 under GammaExpParametrization and *r* = −0.12, *p* = 0.52 under DiscreteParametrization. For *α* the corresponding values are *r* = 0.34, *p* = 0.06 and *r* = 0.19, *p* = 0.31. This lack of signal is expected: at the small sample size used here (*n* = 8), slightly beneficial and slightly deleterious mutations are difficult to distinguish from polymorphism alone, leaving *ω_a_* and *α*weakly identifiable. Including divergence counts in the fit (Methods) leaves the *N_e_* dependence of the deleterious bins and of *ω_na_*unchanged. The key difference is that *ω_a_* now increases significantly with *N_e_* under either parametrization (gamma: *r* = 0.57; discrete: *r* = 0.64; both *p <* 2 × 10^−3^, Figure A5). Because *ω_a_* itself rises—not only the ratio *α*—this cannot arise from the shrinking *ω_na_* denominator alone, and these correlations remain significant despite the reduced power of only 14 independent reference lineages under phylogenetic regression. The divergence-informed signal nonetheless warrants caution, because using divergence to resolve the beneficial component assumes a shared DFE between segregating polymorphism and fixed substitutions (Tataru et al., 2017). Those substitutions accumulated over deep time, along the lineage back to an ancestor inferred from the outgroups, so they reflect the historical *N_e_*, whereas we regress against the contemporary *N_e_* estimated from each population’s SFS. The trend is therefore sensitive to outgroup choice and to past changes in *N_e_* (Eyre-Walker and Keightley, 2009; Rousselle et al., 2018).

### 3.5 Comparison with previous estimates

We compared our results with those of Castellano et al. (2019) by restricting the analysis to the same nine great ape populations. The individuals analyzed by Castellano et al. (2019), drawn from the dataset of Prado-Martinez et al. (2013), form a subset of those used in the present study. In contrast to our approach, their reads were mapped to the human reference genome (Figure A2). As in the original study, we used GammaExpParametrization on unfolded SFS (*n* = 8) and shared the shape parameter *b* across species when fitting the DFE. The results are nearly identical: for example, for *S* ∈ (−∞, −10] and *N_e_*, we have (*r* = 0.89, *p* = 1.68 × 10^−2^) in the new dataset versus (*r* = 0.89, *p* = 1.82 × 10^−2^) in the original analysis (Figure A3). Castellano et al. (2019) also reported a positive correlation between the unscaled mean deleterious selection coefficient *s_d_* and *N_e_*. We recover the same qualitative trend, but in our analysis it is significant in neither the old (*r* = 0.75, *p* = 8.27 × 10^−2^) nor the new dataset (*r* = 0.44, *p* = 3.87 × 10^−1^) and becomes weakly negative and non-significant when DFEs are inferred independently using DiscreteParametrization. Finally, we do not observe the outlier behavior of bonobos and western chimpanzees reported by Castellano et al. (2019). Instead, these populations conform to the same overall trend as the other great apes. Despite the overall agreement between the two analyses (Figure A3), the remaining differences from their reported results may stem from the DFE inference framework, the derivation of the SFS, the regression framework, or the *N_e_* estimates.

### 3.6 GC-biased gene conversion

To assess the influence of gBGC, we additionally constructed the SFS from GC-conservative polymorphisms alone and examined whether this altered the inferred DFE or its covariation with *N_e_*. The resulting DFEs are slightly less deleterious under the deleterious-only model (Figure A10) and marginally so under the full DFE, suggesting that gBGC inflates the apparent strength of purifying selection. This trend is qualitatively consistent with the SFS estimates of Castellano et al. (2019), but more pronounced in our data. In their analysis, the shift did not translate into a consistent difference in inferred DFE parameters. The qualitative association between *N_e_* and the DFE is preserved. Beyond its effect on the DFE for deleterious mutations, recombination-associated gBGC can also generate apparent positive selection without a corresponding evolutionary advantage (Joseph, 2024). Restricting both polymorphism and divergence counts to GC-conservative mutations preserves the divergence-informed *ω_a_*–*N_e_* and *α*–*N_e_* associations under the discrete parametrization (*ω_a_*: *r* = 0.65, *p* = 1.1 × 10^−4^; *α*: *r* = 0.54, *p* = 2.3 × 10^−3^). However, the gamma–exponential fit is uninformative here, since the gamma form cannot simultaneously accommodate substantial strongly deleterious and effectively neutral mass, so that *S_d_*collapses to its bound for 17 of the 38 subspecies (Figure A6).

### 3.7 Gene-set differences

Because each species is analyzed against its own reference genome, the annotated gene sets differ between species. To assess whether this affects our results, we recomputed the spectra with sites restricted to the 14,437 ortholog groups present in all 14 references, defined by one-to-one orthology with human genes in the NCBI gene orthology database, and reinferred the DFE. These groups cover ∼75% of coding sequence per reference (72– 80%). The restricted set yields a more deleterious DFE, possibly because these genes are more conserved, but the associations with *N_e_* are preserved (Figure A11).

### 3.8 Dominance

Lastly, we examined how assuming that mutations have partially recessive (rather than additive) fitness effects changes the inferred DFE, taking the dominance–selection-strength relationship to be inverse, as supported by empirical studies (Huber et al., 2018; Kyriazis and Lohmueller, 2024). We use the representative functional form *h*(*S*) = 0.4 exp(−0.02|*S*|), such that *h* is ∼0.4 for weakly selected mutations and decays toward zero for strongly deleterious ones (specifically, *h*(−100) ≈ 0.05). Empirical estimates of this relationship remain highly uncertain, and the analysis here is intended as a sensitivity check rather than a definitive characterization of the (*h, S*) relationship. The inferred DFEs show only modest changes, consistent across species (Figure 3). The shift toward more deleterious mutations follows from masking in heterozygotes: to reproduce the same SFS under reduced dominance, the underlying homozygous selection coefficient (*S*) must be more negative.

**Figure 3:**
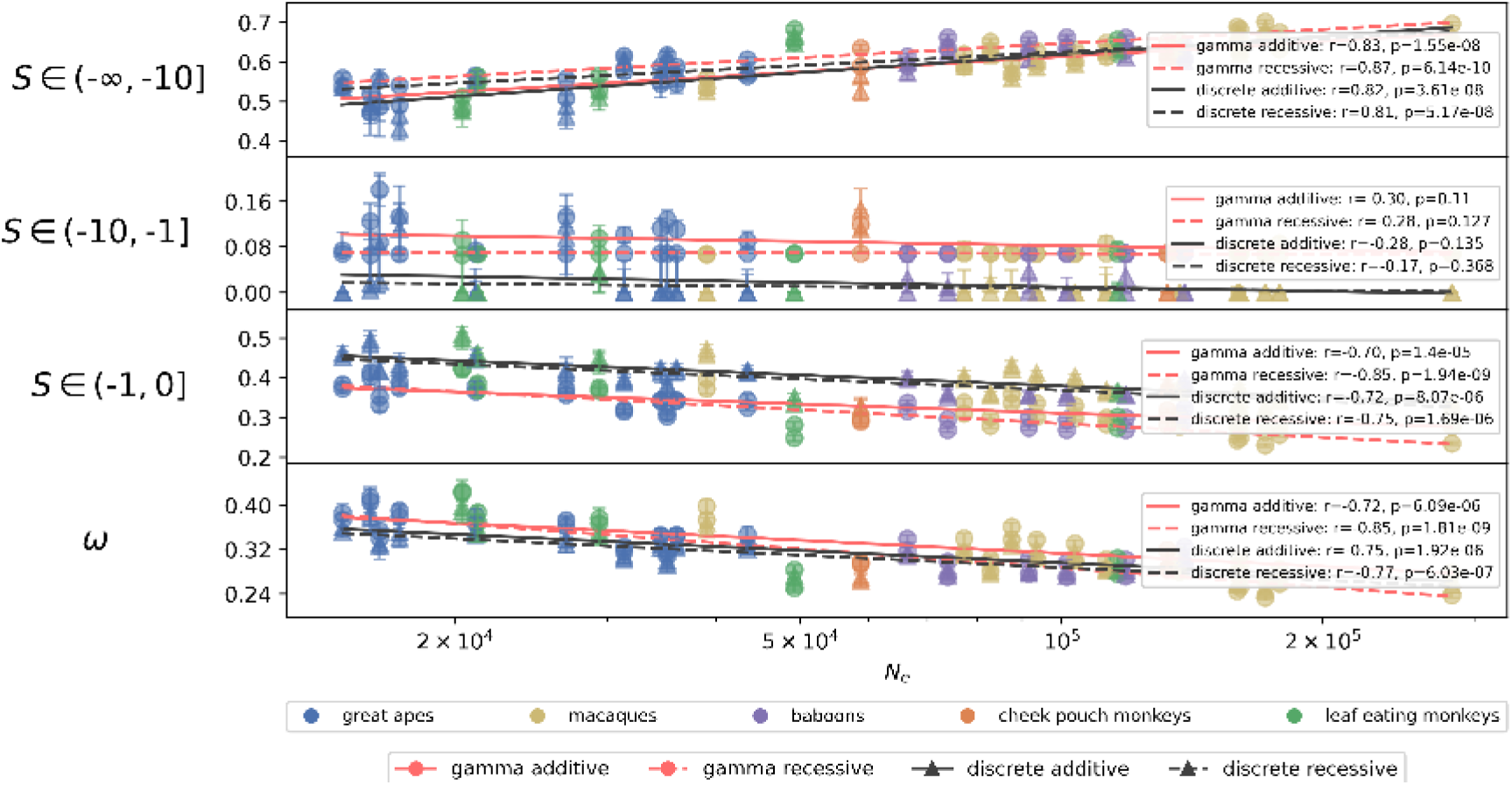
Comparison of the DFE for deleterious mutations under fully additive versus partially recessive mutations. Regression of DFE properties on *N_e_*, inferred from the unfolded SFS (*n* = 8) under both the GammaExpParametrization (red, circles) and DiscreteParametrization (black, triangles); within each, additivity (*h* = 0.5) is shown with solid lines and recessivity with dashed lines. Dominance is modeled as *h*(*S*) = 0.4 exp(−0.02|*S*|), yielding near-additivity (∼0.4) for weakly selected mutations and increasing recessivity with larger |*S*|. The bottom panel shows the predicted total substitution rate *ω* = *dN/dS*, which declines with *N_e_* under both dominance assumptions and both parametrizations, with only small offsets between them.

## 4 Discussion

Across 38 catarrhine subspecies, we find *N_e_* to be the dominant axis of cross-species variation in the DFE for deleterious mutations: the population-scaled DFE shifts systematically with *N_e_*, while the underlying unscaled DFE remains largely conserved. Intrinsic differences in the selective landscape of amino-acid–changing mutations, if any, therefore lie below the resolution of current SFS-based inference at the catarrhine phylogenetic scale, and cross-species DFE differences we observe in scaled units primarily reflect variation in the efficacy of selection rather than in the landscape itself. Previous work in plants has found DFEs to be similar within species across populations, with between-species variation more strongly associated with life-history traits than with demography (Chen et al., 2022b; James et al., 2023). At broader phylogenetic scales, recent comparative work across diverse animal taxa has reported a negative correlation between *N_e_* and the mean unscaled selection coefficient *s* (Lin et al., 2025), suggesting that some properties of the fitness landscape itself may vary across deeper phylogenetic distances or contrasting life-history strategies. Within primates, by contrast, the absence of comparable intrinsic-landscape differences in our analysis may reflect the close phylogenetic relatedness and similar genome architecture of the species analyzed here, including their gene content and recombination landscapes, as well as potential ascertainment bias from focusing on mutations in exons rather than more dynamic genomic regions.

Beyond these empirical comparisons, this result can be placed within existing theory on how the DFE depends on *N_e_*. Models of equilibrium evolution on a fixed landscape with substantial epistasis predict that the population-scaled DFE should be approximately invariant with respect to *N_e_*, because larger populations occupy higher-fitness states where individual mutations have smaller |*s*|, compensating for the greater *N_e_* in the product *N_e_s* (Cherry, 1998; Goldstein, 2013). This is not what we observe. Our results are instead more consistent with site-independent (Ohta, 1992) or weak-epistasis (Latrille and Lartillot, 2021) regimes, in which a largely fixed unscaled DFE combined with variation in *N_e_* shifts mass between population-scaled bins. A consistent reduction in the nearly neutral fraction with *N_e_* was recently reported across 28 mammalian populations by Latrille et al. (2024) (*r*^2^ = 0.31, *p* = 0.001), and the classical nearly neutral signature—a negative *π_N_ /π_S_*–*π_S_* slope—has been documented widely, such as across mammalian mitochondrial genomes (James et al., 2017) and at the nuclear-genome scale across placental mammals (Bastian et al., 2026). However, the DFE alone cannot distinguish a conserved landscape probed at a conserved position from more complex scenarios, and our results are necessarily restricted to the region of the landscape sampled by these populations.

Some cross-species variation remains beyond what *N_e_* explains. This is expected, however, as our bootstrap confidence intervals capture only site-level Poisson sampling and can underestimate the total inference uncertainty. Castellano et al. (2019) reported part of this residual variation across great apes and interpreted it as positive epistasis, whereby new deleterious mutations have smaller effects in already-damaged genetic backgrounds. Our data do not support this interpretation. Their inference rested on the mean deleterious coefficient *S_d_*, a statistic we consider unsuitable for this purpose. Because a substantial fraction of new nonsynonymous mutations are effectively lethal, *S_d_*is dominated by the left tail of the DFE that the SFS cannot constrain, and is thus parametrization-dependent: discrete models place explicit mass at very large |*S*|, inflating *S_d_*, whereas gamma models truncate or smooth that tail. The *N_e_*-slope of *S_d_* therefore tracks the assumed tail shape rather than the data, and cross-parametrization comparisons of *S_d_* are not meaningful. For this reason we largely refrain from reporting *S_d_* except for comparison with previous results, and instead favor interval probabilities (e.g., *P* [*S <* −10]).

Testing the unscaled DFE against *N_e_* directly is likewise problematic, because the same *N_e_* enters both as the predictor and in the denominator of *s* = *S/*(4*N_e_*), so that estimation error in *N_e_* can introduce a spurious *s*–*N_e_* correlation even when the unscaled DFE is invariant. A tail-robust quantile diagnostic avoids these problems by fixing a cumulative-probability level that is insensitive to the unconstrained tail (Methods). Under this diagnostic, the slope of log_10_ |*S_q_*| on log_10_ *N_e_* is statistically indistinguishable from the no-epistasis value of 1 across all parametrizations and quantile levels, so the unscaled DFE for deleterious mutations is consistent with being conserved across *N_e_* at this phylogenetic scale.

Several methodological choices warrant comment. Our *N_e_* estimates (Methods) may be affected by SNP-calling differences across species and by any residual non-neutrality of synonymous sites that would bias 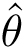. fastDFE also assumes site independence and does not explicitly model linked selection. Its nuisance-parameter machinery should nonetheless absorb some of the distortion that background selection imposes on the *neutral* SFS, since 0-fold and 4-fold sites are physically interspersed within exons and are therefore affected similarly. Forward simulations in Appendix C show that at a human-like recombination rate (*r* = 10^−8^) the simulated SFS departs only slightly from the expectation under site independence. This safeguard is not complete, however—background selection need not distort 0-fold and 4-fold sites equivalently, and recombination-rate heterogeneity can alter DFE inferences (Johri et al., 2020, 2021, 2023; Soni et al., 2025)—so we cannot rule out that residual linked-selection effects contribute to the cross-species patterns we report.

Regardless of these caveats, DFE properties regressed against log_10_ *N_e_* were stable across regression frameworks (OLS, PGLS–Brownian, PGLS–Grafen) and DFE parametrizations. We report *p*-values under PGLS–Grafen, which estimates the degree of phylogenetic correction from the data rather than fixing it a priori (Methods; Appendix Tables B6–B9). The population-scaled DFE for deleterious mutations also carries phylogenetic signal: for the strongly deleterious fraction (*S* ∈ (−∞, −10]), Pagel’s *λ* lies between 0.45 and 0.56, significantly above zero (no phylogenetic structure) but below one (Brownian motion). Closely related species therefore share similar population-scaled DFEs, as expected given that they also share similar *N_e_*. Finally, adding generation time (Kuderna et al., 2023) as a second predictor did not improve fit, indicating that the residual cross-species variation in the DFE for deleterious mutations is not explained by generation time beyond what *N_e_*already captures.

Our analysis of adaptive evolution based on polymorphism alone finds that neither the adaptive fraction *α* nor the adaptive substitution rate *ω_a_* is significantly associated with *N_e_*under any of the parametrizations or SFS variants we considered, although the point estimates are mostly positive. More broadly, the beneficial component is intrinsically difficult to estimate from polymorphism: beneficial mutations are rare and leave only weak signatures in the SFS, and at *n* = 8 haplotypes slightly beneficial mutations cannot be reliably distinguished from slightly deleterious ones, leaving *p_b_* highly variable. Simulations indicate that larger samples would substantially improve power when fitting the full DFE (Appendix C). Jointly modeling lineage-specific divergence is more informative. There, both *α* and *ω_a_* increase significantly with *N_e_* under either parametrization and are robust to phylogenetic correction and gBGC (Figures A5, A6), consistent with a higher rate of adaptive substitution in larger-*N_e_* species. However, we interpret this cautiously, as these substitutions accumulated over deep time and so reflect the historical rather than the contemporary *N_e_*.

Turning to dominance, we find that inferring DFEs under the assumption of additive fitness effects when mutations are in fact partly recessive shifts the inferred DFE toward less strongly deleterious effects, because masking in heterozygotes is not accounted for. This shift is modest and consistent across species, leaving cross-species comparisons and *N_e_*– DFE associations qualitatively unchanged. Simulations further show that *h*is only weakly identifiable from SFS data, because it covaries strongly with the strength of selection, making reliable inference unrealistic in most current empirical settings (Appendix C). Nonetheless, the extension of fastDFE to specify or jointly estimate the dominance coefficient *h*, including models in which *h* covaries with *S*, constitutes a useful addition that users may wish to explore further. Correction for demographic distortions using nuisance parameters also performs well across a wide range of non-equilibrium scenarios (Appendix C).

Taken together, our results are consistent with a regime in which, at the catarrhine phylogenetic scale we examined, the scaled DFE for deleterious mutations shifts primarily as a passive consequence of *N_e_* variation acting on a largely conserved unscaled landscape, rather than as evidence for clade-specific changes in selection pressures or in the underlying genotype-to-fitness map. Whether this picture holds at deeper phylogenetic scales, or across genomic compartments evolving under different constraints, remains an open question. Future work would benefit from incorporating genomic covariates (Yıldırım and James, 2026) to better disentangle how much variation in the DFE occurs within genomes versus across species and to provide finer-grained resolution for detecting local changes in the DFE across species within specific genomic contexts.

## Data Availability

The population genomic data analyzed in this study were generated by Pankratov et al. (2026), except for the human data, which are from the Human Genome Diversity Project (Bergström et al., 2020). Reference genome assemblies and their GenBank accessions are listed in Appendix Table B2, and per-species mutation rates and generation times are from Kuderna et al. (2023). The analysis workflow, derived site-frequency spectra, and results underlying this study are available at https://github.com/Sendrowski/PrimateDFE.

## Acknowledgments

ChatGPT (GPT-4o and GPT-5 series) and Claude (Opus series) were used to assist with code development and to refine the phrasing of the manuscript.

## Funding

This work has been supported by the Novo Nordisk Foundation (Data Science Collaborative Research Programme, grant 0069105).

## Author Contributions

T.B. and J.S. designed the study and carried out the regression analysis, and T.B. supervised the project. J.S. carried out the main analyses and wrote the manuscript, with T.B. contributing to the Introduction. B.M.P. contributed to the analyses and, together with J.B. and V.P., assembled and curated the underlying dataset. T.B. provided detailed feedback on the manuscript, and all authors commented on and approved the final version.

## Competing Interests

The authors declare no competing interests.

## A Supplementary Figures

**Figure A1:**
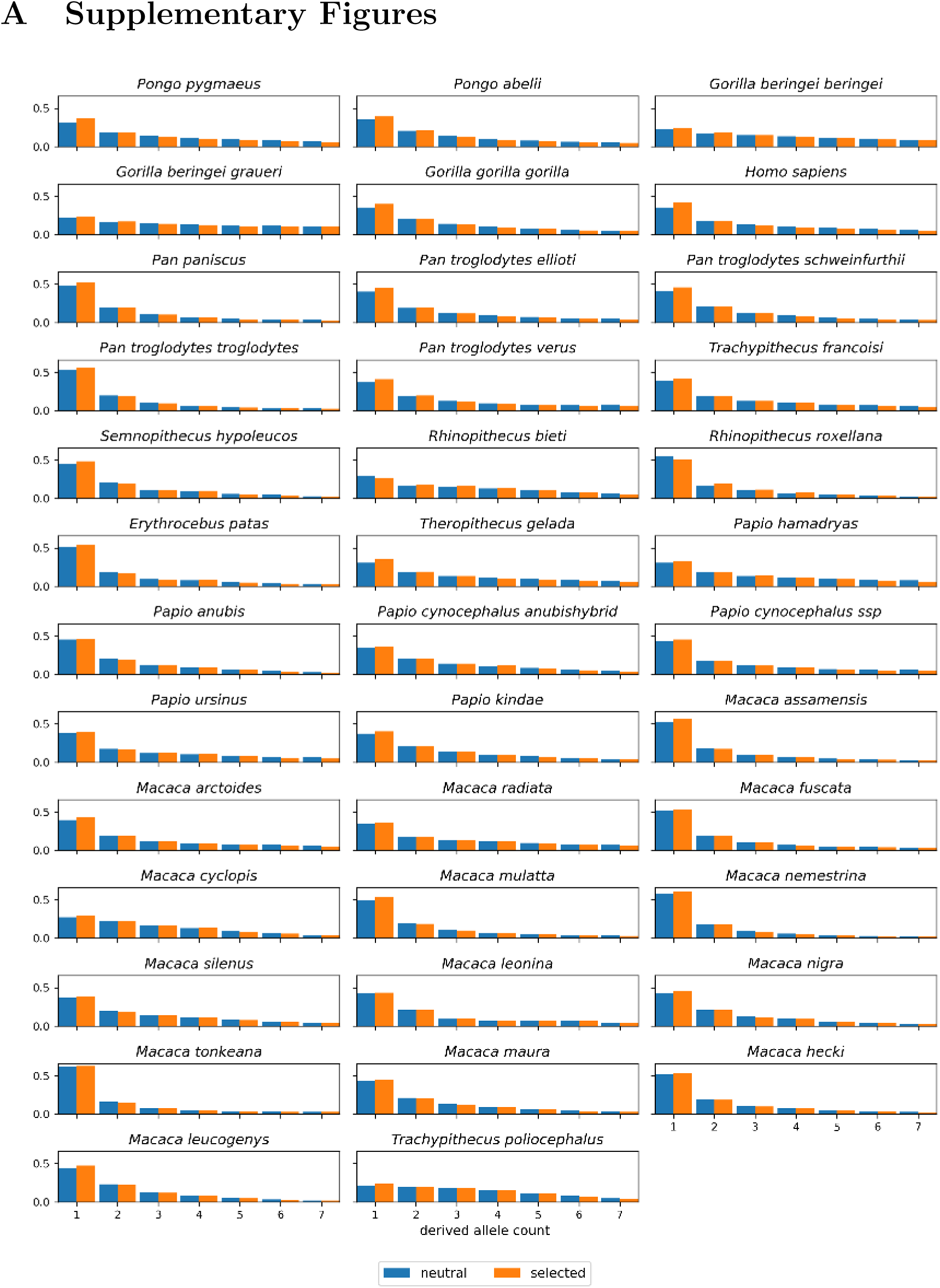
Observed spectra. *Neutral* (4-fold) and *selected* (0-fold) SFS for each of the 38 subspecies, from the unfolded spectra used for inference (*n* = 8). Each spectrum is normalized to sum to one over the polymorphic bins.

**Figure A2:**
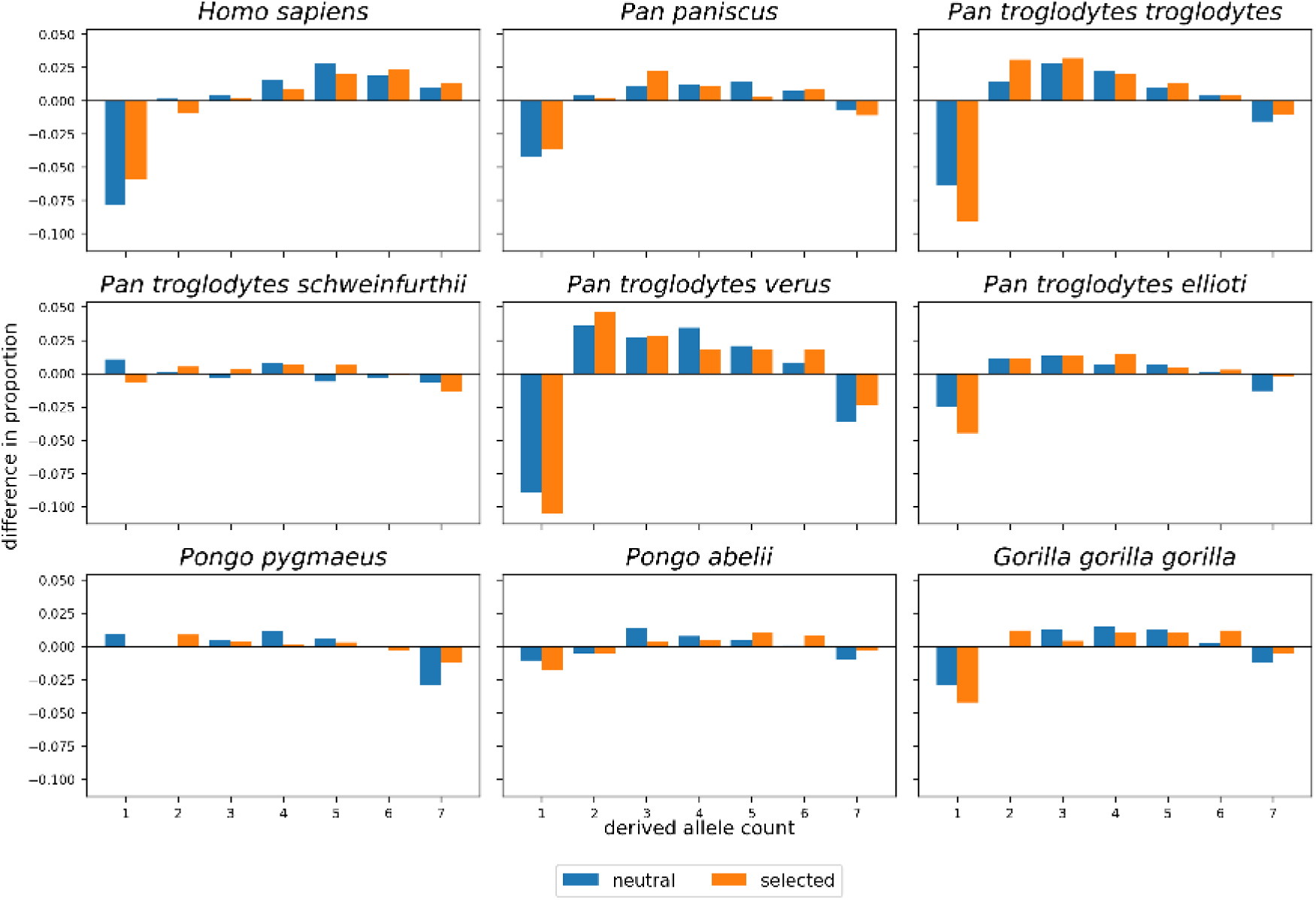
SFS comparison. Per-bin difference between the *neutral* and *selected* spectra of this study and those reported by Castellano et al. (2019) for the same nine great apes. Each spectrum is normalized over its polymorphic bins before differencing, and positive values denote a larger proportion in this study. Their spectra are mapped to the human reference genome, whereas ours are mapped to each species’ own reference. It is not clear whether CpG sites are included in their spectra.

**Figure A3:**
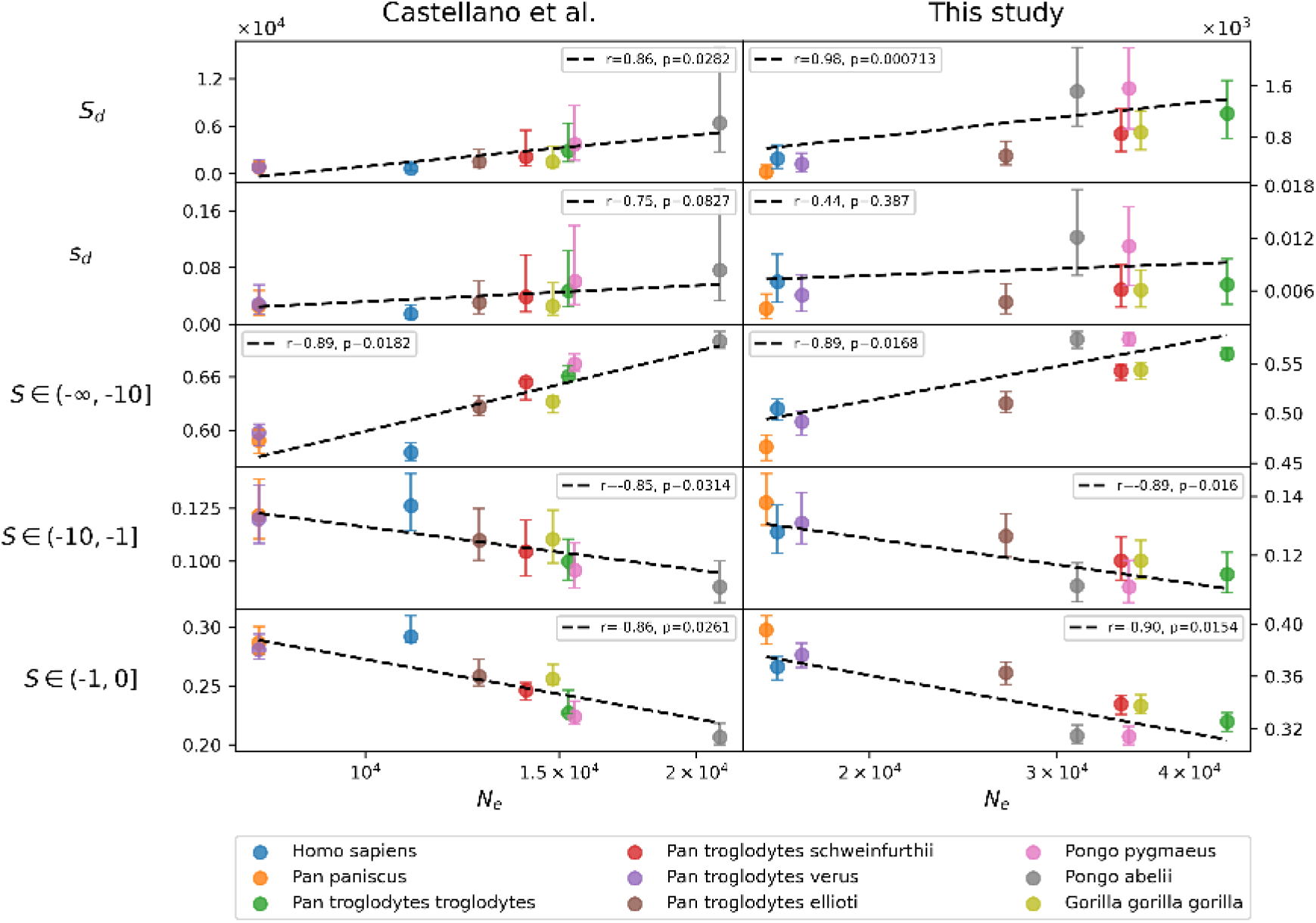
Comparison of the correlation between *N_e_* and DFE parameters with Castellano et al. (2019), using an unfolded SFS (*n* = 8). Both analyses use GammaExpParametrization and fit the SFS under a shape parameter *b* shared across species. No correction for ancestral misidentification is applied, and only the deleterious component of the DFE is estimated, corresponding to model 3S in Castellano et al. (2019). Of the two variants they report, the comparison uses the estimates for all mutations rather than those restricted to GC-conservative sites, and our analysis likewise includes all mutations. Each column uses the *N_e_* estimates of its own study.

**Figure A4:**
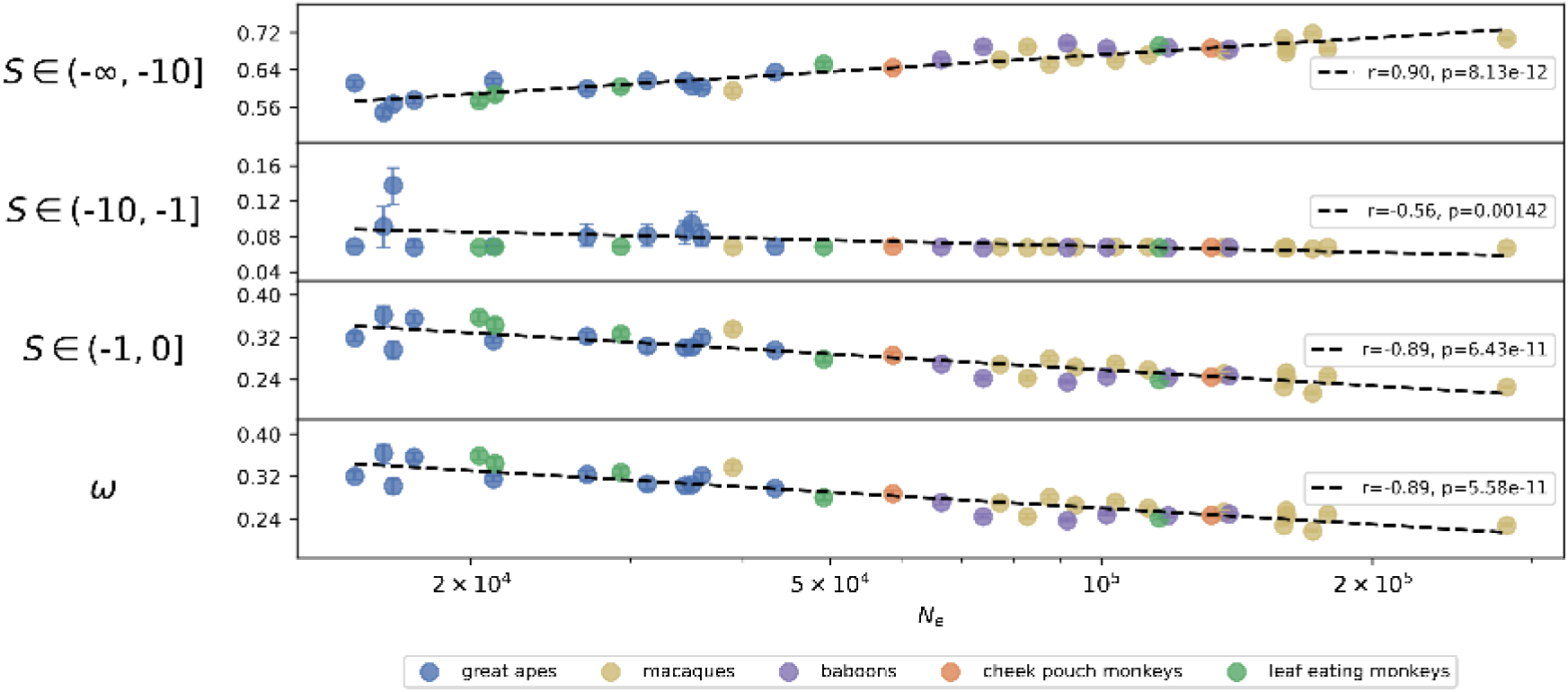
DFE for deleterious mutations. Correlation with *N_e_*, inferred from a folded SFS (*n* = 8) using GammaExpParametrization. The bottom panel shows the predicted total substitution rate *ω* = *dN/dS*. The qualitative pattern matches that in Figure 2.

**Figure A5:**
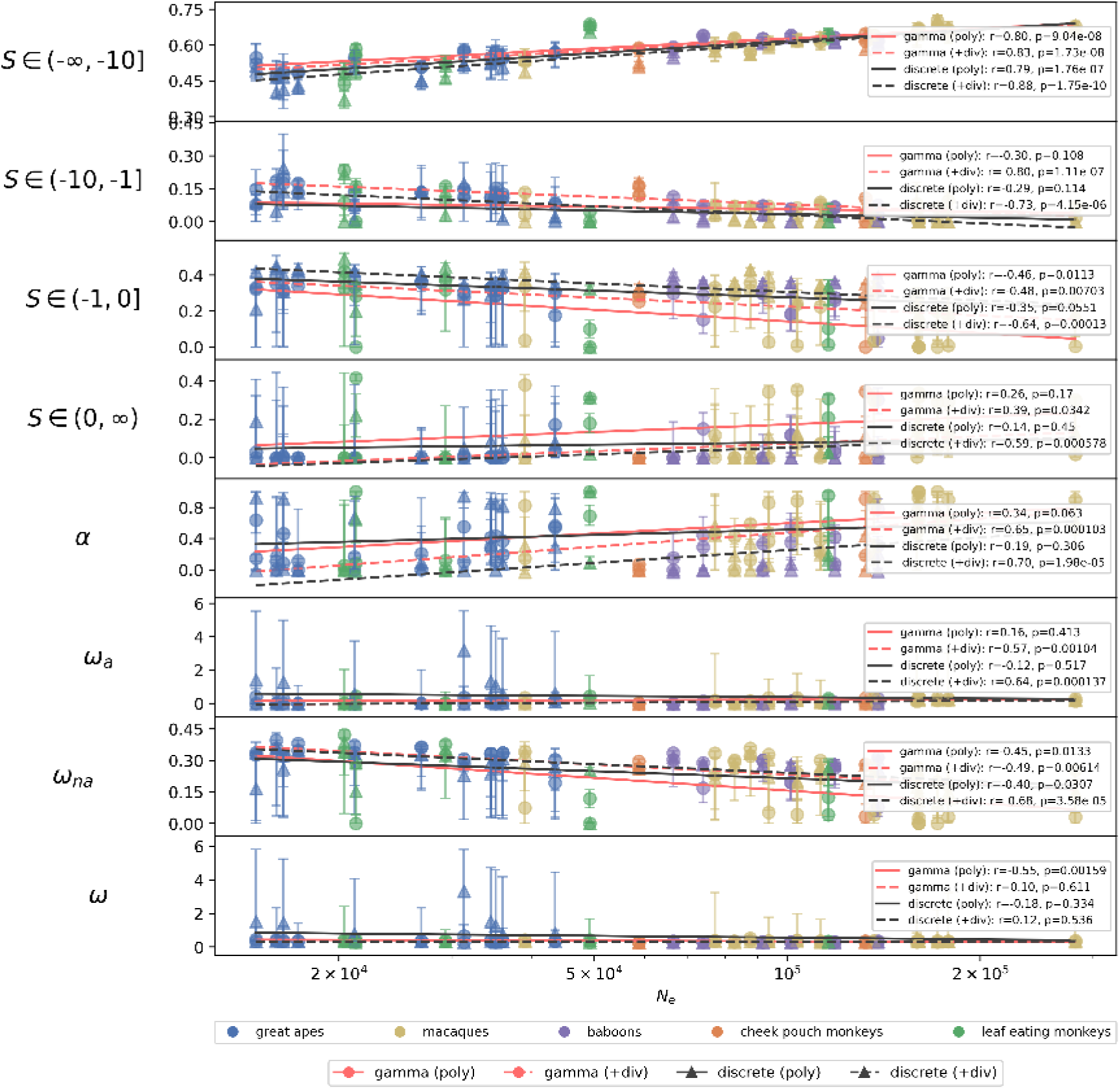
Full DFE versus *N_e_*, inferred from polymorphism data alone (solid lines) and jointly with lineage-specific divergence (dashed lines), under the GammaExpParametrization (red) and DiscreteParametrization (black). Panels show the fraction of mutations in each interval of *S*, the adaptive fraction *α*, the adaptive substitution rate *ω_a_*, the non-adaptive rate *ω_na_*, and the total substitution rate *ω* = *dN/dS*. Lineage-specific divergence counts are the fixed differences at 0-fold and 4-fold coding sites between each reference genome and the inferred ancestral allele (Methods). Adding divergence makes both *α* and *ω_a_* increase significantly with *N_e_*under either parametrization, while *ω* stays flat and *ω_na_* declines.

**Figure A6:**
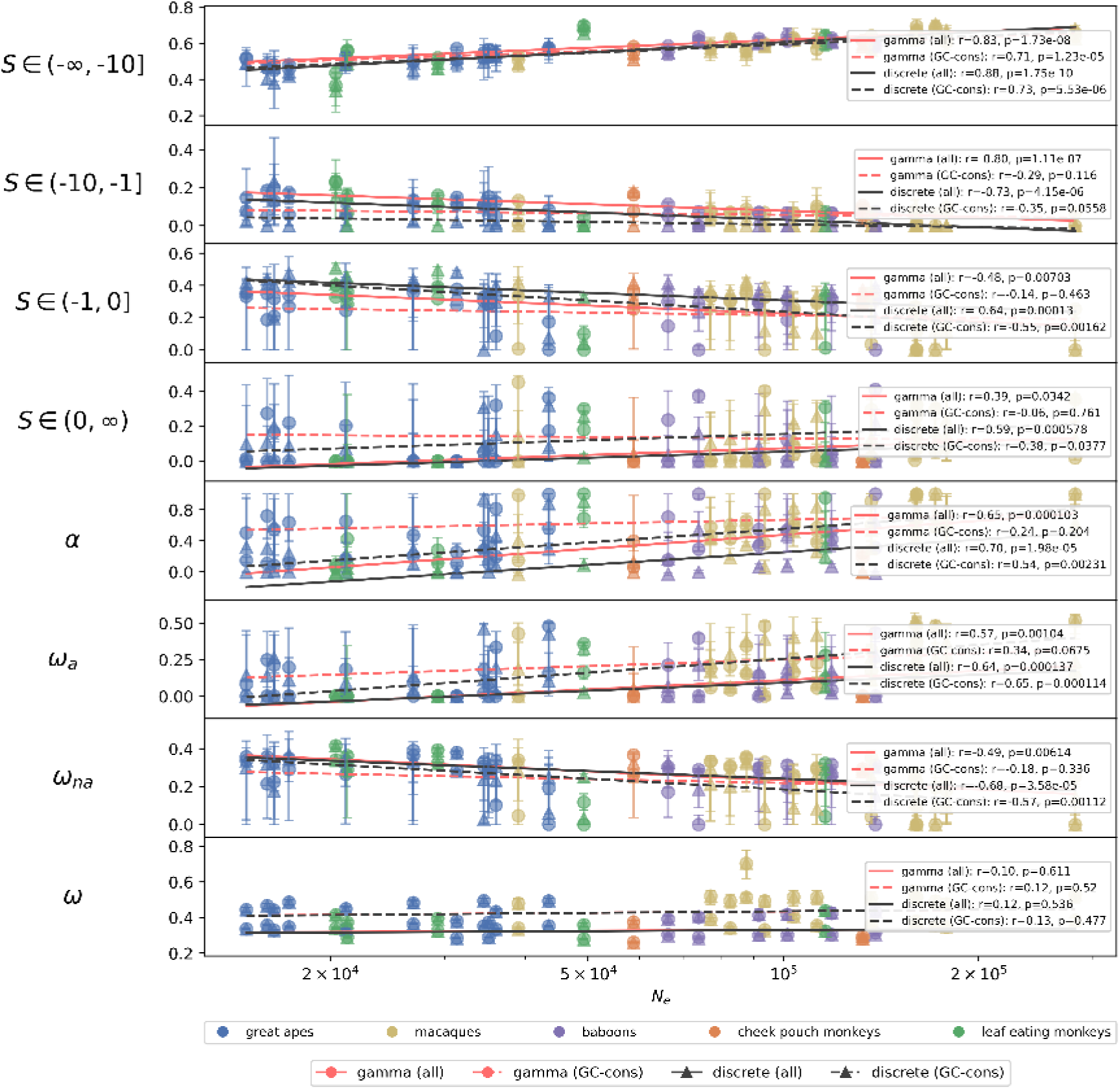
gBGC robustness of the divergence-informed adaptive signal. Full DFE statistics versus *N_e_*across Catarrhini with divergence incorporated, comparing the standard analysis with one further restricted to GC-conservative mutations (A↔T and C↔G; unaffected by gBGC), under both parametrizations. Under DiscreteParametrization the positive *ω_a_*–*N_e_* and *α*–*N_e_*associations persist on GC-conservative sites (*ω_a_*: *r* = 0.65, *p* = 1.1 × 10^−4^; *α*: *r* = 0.54, *p* = 2.3 × 10^−3^), indicating that the adaptive trend is not driven by gBGC. Under GammaExpParametrization the GC-conservative fit is degenerate: the deleterious mean *S_d_*is estimated at its lower bound for 17 of the 38 subspecies (spanning the full *N_e_* range), so that even the *ω_na_*–*N_e_* relationship—robustly negative in all other analyses—is not significant.

**Figure A7:**
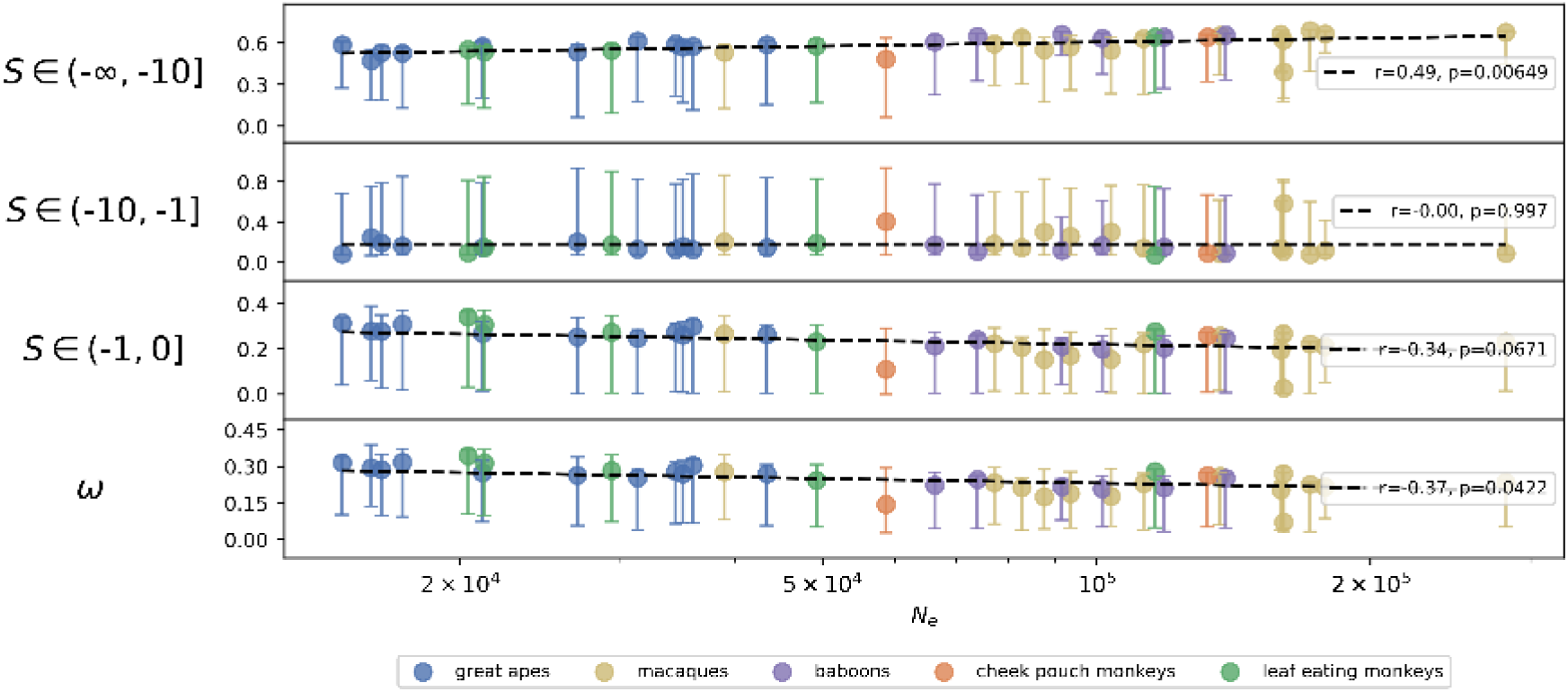
Low sample size DFE. The DFE for deleterious mutations, inferred from an SFS (*n* = 2) using GammaExpParametrization. The bottom panel shows the predicted total substitution rate *ω* = *dN/dS*. At *n* = 2 it is poorly constrained, with wide confidence intervals.

**Figure A8:**
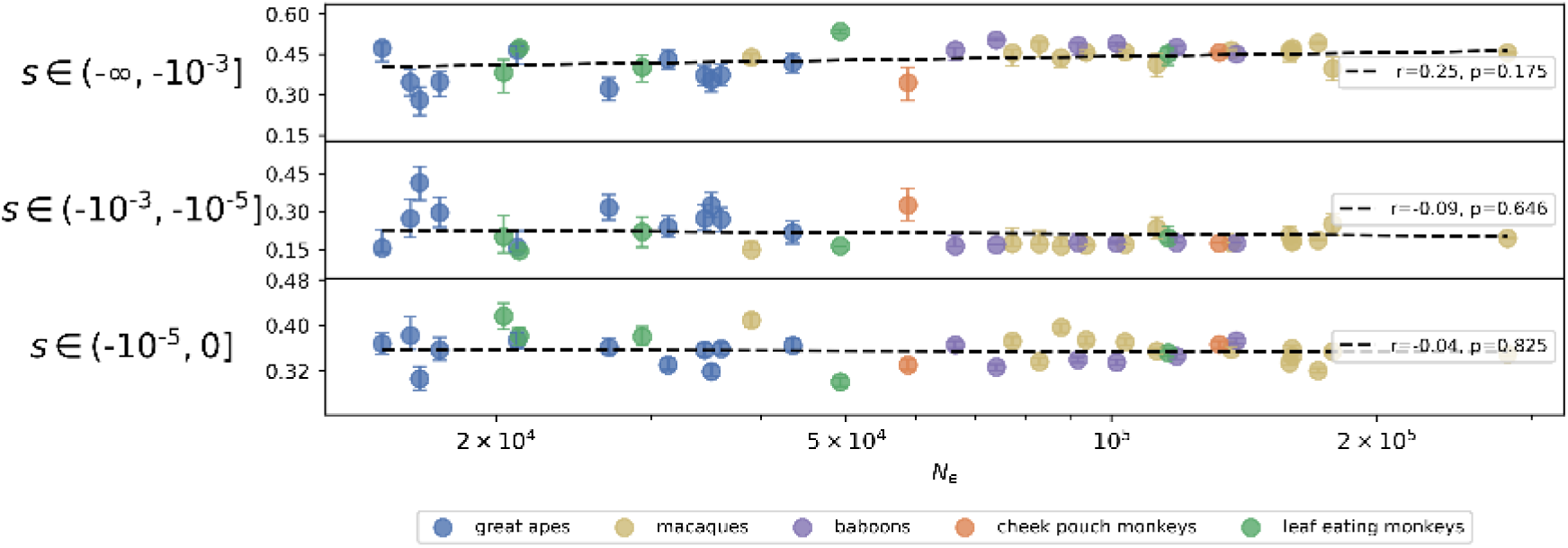
Unscaled DFE for deleterious mutations. Inferred from an SFS (*n* = 8) using GammaExpParametrization, rescaled from *S* to *s* by dividing by 4*N_e_*.

**Figure A9:**
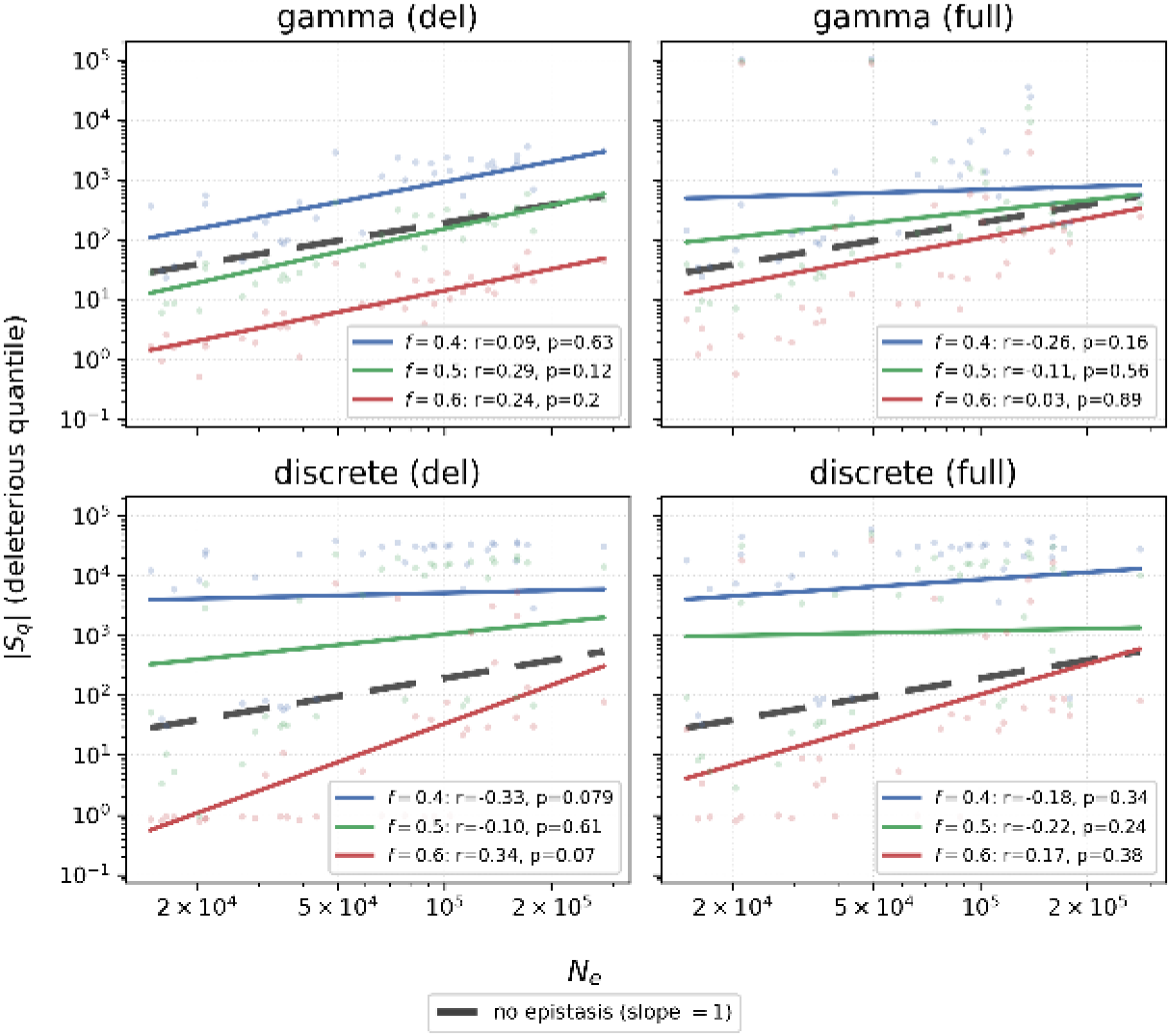
Tail-robust quantile test for epistasis on the DFE for deleterious mutations. PGLS regression of the deleterious-mass quantile log_10_ |*S_q_*| on log_10_ *N_e_*across catarrhine species, from unfolded SFS with *n* = 8, where *S_q_* is the population-scaled coefficient below which a fraction *f* of the deleterious mass lies. Each panel shows one DFE-model variant, with the three quantile levels *f* distinguished by color. Faint points are subspecies-level medians and lines are PGLS fits on species-averaged values. Under no epistasis a fixed quantile rescales as *S_q_* = 4*N_e_s_q_*, so the slope equals 1 (bold dark dashed reference in each panel). Panel legends report *r* and *p* for the no-epistasis null hypothesis, with the corresponding PGLS slopes given in Table 1. On the simulated *N_e_*-rescaled dataset with a fixed unscaled DFE (Figure A15), the quantile slope is exactly 1 at every *f*.

**Figure A10:**
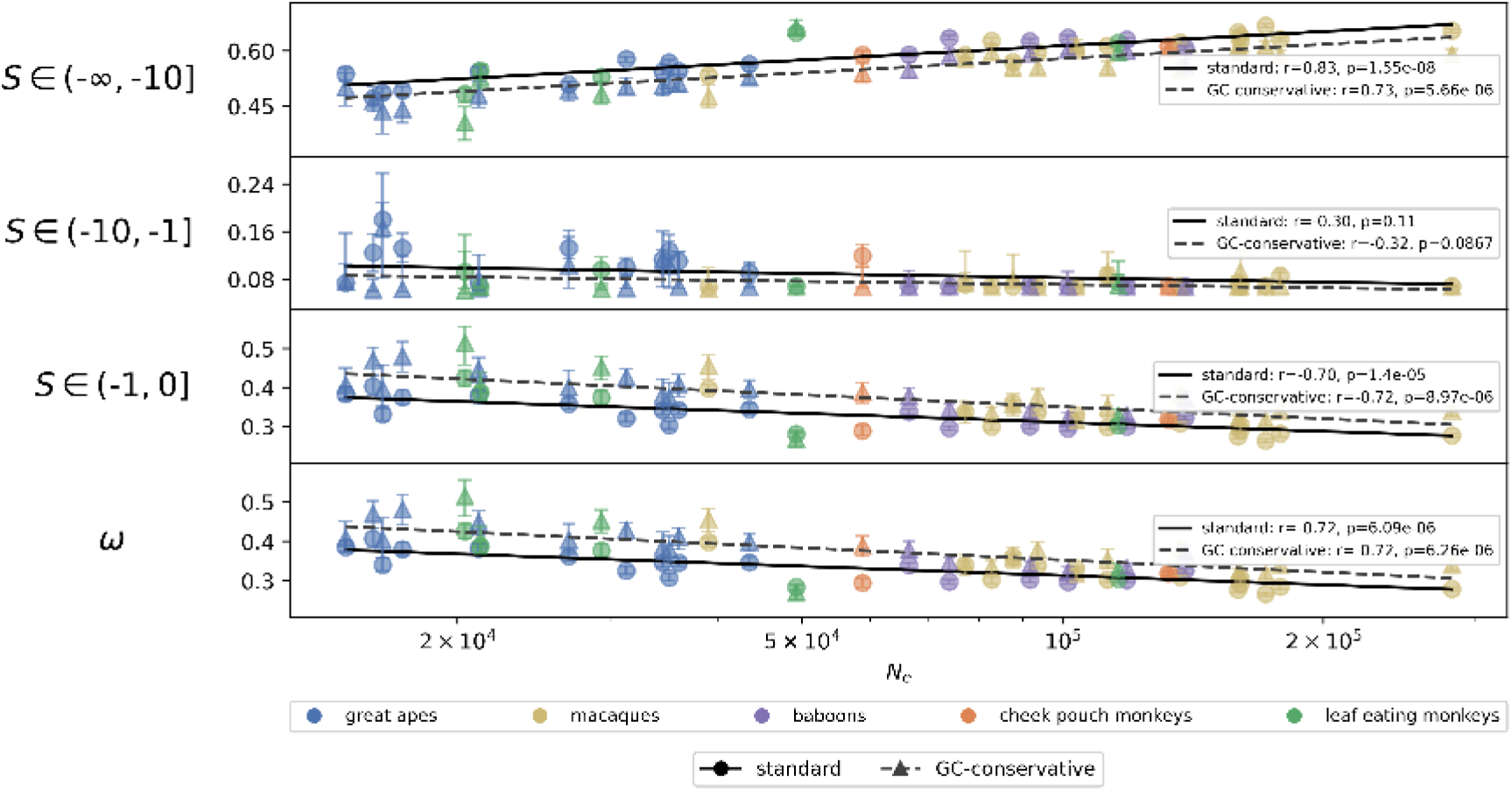
Effect of GC-biased gene conversion on DFE inference. Correlation between the DFE for deleterious mutations and *N_e_*across Catarrhini, comparing the standard analysis with one further restricted to GC-conservative mutations only (A↔T and C↔G), which are unaffected by GC-biased gene conversion (gBGC). DFEs were inferred from the unfolded SFS (*n* = 8) using GammaExpParametrization, estimating only the deleterious component.

**Figure A11:**
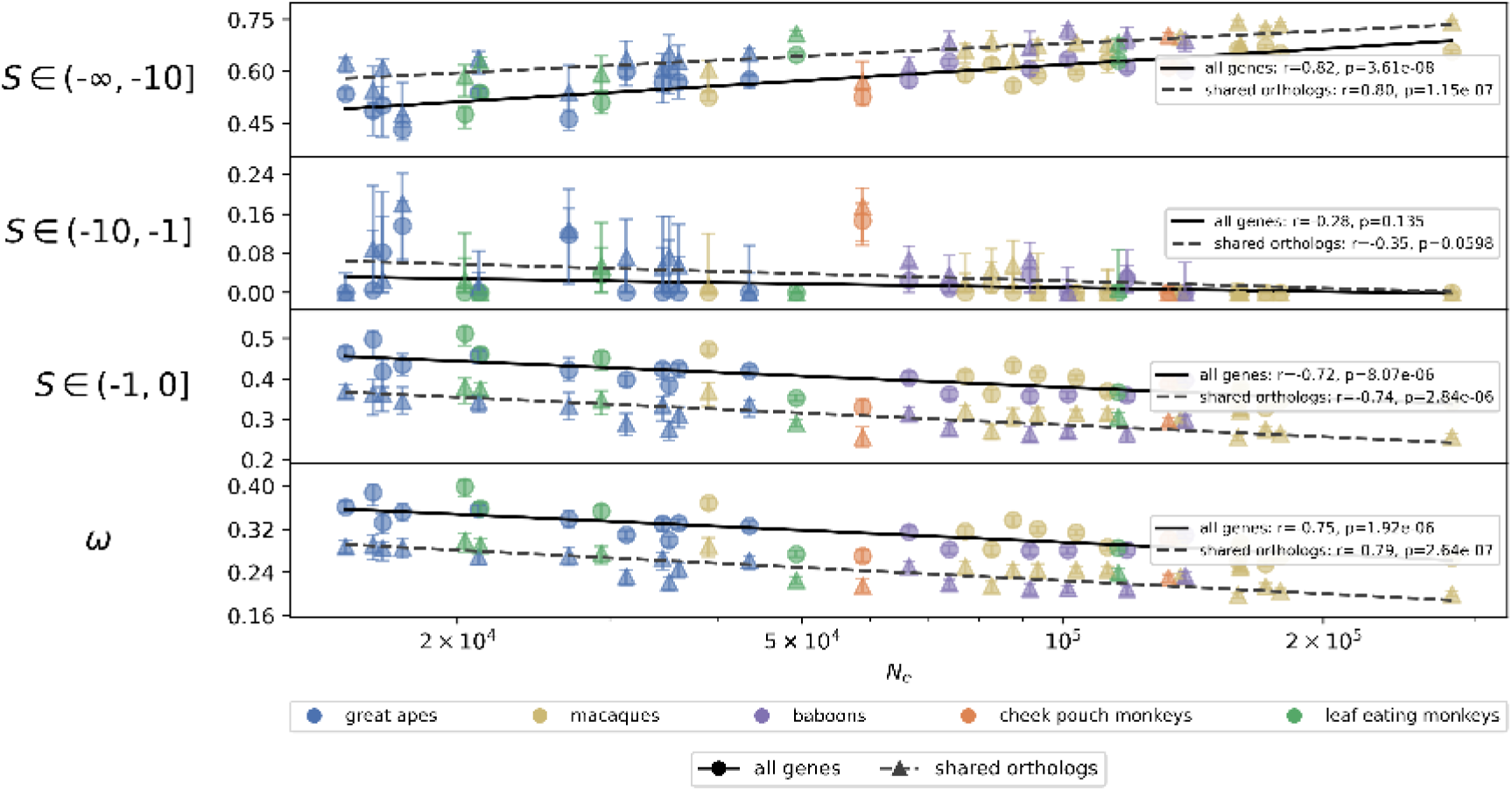
Effect on DFE inference of restricting to a common gene set. Correlation between the DFE for deleterious mutations and *N_e_* across Catarrhini, comparing the standard analysis, which uses all genes annotated in each reference, with one restricted to the 14,437 ortholog groups annotated in all 14 references. These groups cover ∼75% of coding sequence per reference (72–80%), with 4-fold and 0-fold sites represented in nearly equal proportion (74.5% and 75.3% on average). They nevertheless contain only 60% of *selected* SNPs against 72% of *neutral* ones, consistent with stronger selective constraint in these genes. DFEs were inferred from the unfolded SFS (*n* = 8) using DiscreteParametrization, estimating only the deleterious component.

**Figure A12:**
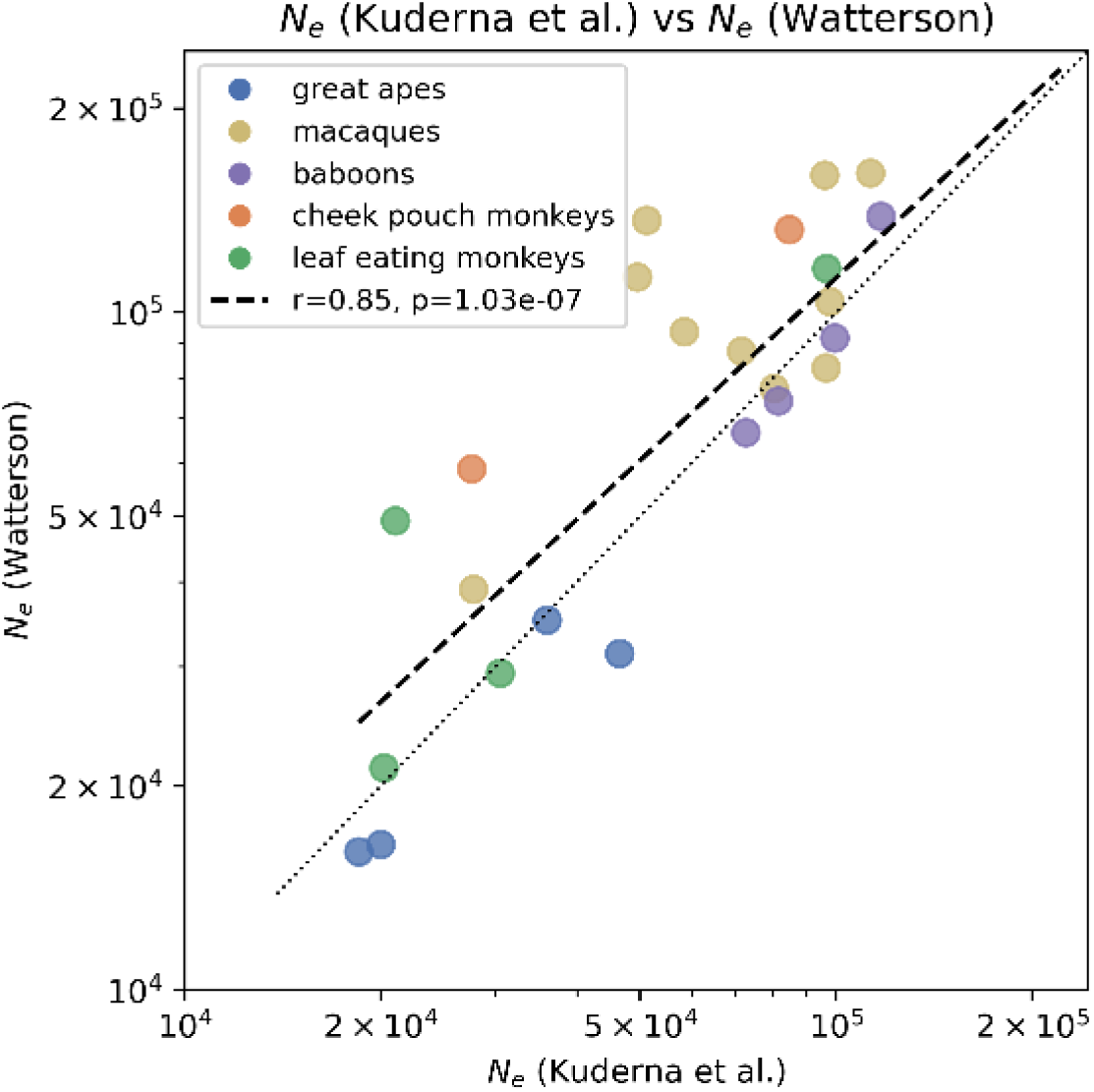
*N_e_* comparison. Comparison between our (Watterson-based) *N_e_*estimates and those from Kuderna et al. (2023). The dashed line is a PGLS regression of log_10_ *N_e_* (Watterson) on log_10_ *N_e_* (Kuderna) across catarrhine species. The dotted line is *y* = *x*.

**Figure A13:**
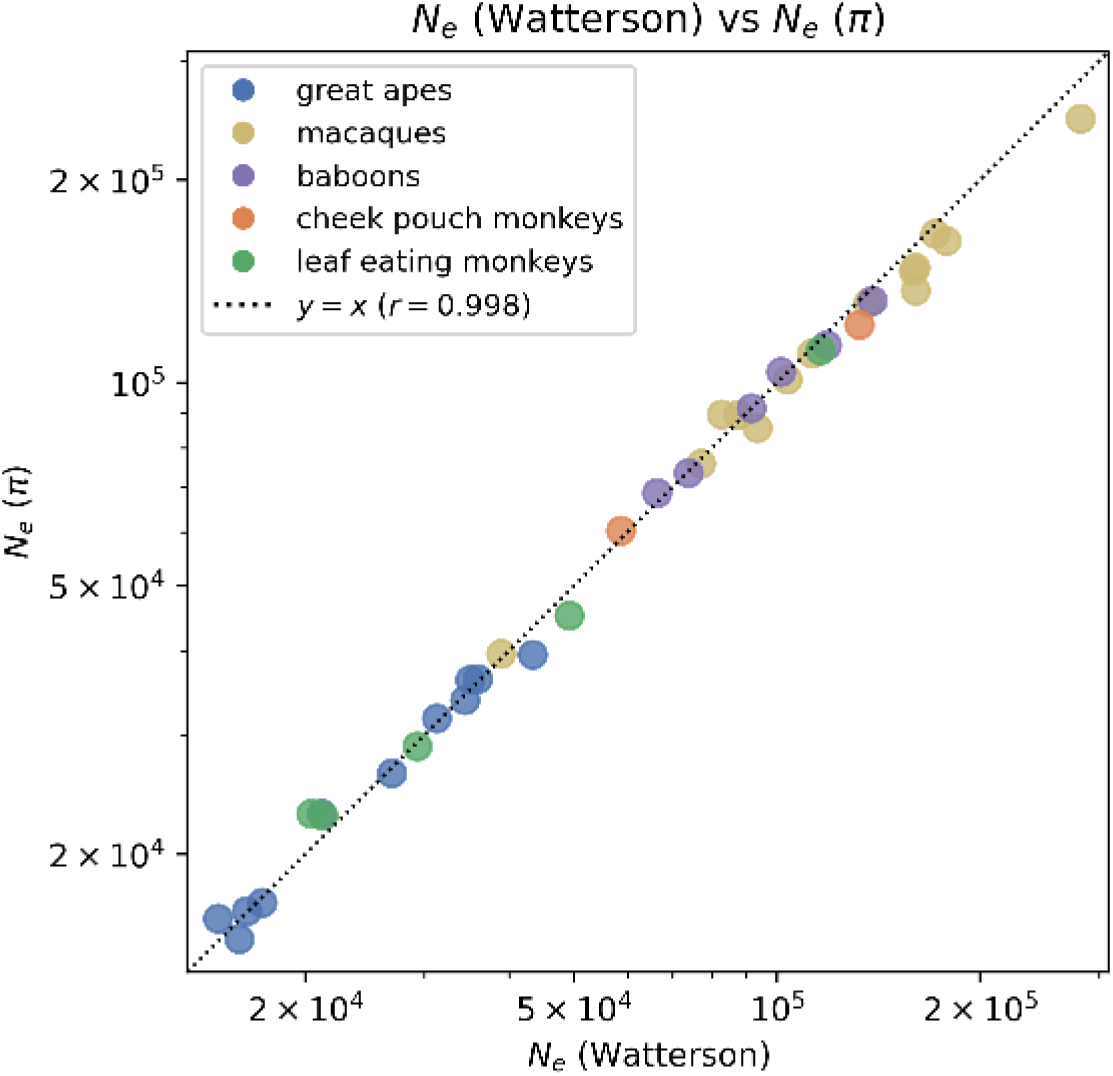
Effect of the *θ* estimator on *N_e_*. *N_e_* based on nucleotide diversity *π*^ against *N_e_* based on Watterson’s estimator, for each catarrhine population. Both estimate *θ* = 4*N_e_µ* and use the same unpolarized folded SFS (*n* = 8) and species-specific mutation rates. The dotted line is *y* = *x*. Here *r* denotes the Pearson correlation between the two sets of estimates on the log_10_ scale.

**Figure A14:**
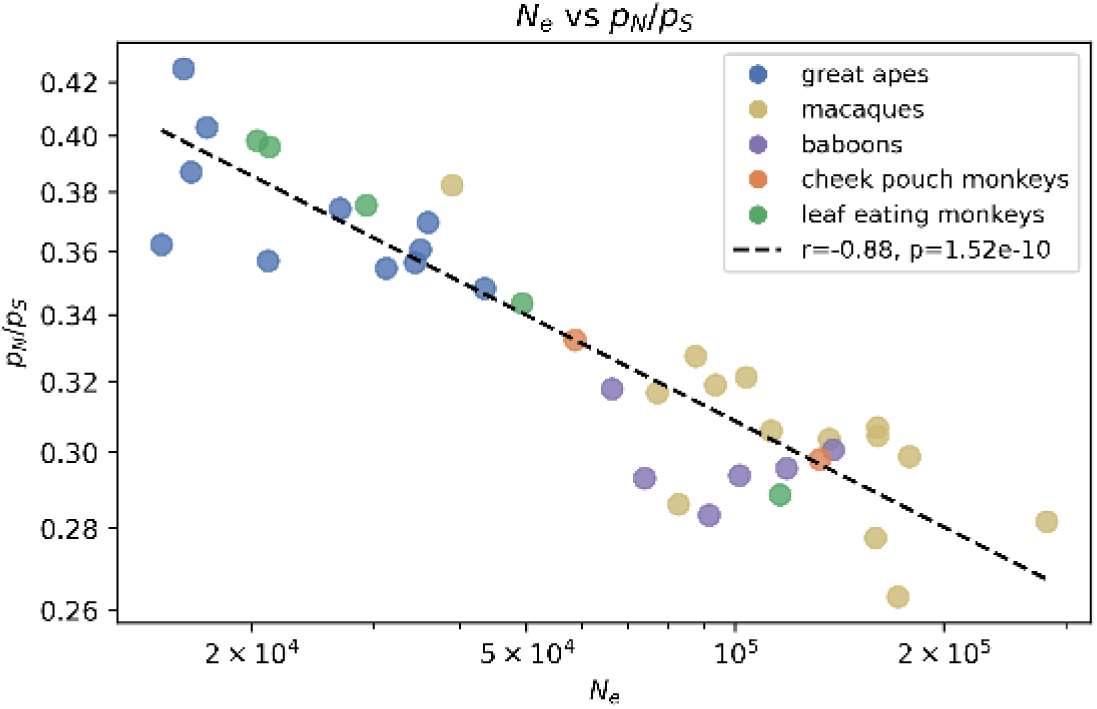
*N_e_*–*π_N_ /π_S_* associations. The dashed line is a PGLS regression of log_10_(*π_N_ /π_S_*) on log_10_ *N_e_* across catarrhine species.

**Figure A15:**
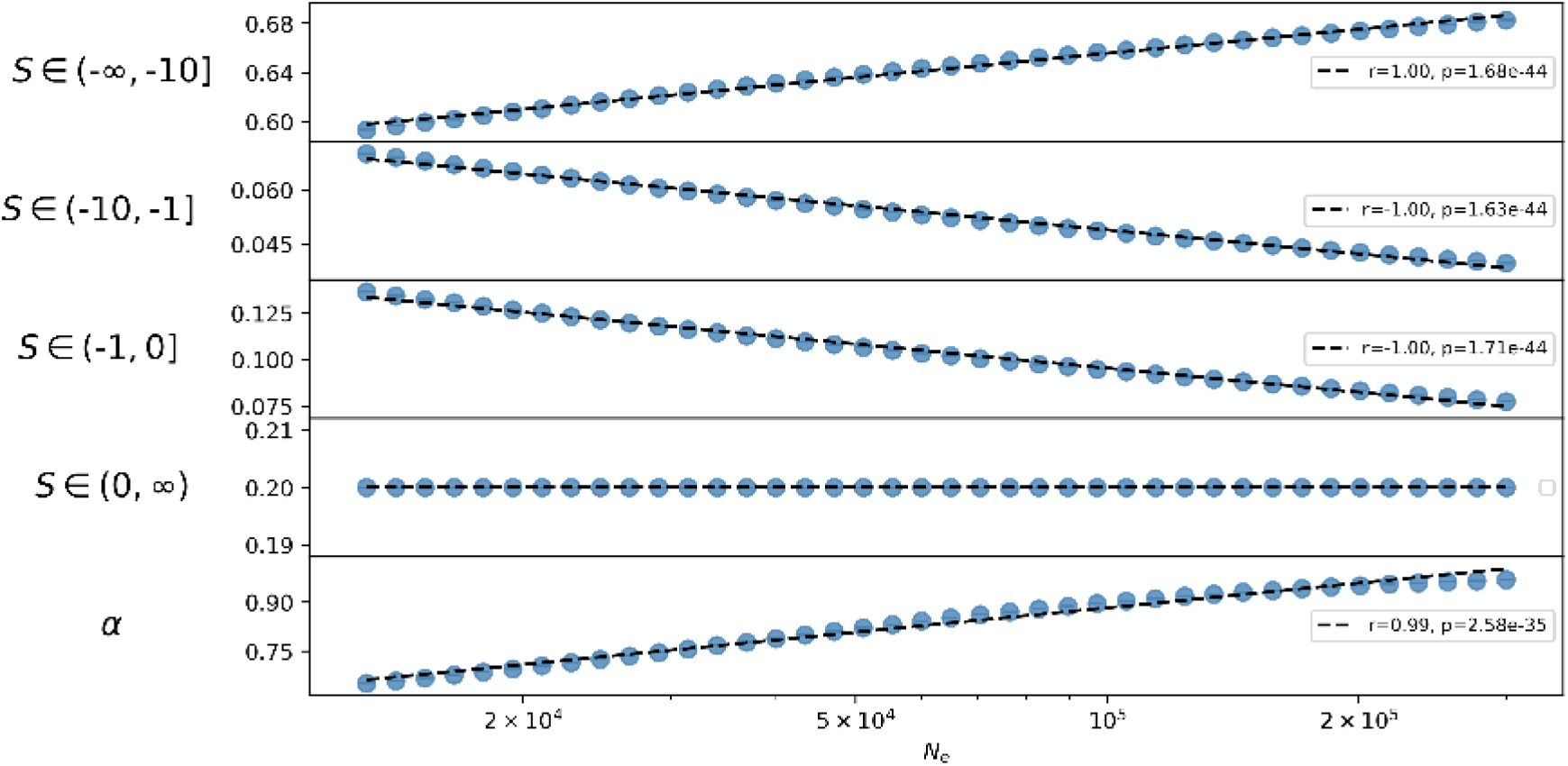
Simulated population-scaled DFE under a fixed underlying unscaled DFE. Under GammaExpParametrization, the unscaled DFE is a gamma distribution with mean *s_d_* and shape *b* for the deleterious mutations (fraction 1 − *p_b_*) and an exponential distribution with mean *s_b_* for the weakly beneficial mutations. The parameters were *s_d_* = −0.1, *b* = 0.18, *p_b_* = 0.2, and *s_b_* = 10^−5^, chosen to approximately match the population-scaled DFE in Figure 2. *N_e_* values were selected to span the same range as in the original dataset, and *s_d_* and *s_b_* were scaled by 4*N_e_* for each species.

**Figure A16:**
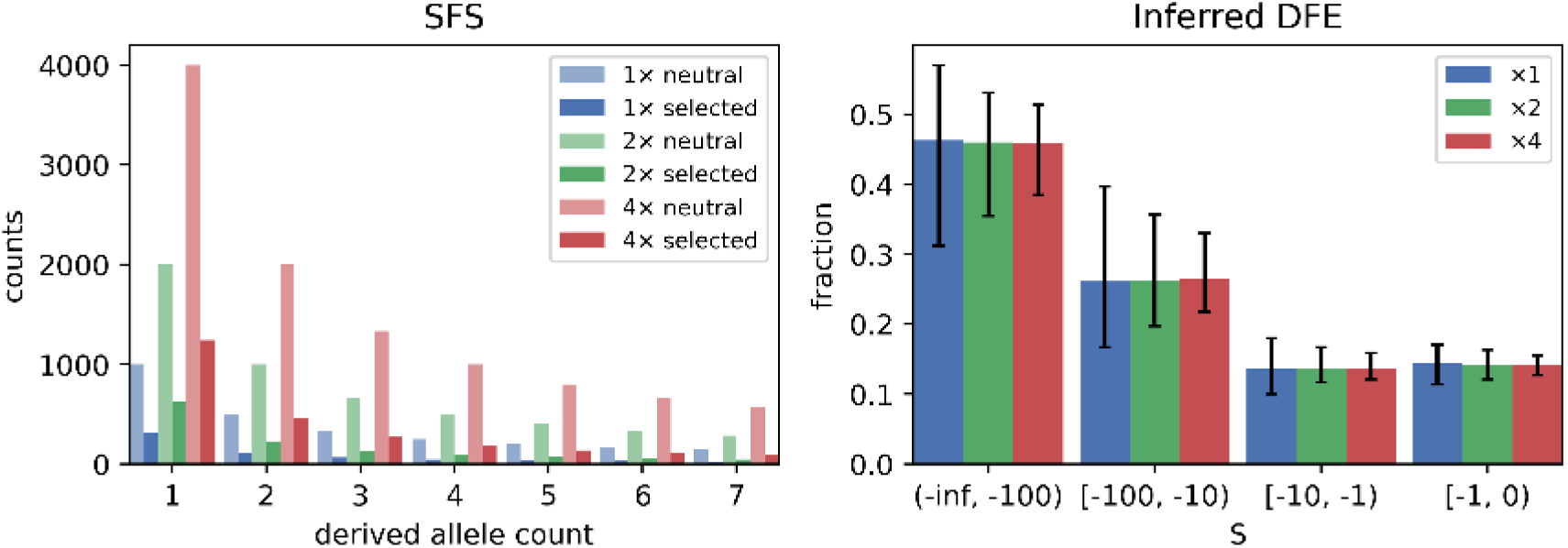
DFE invariance. Illustration of DFE shape invariance under uniform rescaling of the *neutral*and *selected* SFS by a scalar. Counts in the site-frequency spectra were simulated under a DFE with *S_d_* = −300, *b* = 0.3, and *p_b_* = 0 using GammaExpParametrization, and were subsequently multiplied by factors of 1, 2, or 4.

**Figure A17:**
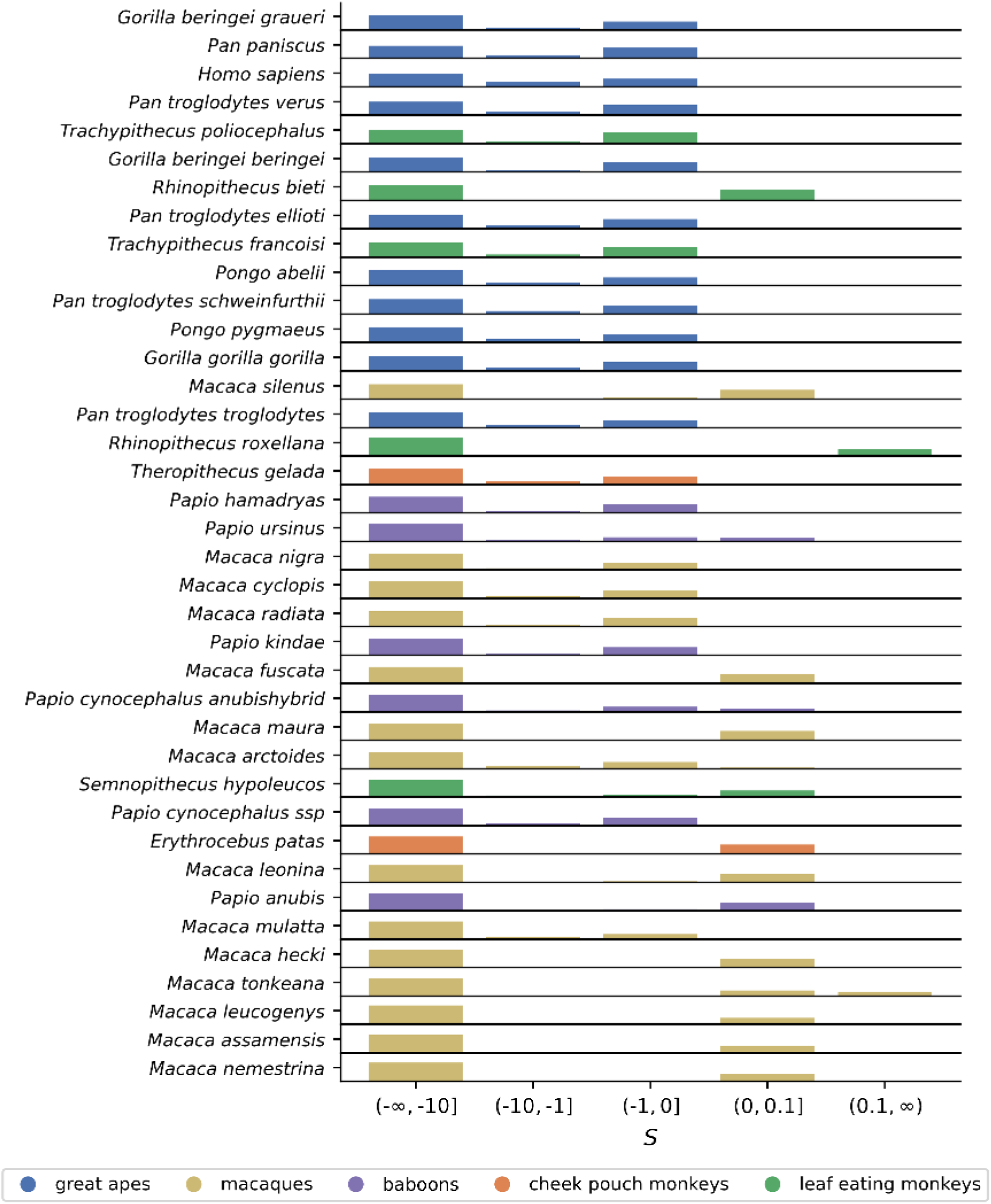
Discretized full DFEs. Discretized DFE for each subspecies, inferred from the unfolded SFS (*n* = 8) using GammaExpParametrization. Subspecies are ordered from top to bottom by increasing *N_e_*. Bar heights represent the median proportion of the DFE in the bin across bootstrap replicates.

## B Supplementary Tables

**Table B1:** Outgroups used for each subspecies in the analysis.

| Subspecies | Outgroups |
| --- | --- |
| *Homo sapiens* | *Pan troglodytes*, *Gorilla gorilla*, *Pongo abelii* |
| *Pan paniscus* | *Homo sapiens*, *Gorilla gorilla*, *Pongo abelii* |
| *Pan troglodytes ellioti* | *Homo sapiens*, *Gorilla gorilla*, *Pongo abelii* |
| *Pan troglodytes schweinfurthii* | *Homo sapiens*, *Gorilla gorilla*, *Pongo abelii* |
| *Pan troglodytes troglodytes* | *Homo sapiens*, *Gorilla gorilla*, *Pongo abelii* |
| *Pan troglodytes verus* | *Homo sapiens*, *Gorilla gorilla*, *Pongo abelii* |
| *Gorilla beringei beringei* | *Homo sapiens*, *Pongo abelii* |
| *Gorilla beringei graueri* | *Homo sapiens*, *Pongo abelii* |
| *Gorilla gorilla gorilla* | *Homo sapiens*, *Pongo abelii* |
| *Pongo abelii* | *Homo sapiens*, *Nomascus leucogenys* |
| *Pongo pygmaeus* | *Homo sapiens*, *Nomascus leucogenys* |
| *Macaca cyclopis* | *Macaca thibetana*, *Macaca nemestrina*, *Papio anubis* |
| *Macaca fuscata* | *Macaca thibetana*, *Macaca nemestrina*, *Papio anubis* |
| *Macaca mulatta* | *Macaca thibetana*, *Macaca nemestrina*, *Papio anubis* |
| *Macaca hecki* | *Macaca nemestrina*, *Macaca mulatta*, *Papio anubis* |
| *Macaca leonina* | *Macaca nemestrina*, *Macaca mulatta*, *Papio anubis* |
| *Macaca maura* | *Macaca nemestrina*, *Macaca mulatta*, *Papio anubis* |
| *Macaca nemestrina* | *Macaca leonina*, *Macaca mulatta*, *Papio anubis* |
| *Macaca nigra* | *Macaca nemestrina*, *Macaca mulatta*, *Papio anubis* |
| *Macaca radiata* | *Macaca mulatta*, *Macaca nemestrina*, *Papio anubis* |
| *Macaca silenus* | *Macaca nemestrina*, *Macaca mulatta*, *Papio anubis* |
| *Macaca tonkeana* | *Macaca nemestrina*, *Macaca mulatta*, *Papio anubis* |
| *Macaca arctoides* | *Macaca mulatta*, *Macaca nemestrina*, *Papio anubis* |
| *Macaca assamensis* | *Macaca mulatta*, *Macaca nemestrina*, *Papio anubis* |
| *Macaca leucogenys* | *Macaca mulatta*, *Macaca nemestrina*, *Papio anubis* |
| *Papio anubis* | *Papio hamadryas*, *Papio kindae*, *Macaca mulatta* |
| *Papio cynocephalus anubishybrid* | *Papio ursinus*, *Papio anubis*, *Macaca mulatta* |
| *Papio cynocephalus ssp* | *Papio ursinus*, *Papio anubis*, *Macaca mulatta* |
| *Papio hamadryas* | *Papio anubis*, *Papio kindae*, *Macaca mulatta* |
| *Papio kindae* | *Papio cynocephalus*, *Papio anubis*, *Macaca mulatta* |
| *Papio ursinus* | *Papio cynocephalus*, *Papio anubis*, *Macaca mulatta* |
| *Theropithecus gelada* | *Papio anubis*, *Macaca mulatta*, *Allenopithecus nigroviridis* |
| *Erythrocebus patas* | *Chlorocebus sabaeus*, *Papio anubis*, *Macaca mulatta* |
| *Rhinopithecus bieti* | *Rhinopithecus roxellana*, *Trachypithecus francoisi*, *Colobus guereza* |
| *Rhinopithecus roxellana* | *Pygathrix cinerea*, *Trachypithecus francoisi*, *Colobus guereza* |
| *Semnopithecus hypoleucos* | *Trachypithecus francoisi*, *Rhinopithecus roxellana*, *Colobus guereza* |
| *Trachypithecus francoisi* | *Semnopithecus entellus*, *Colobus guereza* |
| *Trachypithecus poliocephalus* | *Trachypithecus geei*, *Semnopithecus entellus*, *Colobus guereza* |

**Table B2:**
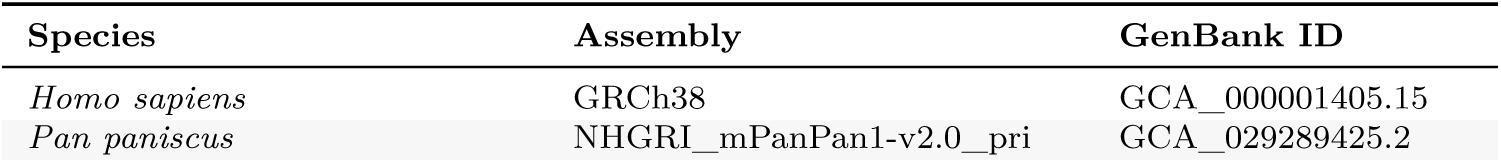
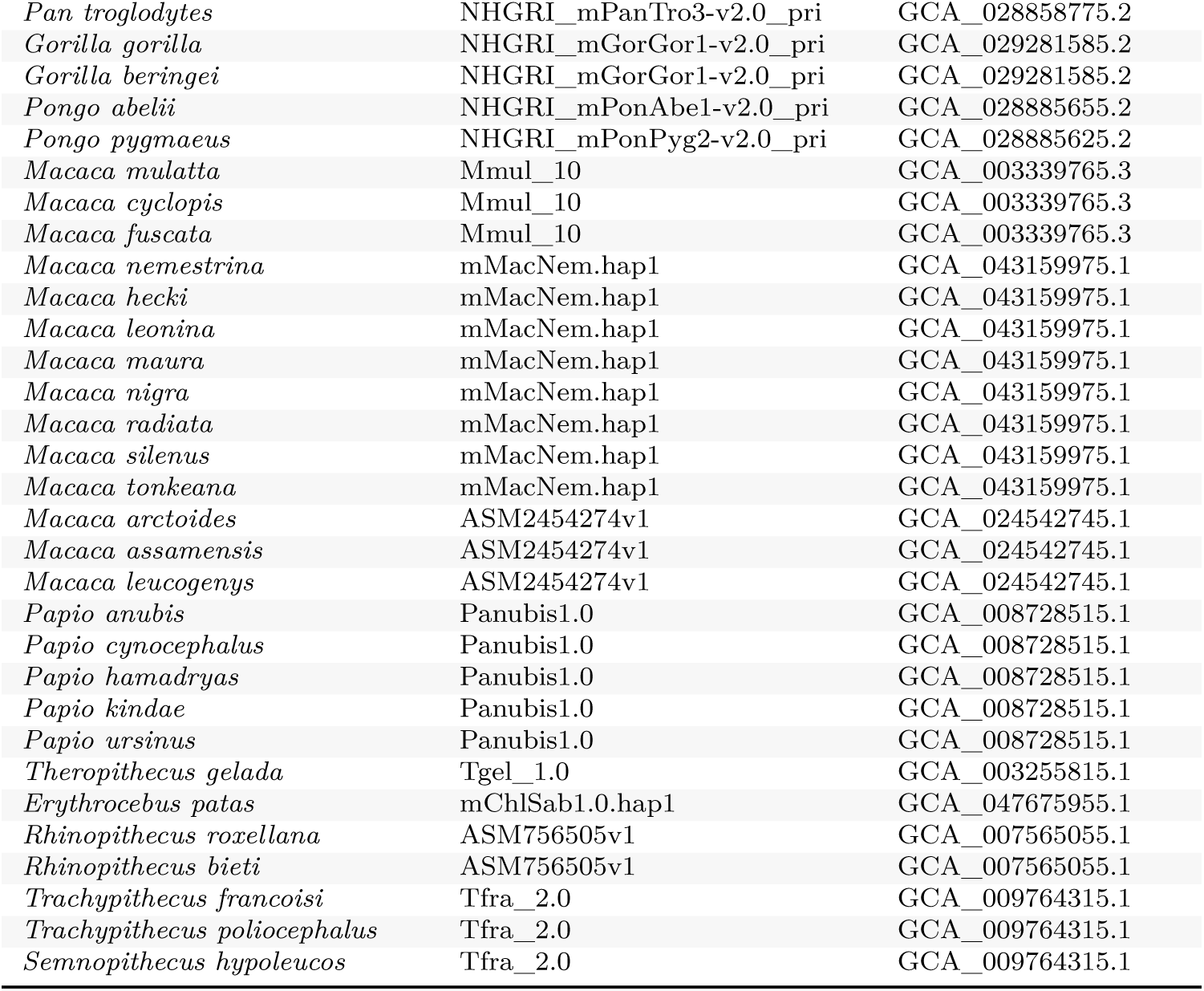
Reference genome assembly used for each species. Closely related species are mapped to the same assembly when no species-specific reference is available.

| Species | Assembly | GenBank ID |
| --- | --- | --- |
| *Homo sapiens* | GRCh38 | GCA_000001405.15 |
| *Pan paniscus* | NHGRI_mPanPan1-v2.0_pri | GCA_029289425.2 |

| Species | Assembly | GenBank ID |
| --- | --- | --- |
| *Pan troglodytes* | NHGRI_mPanTro3-v2.0_pri | GCA_028858775.2 |
| *Gorilla gorilla* | NHGRI_mGorGor1-v2.0_pri | GCA_029281585.2 |
| *Gorilla beringei* | NHGRI_mGorGor1-v2.0_pri | GCA_029281585.2 |
| *Pongo abelii* | NHGRI_mPonAbe1-v2.0_pri | GCA_028885655.2 |
| *Pongo pygmaeus* | NHGRI_mPonPyg2-v2.0_pri | GCA_028885625.2 |
| *Macaca mulatta* | Mmul_10 | GCA_003339765.3 |
| *Macaca cyclopis* | Mmul_10 | GCA_003339765.3 |
| *Macaca fuscata* | Mmul_10 | GCA_003339765.3 |
| *Macaca nemestrina* | mMacNem.hap1 | GCA_043159975.1 |
| *Macaca hecki* | mMacNem.hap1 | GCA_043159975.1 |
| *Macaca leonina* | mMacNem.hap1 | GCA_043159975.1 |
| *Macaca maura* | mMacNem.hap1 | GCA_043159975.1 |
| *Macaca nigra* | mMacNem.hap1 | GCA_043159975.1 |
| *Macaca radiata* | mMacNem.hap1 | GCA_043159975.1 |
| *Macaca silenus* | mMacNem.hap1 | GCA_043159975.1 |
| *Macaca tonkeana* | mMacNem.hap1 | GCA_043159975.1 |
| *Macaca arctoides* | ASM2454274v1 | GCA_024542745.1 |
| *Macaca assamensis* | ASM2454274v1 | GCA_024542745.1 |
| *Macaca leucogenys* | ASM2454274v1 | GCA_024542745.1 |
| *Papio anubis* | Panubis1.0 | GCA_008728515.1 |
| *Papio cynocephalus* | Panubis1.0 | GCA_008728515.1 |
| *Papio hamadryas* | Panubis1.0 | GCA_008728515.1 |
| *Papio kindae* | Panubis1.0 | GCA_008728515.1 |
| *Papio ursinus* | Panubis1.0 | GCA_008728515.1 |
| *Theropithecus gelada* | Tgel_1.0 | GCA_003255815.1 |
| *Erythrocebus patas* | mChlSab1.0.hap1 | GCA_047675955.1 |
| *Rhinopithecus roxellana* | ASM756505v1 | GCA_007565055.1 |
| *Rhinopithecus bieti* | ASM756505v1 | GCA_007565055.1 |
| *Trachypithecus francoisi* | Tfra_2.0 | GCA_009764315.1 |
| *Trachypithecus poliocephalus* | Tfra_2.0 | GCA_009764315.1 |
| *Semnopithecus hypoleucos* | Tfra_2.0 | GCA_009764315.1 |

**Table B3:** Per-subspecies mutation rate and inferred effective population size. Mutation rates *µ* (per site per generation) are taken from Kuderna et al. (2023) at the species level and applied to all subspecies of that species; for the four subspecies absent from Kuderna et al. (2023) (*T. poliocephalus*, *M. hecki*, *M. leucogenys*, *M. assamensis*), *µ* is the value reported for the reference species onto which they were mapped. Effective population sizes *N_e_* are estimated as 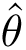/(4*µ*) using Watterson’s estimator on neutral (4-fold) sites (see Methods). Subspecies are ordered by *N_e_*.

| Subspecies | $\mu$ ($10^{-8}$/site/gen) | $N_e$ ($10^4$) |
| --- | --- | --- |
| *Gorilla beringei* graueri | 1.20 | 1.49 |
| *Pan paniscus* | 1.17 | 1.60 |
| *Homo sapiens* | 1.20 | 1.64 |
| *Pan troglodytes* verus | 1.31 | 1.73 |
| *Trachypithecus poliocephalus* | 0.72 | 2.04 |
| *Gorilla beringei* beringei | 1.20 | 2.12 |
| *Rhinopithecus bieti* | 0.52 | 2.13 |
| *Pan troglodytes* ellioti | 1.31 | 2.69 |
| *Trachypithecus francoisi* | 0.72 | 2.93 |
| *Pongo abelii* | 1.47 | 3.13 |
| *Pan troglodytes* schweinfurthii | 1.31 | 3.45 |
| *Pongo pygmaeus* | 1.28 | 3.51 |
| *Gorilla gorilla* gorilla | 1.33 | 3.60 |
| *Macaca silenus* | 0.55 | 3.90 |
| *Pan troglodytes* troglodytes | 1.31 | 4.35 |
| *Rhinopithecus roxellana* | 0.49 | 4.92 |
| *Theropithecus gelada* | 0.44 | 5.87 |
| *Papio hamadryas* | 0.61 | 6.64 |
| *Papio ursinus* | 0.69 | 7.40 |

Table B3: Per-subspecies mutation rate and inferred effective population size.
| Subspecies | $\mu$ ($10^{-8}$/site/gen) | $N_e$ ($10^4$) |
| --- | --- | --- |
| *Macaca nigra* | 0.49 | 7.72 |
| *Macaca cyclopis* | 0.44 | 8.28 |
| *Macaca radiata* | 0.44 | 8.76 |
| *Papio kindae* | 0.72 | 9.17 |
| *Macaca fuscata* | 0.43 | 9.36 |
| *Papio cynocephalus* anubishybrid | 0.59 | 10.14 |
| *Macaca maura* | 0.54 | 10.37 |
| *Macaca arctoides* | 0.48 | 11.27 |
| *Semnopithecus hypoleucos* | 0.49 | 11.60 |
| *Papio cynocephalus* ssp | 0.59 | 11.86 |
| *Erythrocebus patas* | 0.49 | 13.25 |
| *Macaca leonina* | 0.51 | 13.67 |
| *Papio anubis* | 0.39 | 13.86 |
| *Macaca mulatta* | 0.56 | 15.93 |
| *Macaca hecki* | 0.37 | 16.04 |
| *Macaca tonkeana* | 0.53 | 16.05 |
| *Macaca leucogenys* | 0.39 | 17.16 |
| *Macaca assamensis* | 0.39 | 17.84 |
| *Macaca nemestrina* | 0.37 | 28.17 |

**Table B4:**
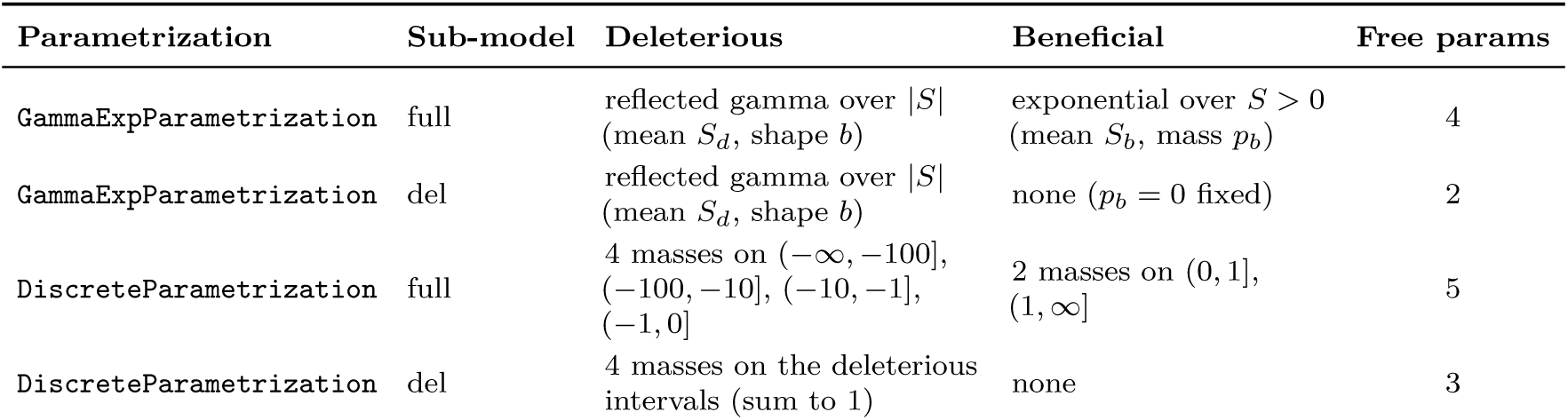
Parameters fitted under each DFE-model variant. The four variants used throughout the manuscript are two parametrizations (gamma–exponential and discrete) crossed with two sub-models (full = deleterious plus beneficial; del = deleterious only). The ancestral misidentification parameter *ε* is fixed to 0 and the dominance coefficient *h* to 0.5 in all main-text inferences (see Methods). The recessive comparison in Figure 3 of the main text uses an alternative *h*(*S*).

**Table B5:** Per-subspecies model comparison across the four DFE-model variants of Table B4 (GammaExpParametrization and DiscreteParametrization, each deleterious-only (*del*) or full (*full*)). LL is the log-likelihood at the MLE; AIC = 2*k* − 2 LL with free-parameter counts *k* as in Table B4. LRT *p*-values are from a *χ*^2^ test of the nested deleterious-only vs. full model within each parametrization: small *p* (*p <* 0.05) indicates that adding the beneficial component significantly improves the fit, while values close to one indicate that the simpler deleterious-only model is preferred. The final column reports ΔAIC = AIC*_γ_* _full_−AIC_discrete full_; positive values favor the discrete parametrization. The bottom row reports per-column means across subspecies (means are omitted for the LRT *p*-values, which are not meaningfully averaged). Overall, the deleterious-only model is preferred for 23/38 subspecies (LRT *p* ≥ 0.05 in both parametrizations), and the gamma and discrete full-DFE fits are close in AIC (mean ΔAIC = +4.0 in favor of discrete; |ΔAIC| *<* 5 in 21/38 subspecies), consistent with the qualitative agreement between the two parametrizations reported in the main text. Inferences use unfolded SFS at *n* = 8 haplotypes with *ε* = 0.

| Subspecies | GammaExp | | | | | Discrete | | | | | $\Delta\text{AIC}$ |
| --- | --- | --- | --- | --- | --- | --- | --- | --- | --- | --- | --- |
| | LL<sub>del</sub> | LL<sub>full</sub> | AIC<sub>del</sub> | AIC<sub>full</sub> | LRT $p$ | LL<sub>del</sub> | LL<sub>full</sub> | AIC<sub>del</sub> | AIC<sub>full</sub> | LRT $p$ | |
| *Homo sapiens* | -32.3 | -31.8 | 68.5 | 71.6 | 0.65 | -32.5 | -31.8 | 71.0 | 73.5 | 0.49 | -1.9 |
| *Pan paniscus* | -31.8 | -31.8 | 67.5 | 71.5 | 1.00 | -31.4 | -31.4 | 68.7 | 72.7 | 1.00 | -1.2 |
| *Pan troglodytes ellioti* | -37.4 | -37.4 | 78.7 | 82.7 | 1.00 | -33.8 | -33.8 | 73.6 | 77.6 | 0.98 | +5.2 |
| *Pan troglodytes schweinfurthii* | -34.4 | -34.2 | 72.8 | 76.4 | 0.81 | -34.9 | -34.0 | 75.8 | 77.9 | 0.38 | -1.5 |
| *Pan troglodytes troglodytes* | -34.5 | -34.3 | 73.1 | 76.6 | 0.78 | -35.5 | -34.1 | 77.0 | 78.2 | 0.25 | -1.6 |
| *Pan troglodytes verus* | -40.4 | -40.4 | 84.7 | 88.7 | 1.00 | -33.2 | -33.2 | 72.4 | 76.4 | 1.00 | +12.3 |
| *Gorilla beringei beringei* | -34.3 | -34.3 | 72.5 | 76.5 | 1.00 | -33.7 | -33.1 | 73.4 | 76.1 | 0.54 | +0.4 |
| *Gorilla beringei graueri* | -36.0 | -34.4 | 76.0 | 76.8 | 0.20 | -39.9 | -34.2 | 85.8 | 78.3 | $3.1 \times 10^{-3}$ | -1.5 |
| *Gorilla gorilla gorilla* | -34.8 | -34.5 | 73.5 | 77.0 | 0.77 | -35.6 | -34.5 | 77.1 | 79.0 | 0.34 | -2.0 |
| *Pongo abelii* | -39.2 | -36.4 | 82.4 | 80.8 | 0.06 | -41.7 | -35.0 | 89.3 | 80.1 | $1.3 \times 10^{-3}$ | +0.7 |
| *Pongo pygmaeus* | -34.9 | -34.4 | 73.9 | 76.8 | 0.59 | -35.4 | -34.2 | 76.8 | 78.5 | 0.32 | -1.7 |
| *Macaca cyclopis* | -46.1 | -46.1 | 96.2 | 100.2 | 1.00 | -42.3 | -42.3 | 90.6 | 94.6 | 1.00 | +5.6 |
| *Macaca fuscata* | -40.9 | -33.7 | 85.9 | 75.4 | $7.1 \times 10^{-4}$ | -37.2 | -33.6 | 80.3 | 77.2 | 0.03 | -1.8 |
| *Macaca mulatta* | -36.5 | -35.2 | 77.1 | 78.4 | 0.26 | -39.0 | -35.2 | 84.0 | 80.3 | 0.02 | -1.9 |
| *Macaca hecki* | -62.8 | -46.6 | 129.6 | 101.2 | $9.2 \times 10^{-8}$ | -57.9 | -48.3 | 121.8 | 106.5 | $6.5 \times 10^{-5}$ | -5.3 |
| *Macaca leonina* | -90.8 | -65.0 | 185.7 | 138.0 | $6.0 \times 10^{-12}$ | -59.3 | -55.7 | 124.6 | 121.5 | 0.03 | +16.5 |
| *Macaca maura* | -52.7 | -38.4 | 109.5 | 84.9 | $6.2 \times 10^{-7}$ | -45.1 | -38.4 | 96.2 | 86.7 | $1.2 \times 10^{-3}$ | -1.8 |
| *Macaca nemestrina* | -46.8 | -35.7 | 97.6 | 79.4 | $1.5 \times 10^{-5}$ | -45.0 | -35.9 | 96.0 | 81.8 | $1.2 \times 10^{-4}$ | -2.5 |
| *Macaca nigra* | -37.5 | -37.5 | 79.0 | 83.0 | 1.00 | -38.4 | -37.5 | 82.8 | 85.0 | 0.41 | -2.0 |
| *Macaca radiata* | -47.1 | -47.1 | 98.2 | 102.2 | 0.99 | -39.2 | -39.2 | 84.5 | 88.5 | 1.00 | +13.8 |
| *Macaca silenus* | -38.7 | -38.0 | 81.5 | 84.0 | 0.47 | -37.8 | -37.1 | 81.6 | 84.3 | 0.53 | -0.3 |
| *Macaca tonkeana* | -119.2 | -43.8 | 242.4 | 95.6 | $1.8 \times 10^{-33}$ | -78.5 | -45.3 | 162.9 | 100.6 | $4.0 \times 10^{-15}$ | -5.0 |
| *Macaca arctoides* | -37.8 | -37.8 | 79.5 | 83.5 | 1.00 | -37.0 | -36.7 | 80.0 | 83.5 | 0.79 | +0.0 |
| *Macaca assamensis* | -43.3 | -35.7 | 90.6 | 79.3 | $4.8 \times 10^{-4}$ | -44.4 | -35.7 | 94.9 | 81.4 | $1.6 \times 10^{-4}$ | -2.1 |
| *Macaca leucogenys* | -59.0 | -41.6 | 122.0 | 91.2 | $2.7 \times 10^{-8}$ | -57.5 | -43.1 | 121.1 | 96.2 | $5.4 \times 10^{-7}$ | -5.1 |
| *Papio anubis* | -58.2 | -45.2 | 120.4 | 98.3 | $2.2 \times 10^{-6}$ | -42.9 | -41.6 | 91.9 | 93.2 | 0.26 | +5.1 |
| *Papio cynocephalus anubishybrid* | -73.0 | -71.9 | 150.1 | 151.9 | 0.34 | -63.4 | -63.4 | 132.8 | 136.8 | 1.00 | +15.1 |
| *Papio cynocephalus ssp* | -59.7 | -59.6 | 123.3 | 127.3 | 0.98 | -47.0 | -47.0 | 100.0 | 104.0 | 1.00 | +23.3 |
| *Papio hamadryas* | -70.7 | -70.7 | 145.4 | 149.4 | 1.00 | -53.3 | -53.3 | 112.5 | 116.5 | 1.00 | +32.9 |
| *Papio kindae* | -50.3 | -50.3 | 104.5 | 108.5 | 1.00 | -40.8 | -40.8 | 87.6 | 91.6 | 1.00 | +16.9 |
| *Papio ursinus* | -80.7 | -78.7 | 165.4 | 165.5 | 0.14 | -64.6 | -64.6 | 135.2 | 139.2 | 1.00 | +26.3 |
| *Theropithecus gelada* | -47.6 | -47.6 | 99.2 | 103.2 | 1.00 | -36.6 | -36.6 | 79.3 | 83.3 | 1.00 | +19.9 |
| *Erythrocebus patas* | -61.9 | -55.2 | 127.9 | 118.4 | $1.2 \times 10^{-3}$ | -62.1 | -57.1 | 130.1 | 124.2 | $7.1 \times 10^{-3}$ | -5.8 |
| *Rhinopithecus bieti* | -88.9 | -65.2 | 181.7 | 138.4 | $5.2 \times 10^{-11}$ | -76.7 | -66.6 | 159.5 | 143.1 | $3.8 \times 10^{-5}$ | -4.8 |
| *Rhinopithecus roxellana* | -214.0 | -86.4 | 432.1 | 180.8 | $3.7 \times 10^{-56}$ | -152.2 | -86.6 | 310.4 | 183.2 | $3.2 \times 10^{-29}$ | -2.4 |
| *Semnopithecus hypoleucos* | -62.4 | -62.3 | 128.8 | 132.5 | 0.86 | -60.4 | -60.1 | 126.8 | 130.2 | 0.73 | +2.3 |
| *Trachypithecus francoisi* | -42.2 | -42.2 | 88.4 | 92.4 | 1.00 | -38.6 | -38.6 | 83.1 | 87.1 | 1.00 | +5.3 |
| *Trachypithecus poliocephalus* | -35.6 | -35.6 | 75.2 | 79.2 | 1.00 | -33.7 | -33.7 | 73.4 | 77.4 | 1.00 | +1.8 |
| **Mean** | -55.1 | -45.7 | 114.2 | 99.4 | — | -47.9 | -42.7 | 101.7 | 95.4 | — | +4.0 |

**Table B6:** Regression of the fraction of the DFE with *S* ∈ (−∞, −10] on log_10_ *N_e_* and generation time, under the discrete DFE allowing for beneficial mutations. Coefficients and their associated *p*-values are reported for ordinary least squares (OLS) and phylogenetic generalized least squares (PGLS) regressions. For the Grafen PGLS model, *ρ* is the branch-length scaling parameter inferred from the data. Tables B7–B9 report the same quantities for the remaining three DFE-model variants.

| Model | AIC | $R^2$ | Intercept | p(Int) | $\log_{10} N_e$ | $p(\log_{10} N_e)$ | gen_time | p(gen_time) |
| --- | --- | --- | --- | --- | --- | --- | --- | --- |
| OLS | -108.50 | 0.735 | 0.022 | 0.881 | 0.126 | $8.18 \times 10^{-5}$ | -0.003 | 0.084 |
| PGLS Brownian | -108.58 | 0.724 | -0.129 | 0.28 | 0.153 | $4.31 \times 10^{-8}$ | -0.001 | 0.63 |
| PGLS Grafen ($\rho = 0.21$) | -107.09 | 0.727 | -0.128 | 0.378 | 0.154 | $3.37 \times 10^{-6}$ | -0.002 | 0.457 |

**Table B7:**
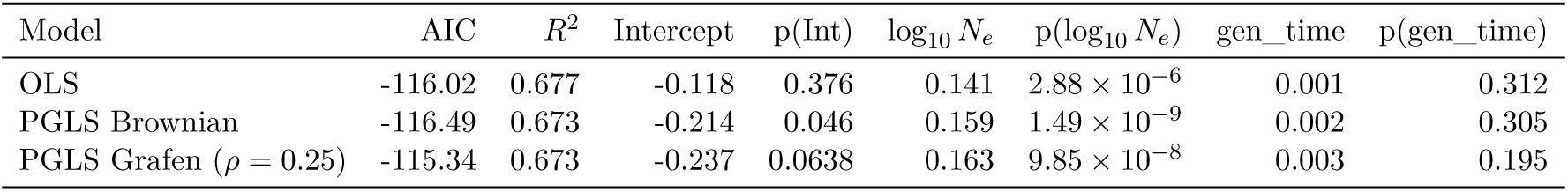
As Table B6, but under the discrete DFE with deleterious mutations only.

**Table B8:** As Table B6, but under the gamma–exponential DFE allowing for beneficial mutations.

| Model | AIC | $R^2$ | Intercept | p(Int) | $\log_{10} N_e$ | $p(\log_{10} N_e)$ | gen_time | p(gen_time) |
| --- | --- | --- | --- | --- | --- | --- | --- | --- |
| OLS | -117.44 | 0.715 | 0.060 | 0.643 | 0.117 | $3.09 \times 10^{-5}$ | -0.001 | 0.33 |
| PGLS Brownian | -115.02 | 0.707 | -0.077 | 0.469 | 0.142 | $2.17 \times 10^{-8}$ | -0.000 | 0.955 |
| PGLS Grafen ($\rho = 0.18$) | -115.97 | 0.709 | -0.049 | 0.698 | 0.137 | $2.38 \times 10^{-6}$ | -0.000 | 0.838 |

**Table B9:** As Table B6, but under the gamma DFE with deleterious mutations only.

| Model | AIC | $R^2$ | Intercept | p(Int) | $\log_{10} N_e$ | $p(\log_{10} N_e)$ | gen_time | p(gen_time) |
| --- | --- | --- | --- | --- | --- | --- | --- | --- |
| OLS | -125.42 | 0.747 | 0.004 | 0.973 | 0.123 | $2.12 \times 10^{-6}$ | -0.000 | 0.795 |
| PGLS Brownian | -117.03 | 0.738 | -0.071 | 0.491 | 0.134 | $3.27 \times 10^{-8}$ | 0.001 | 0.712 |
| PGLS Grafen ($\rho = 0.12$) | -124.01 | 0.744 | -0.047 | 0.674 | 0.131 | $6.30 \times 10^{-7}$ | 0.000 | 0.862 |

## C Simulation Results

We performed additional simulations to quantify the impact of demographic distortions and SFS sample size on inference accuracy. Our previous work on fastDFE (Sendrowski and Bataillon, 2024) includes benchmarks against polyDFE (Tataru et al., 2017), which was at the time validated using the forward simulator SFS_CODE (Hernandez, 2008). Those simulations covered a range of DFEs and demographic histories, including moderate population growth and decline, as well as exponential changes to twice or half the original population size. Here, we extend this validation using SLiM (Haller and Messer, 2023) and assess inference performance under more pronounced demographic scenarios that induce stronger distortions in the SFS. We also evaluate robustness to common model violations, including population substructure and dominance effects. Finally, inference accuracy is examined as a function of SFS sample size and the number of available SNPs (Andersson et al., 2023).

### Simulation setup

We modeled a Wright–Fisher population of 1000 individuals, which is large enough to limit strong genetic drift while remaining computationally tractable. Larger population sizes were avoided because they substantially increase both runtime per generation and the number of generations required to reach equilibrium. Each simulation was run for 10,000 generations to ensure equilibrium. We simulated 10^9^ sites under uniform mutation and recombination rates of 10^−8^ and 10^−7^ per site, respectively, unless stated otherwise. Both rates are broadly consistent with estimates in humans. The elevated recombination rate was chosen to reduce variance introduced by background selection, which is not explicitly modeled in fastDFE. To assess its potential impact, we additionally performed simulations with stronger background selection. To enable parallelization, simulations were divided into 100 chunks of 10^7^ sites each, which were subsequently merged.

Mutations were assigned selection coefficients drawn from a specified DFE, assuming semidominance (additive effects) unless stated otherwise. We used the default GammaExpParametrization, which models deleterious effects with a gamma distribution and beneficial effects with an exponential distribution. As defined above, *S_d_* and *S_b_*denote the mean population-scaled selection coefficients of deleterious and beneficial mutations, respectively, *b* is the gamma shape parameter controlling dispersion, and *p_b_* is the fraction of beneficial mutations. fastDFE defines selection coefficients such that heterozygous and homozygous genotypes have fitnesses 1 + 2*hs* and 1 + 2*s*, respectively. In contrast, SLiM uses 1 + *hs* and 1 + *s*. We therefore follow fastDFE’s convention and scale selection coefficients by a factor of two when using SLiM. Parameters *S_d_* and *S_b_*are scaled by the effective population size (*S* = 4*N_e_s*), which is not known a priori because it depends on both the DFE and the demographic history. We therefore estimate *N_e_*post hoc using Watterson’s estimator, consistent with the procedure implemented in fastDFE, and scale the simulated selection coefficients accordingly when comparing results. All sites were assumed to be coding and thus eligible for mutation affecting fitness. Neutral mutations were subsequently added to the generated tree sequences using msprime (Baumdicker et al., 2022), as this is computationally more efficient than simulating them forward in time.

Site-frequency spectra were computed using a hypergeometric sampling scheme, drawing without replacement from the full population, which reduces sampling variance relative to random subsampling. By default, we subsampled 20 individuals independently at each site, and additionally considered other sample sizes to assess their effect on inference accuracy. Finally, fastDFE was applied to recover the DFE jointly from the simulated *selected* SFS, shaped by both demography and selection, and the *neutral* SFS, shaped by demography alone.

### Recovery under a constant population size

We evaluated recoverability for four DFEs: (1) a strongly deleterious DFE (*s_d_* = −0.3, *b* = 0.3), (2) a weakly deleterious DFE (*s_d_* = −0.03, *b* = 0.1), (3) a strongly deleterious DFE with a small fraction of beneficial mutations (*s_d_* = −0.3, *b* = 0.1, *p_b_* = 0.01, *s_b_* = 0.01), and (4) a weakly deleterious DFE with a larger fraction of mildly beneficial mutations (*s_d_* = −0.03, *b* = 0.3, *p_b_* = 0.05, *s_b_* = 0.001). Figure C1 shows the results under a constant population size. The inferred DFEs closely match the simulated ones. Greater variance is observed for beneficial mutations, particularly near selective neutrality (|*S*| *<* 1), where slightly deleterious and beneficial mutations are difficult to distinguish (not shown). This increase in uncertainty is expected, as the full DFE has four parameters rather than two, substantially increasing the degrees of freedom. Beneficial mutations also leave a weaker signal in the data, as they tend to be rarer and of smaller effect. Inference based on folded SFS performs well overall, but shows reduced accuracy for beneficial mutations, as folding removes the excess of high-frequency derived alleles, which is a key signal of positive selection (Figure C6).

**Figure C1:**
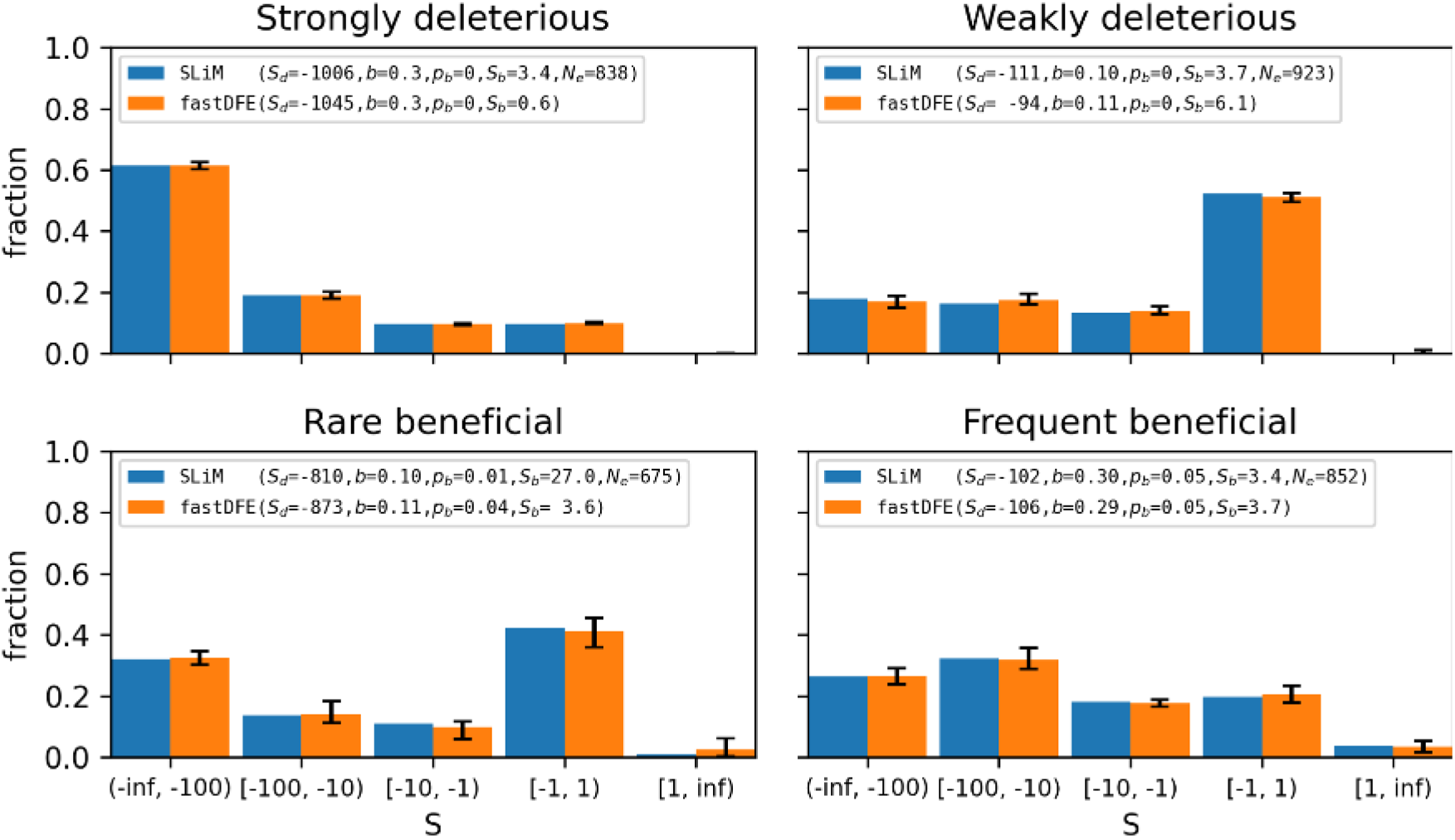
DFEs simulated with SLiM and inferred with fastDFE under a constant Wright–Fisher population of 1000 individuals. Legends report the simulated and inferred parameters of the GammaExpParametrization model. Bars represent 90% confidence intervals, and point estimates correspond to mean values across bootstrap replicates.

### Demographic scenarios and model violations

We explored a range of demographic scenarios, including population expansion, reduction, bottlenecks, substructure, and the effects of recessive mutations. Hyperparameters for these scenarios were chosen to induce relatively pronounced distortions in the SFS (Figure C2). In the expansion scenario, population size instantaneously increased fourfold from 1000 to 4000 individuals 500 generations before the end of the simulation. Similarly, in the reduction scenario, population size instantaneously decreased fourfold from 1000 to 250 individuals 500 generations before the end. In the bottleneck scenario, population size dropped from 1000 to 50 individuals 600 generations before the end of the simulation and recovered to 1000 individuals 100 generations later. In the substructure scenario, two sub-populations of 500 individuals exchanged migrants at a rate of *m* = 0.0001 per generation, giving rise to an *F_ST_* of approximately 0.3 to 0.45 depending on the underlying DFE. All these scenarios produce a non-equilibrium SFS and represent demographic, non-selective processes that are reflected in the *neutral* SFS, but may interact with selection in complex ways. In addition, we tested the impact of recessive mutations by simulating DFEs with a recessive dominance coefficient of *h* = 0.2 and inferring the DFE under the assumption of semidominance (*h* = 0.5). Figure C2 shows SFS simulated with both SLiM and fastDFE under these scenarios for a purely deleterious DFE with *s_d_* = −0.3 and *b* = 0.3. Selection coefficients were rescaled with the *N_e_* estimated from the *neutral* SFS, as described above. SLiM explicitly models the full evolutionary process, whereas fastDFE relies on the diffusion approximation and accounts for demography using nuisance parameters inferred from the *neutral* SFS. This correction corresponds to multiplicatively rescaling the expected *selected* SFS entries by the deviation of the corresponding *neutral* SFS entries from the constant-size expectation. Despite substantial demographic distortions, the *selected* SFS is predicted accurately by fastDFE. Without this correction, the shape of the SFS simulated with fastDFE would be indistinguishable from that of the constant-size scenario across all demographic models. Large deviations occur in the recessive case, as dominance effects are not reflected in the *neutral* SFS.

**Figure C2:**
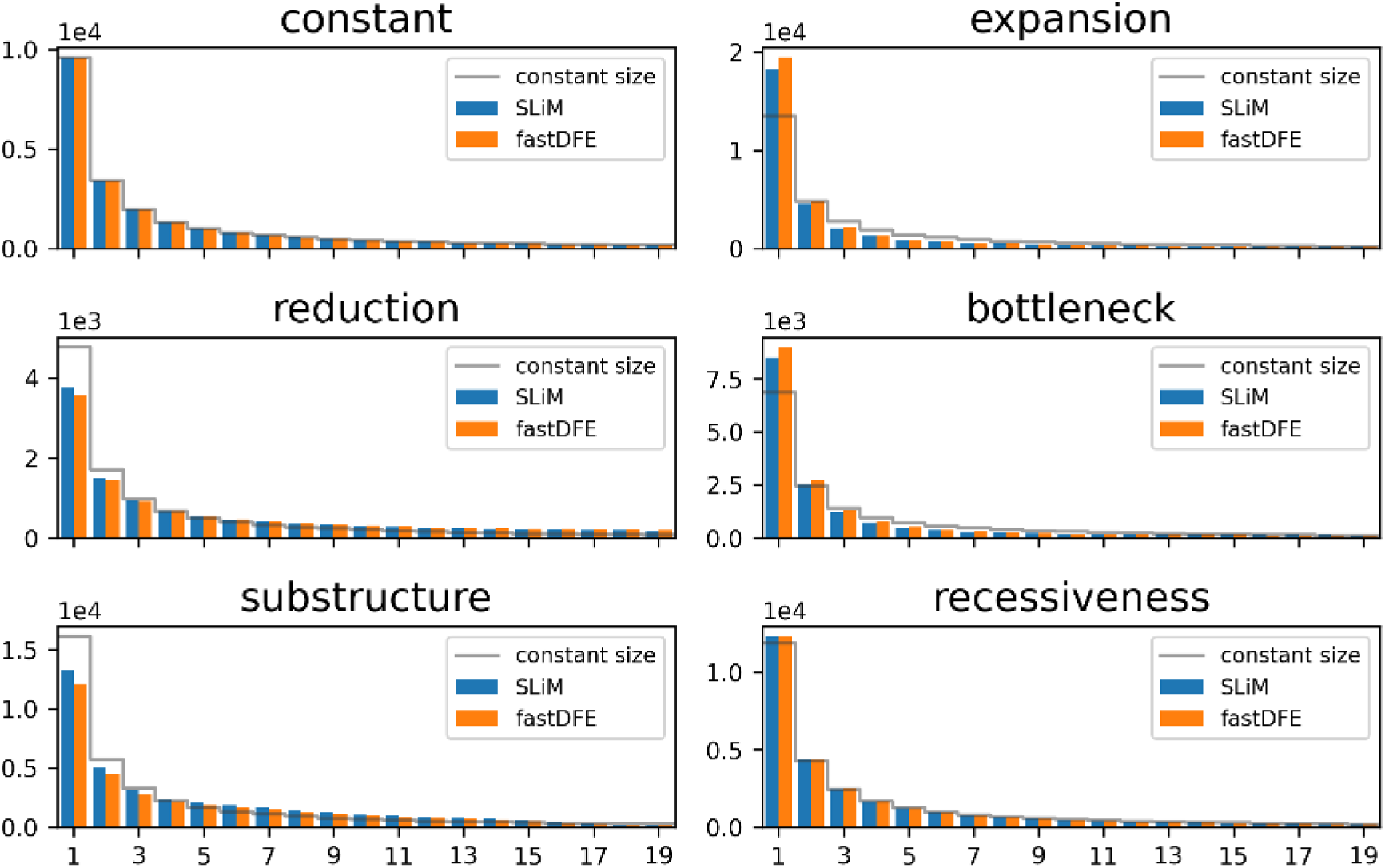
Site-frequency spectra simulated with SLiM and fastDFE using nuisance parameters to correct for demography, shown for *selected* sites. The constant-size spectrum is repeated in every panel as a reference, rescaled to the total of that panel so that it marks the expected shape rather than a count. The underlying DFE is purely deleterious with *s_d_*= −0.3 and *b* = 0.3 (cf. strongly deleterious DFE in Figure C1). Population expansion results in an excess of low-frequency derived variants. Population reduction produces a relative deficit of low-frequency derived variants. Bottlenecks additionally reduce overall diversity, particularly at intermediate and high frequencies, since new mutations at low frequencies have had time to accumulate after the bottleneck. The two-split substructure scenario produces irregular spectra at intermediate frequencies, but these effects can be corrected with good accuracy using the *neutral* SFS. Making mutations recessive (*h* = 0.2) generates a relative excess of low-frequency derived variants due to masking of deleterious alleles in heterozygotes. In the constant-size panel the two spectra are identical, aside from variance due to genetic drift in the SLiM results.

Next, we assessed how well fastDFE recovers the ground-truth DFE under these demographic scenarios. Overall, DFEs can be recovered with good accuracy. Population expansion leads to a slight bias toward more deleterious DFEs (Figure C7), whereas population reduction results in DFEs that are consistently inferred as less deleterious than the true ones (Figure C8). The bottleneck scenario also induces a bias toward more deleterious DFEs, although this effect is weaker than in the expansion case (Figure C9). Uncertainty in DFE estimates is largest for the population reduction and bottleneck scenarios, particularly when inferring the full DFE. This likely reflects the reduced genetic diversity in these scenarios and the consequently smaller number of SNPs. In the population structure scenario, DFEs are recovered accurately, with only a slight bias toward less deleterious effects (Figure C10). When mutations are simulated as recessive (*h* = 0.2) but inferred assuming semidominance (*h* = 0.5), a clear bias toward less deleterious DFEs is observed, as recessive mutations are interpreted as being under weaker selection. In contrast, the DFE is recovered well when the correct dominance coefficient is used during inference (Figure C11). Whether the inferred DFE is biased toward more or less deleterious effects is clearly reflected in the relative deficit or excess of singletons apparent in Figure C2 (Figure C5). Masking singletons from the SFS, however, yielded essentially identical mean DFE estimates, albeit with wider confidence intervals (not shown).

Even when parameter estimates deviate from the true values, the overall shape of the DFE, as reflected by the distribution of probability mass across the discretization bins, is often well captured. In some cases, we observe an inverse relationship between the mean strength of deleterious selection and the gamma shape parameter (*b* ∼ |*S_d_*|^−1^). Similarly, the inferred fraction of beneficial mutations tends to decrease as their mean strength increases (*p_b_*∼*S_b_*^-1^). These patterns indicate mild parameter non-identifiability in the GammaExpParametrization, whereby moderate numbers of strongly selected mutations can compensate for larger numbers of more weakly selected mutations. Nevertheless, the shape parameter *b* of the DFE for deleterious mutations is often estimated accurately, even when mean *S_d_* itself is biased.

Figure C3 shows simulations in which a purely deleterious DFE with *S_d_* = −300 and *b* = 0.3 was simulated under fixed dominance coefficients *h* ∈ [0, 1] with fastDFE, and both the DFE parameters and *h* were jointly inferred from an SFS of sample size 20. Mean estimates of *h* broadly track the true values, but uncertainty remains substantial, decreasing only moderately with larger sample sizes (not shown). Joint bootstrap distributions of (*h, S_d_*) reveal a strong negative covariance between *h* and *S_d_*, indicating partial non-identifiability whereby stronger selection can be compensated by increased recessiveness. Overall, SFS data can constrain plausible dominance regimes, but accurate inference of *h* remains difficult from the SFS alone, and more so in realistic settings with additional sources of noise.

**Figure C3:**
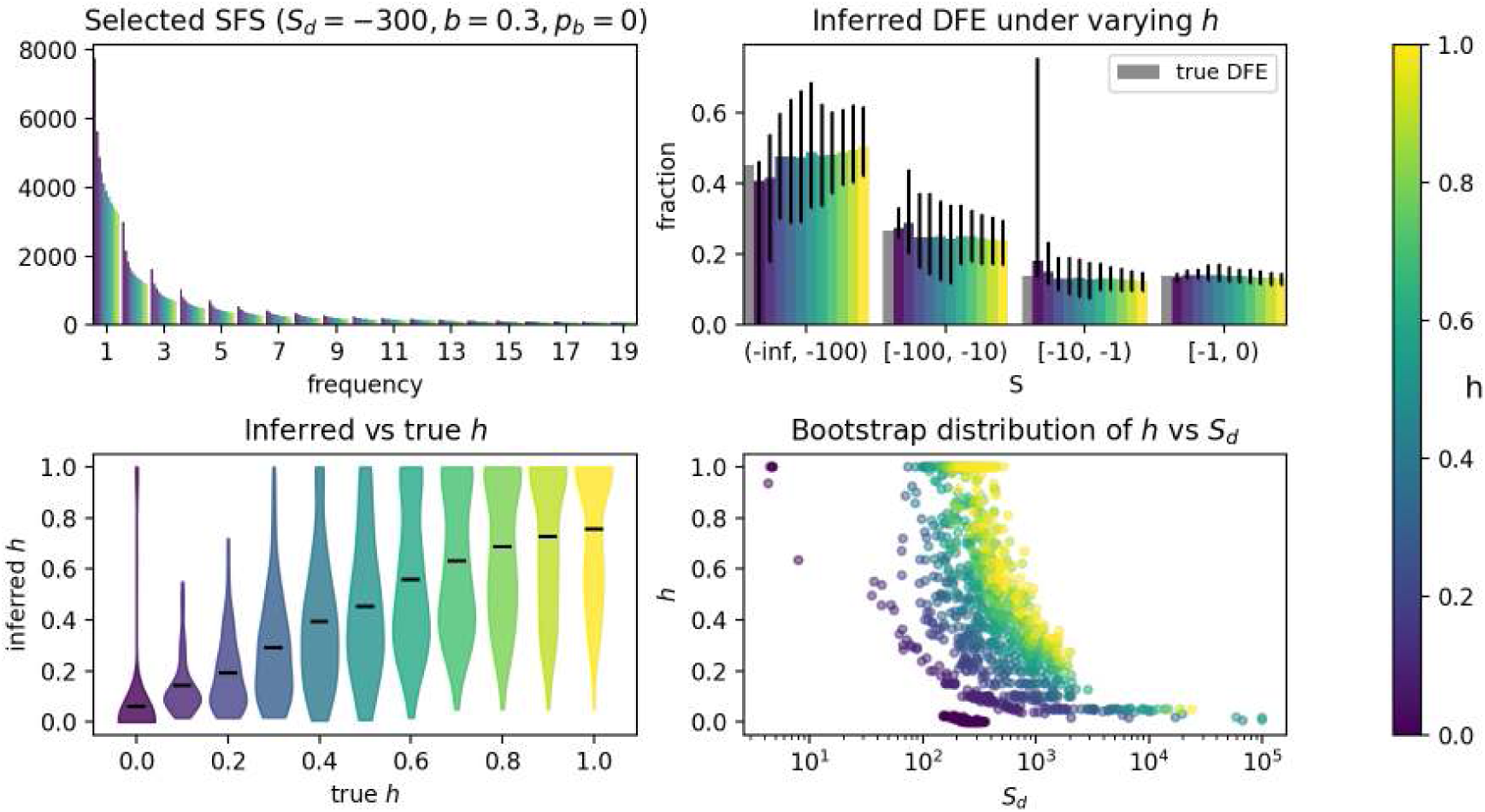
Inference of dominance coefficients from simulated SFS data. Each dataset was generated under a fixed purely deleterious DFE (*S_d_* = −300, *b* = 0.3, *p_b_* = 0) with different dominance coefficients *h* ∈ [0, 1]. Top left: Simulated SFS. Top right: True (gray) and inferred DFEs. Bars indicate 90% confidence intervals and point estimates correspond to mean values across bootstrap replicates. Uncertainty is substantial, with a mild bias toward more deleterious DFEs for more dominant mutations, but mean bootstrap estimates remain broadly consistent across *h*. Bottom left: Violin plot comparing the inferred *h*to its true value. Estimates vary considerably, although mean values remain close to the truth except under strong dominance. Bottom right: Joint bootstrap distribution of *S_d_* and *h*, illustrating an inverse relationship in which stronger selection can be compensated by increased recessiveness (100 bootstrap replicates).

### Inferring dominance from SFS data

As shown above, assuming semidominance when mutations are in fact recessive biases inference toward less deleterious DFEs. We also expect that estimating the dominance coefficient itself, particularly its relationship with selection strength, will be challenging: previous attempts to fit SFS data while relaxing the assumption of *h* = 0.5 were compatible with a wide range of dominance models (Kyriazis and Lohmueller, 2024). We therefore performed simulations to quantify how well *h* can be inferred from moderately sized SFS data sets when jointly fitting a DFE and *h*.

### Sources of residual discrepancy

Beyond demographic effects, two main sources of variation contribute to discrepancies between the ground truth and inferred DFEs. First, genetic drift remains non-negligible at *N* = 1000, although its impact is limited (Figure C12). Second, background selection induces linkage effects that are not modeled by fastDFE, which assumes site independence. We partially mitigated these effects by using a relatively high recombination rate, and deviations indeed become more pronounced under stronger selection or lower recombination (Figure C13). Because background selection also affects the *neutral* SFS, some degree of correction is expected through the nuisance parameters. Increasing the sample size and the number of SNPs beyond those used here would further improve inference accuracy (Figure C14).

### Data requirements

DFE inference accuracy depends on the amount of data available, namely the number of SNPs and the number of sampled individuals. We disentangle these two factors by simulating SFS data with fastDFE’s simulation module, which varies them independently. Figure C4 shows results for two DFEs: one purely deleterious and one including beneficial mutations. In one setup, we fix the total number of SNPs to 2 × 10^4^ and vary the SFS sample size from 6 to 100 individuals. *Neutral* and *selected* spectra were matched to contain the same number of SNPs, representing an idealized setting, as real data typically contain more *selected* than *neutral* SNPs. In a second setup, we fix the SFS sample size to 20 individuals and vary the total number of SNPs from 3 × 10^3^ to 3 × 10^6^. Spectra were simulated without demographic distortions or sampling noise, but Poisson resampling during bootstrapping quantifies uncertainty due to finite SNP counts. Estimation of the DFE for deleterious mutations (*S_d_* = −300, *b* = 0.3) is robust to sample size and SNP count, performing well even at *n*= 6 and a few thousand SNPs. In contrast, accurate and stable inference of the full DFE (*S_d_* = −300, *b* = 0.3, *p_b_* = 0.05, *S_b_* = 1) requires larger sample sizes and substantially more SNPs, typically *n* ≥ 20 and #SNP ≥ 10^4^. Disentangling weakly deleterious (−1 ≤ *S* ≤ 0) from weakly beneficial (0 ≤ *S* ≤ 1) mutations requires even larger numbers of SNPs. Increasing the SFS sample size at a fixed number of SNPs yields diminishing returns. When SNP counts are low, larger subsample sizes result in higher sampling variance across SFS bins, leading to less accurate DFE estimates. However, with the subsample size held fixed, increasing the number of SNPs improves inference accuracy, as Poisson resampling during bootstrapping becomes less variable.

**Figure C4:**
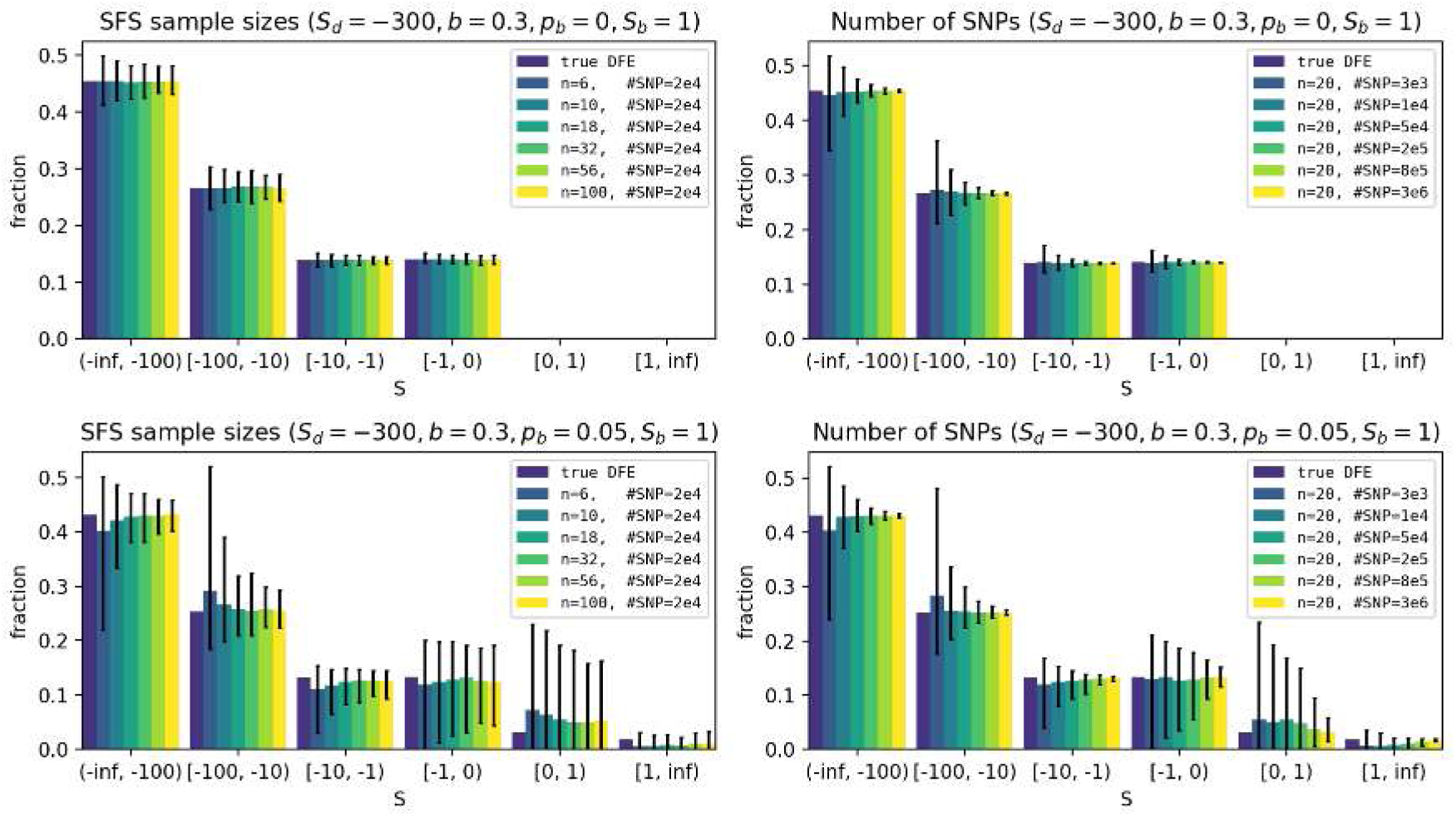
Ground truth and inferred DFEs under varying SFS sample sizes (left) and SNP counts (right). DFEs were generated using fastDFE’s simulation module without demographic distortions or sampling noise. Results are shown for a purely deleterious two-parameter DFE (top) and a four-parameter DFE including beneficial mutations (bottom). The DFE for deleterious mutations is recovered accurately even for small sample sizes and low SNP counts. Recovering the beneficial DFE requires larger sample sizes and substantially more SNPs. For a fixed number of SNPs, increasing the SFS sample size improves accuracy only up to a point. Bars indicate 90% confidence intervals.

### Summary

Overall, these results demonstrate that DFEs can be recovered with reasonable accuracy using nuisance parameters, even under substantial demographic distortions. How the compounded effects of many slight model violations affect inference in real data remains an open question. The dominance coefficient is difficult to infer accurately from SFS data alone, but can be partially constrained in idealized settings, despite substantial nonidentifiability with *S_d_*. Finally, inference accuracy increases with the amount of available data, with larger sample sizes and SNP counts being particularly important for inferring the full DFE.

## Additional figures

**Figure C5:**
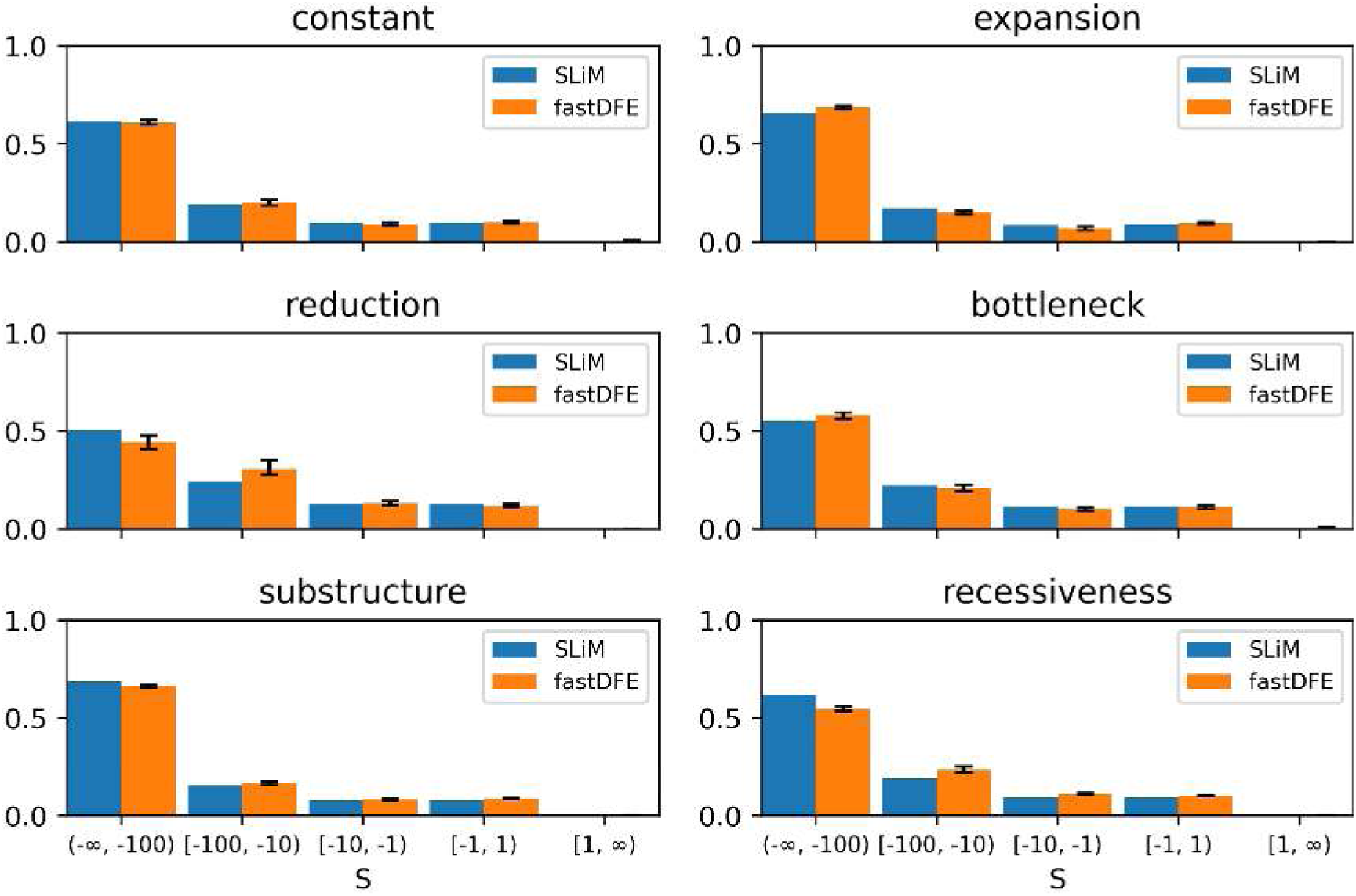
Ground truth and inferred DFEs under the different demographic scenarios in Figure C2, where SLiM provides the ground truth and fastDFE the inferred estimates. The underlying DFE is purely deleterious with *s_d_* = −0.3 and *b* = 0.3 (cf. strongly deleterious DFE in Figure C1).

**Figure C6:**
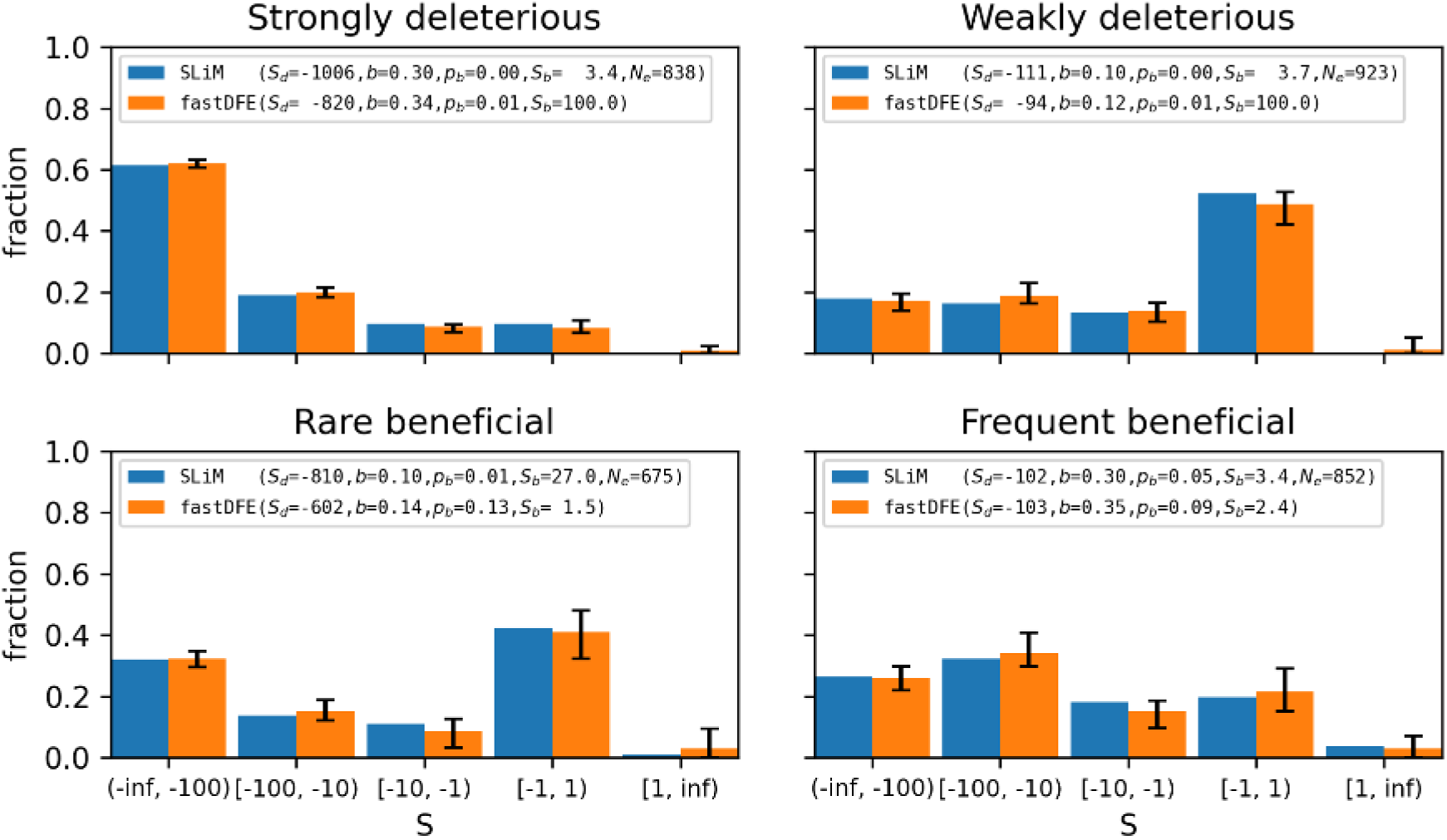
Ground truth and inferred DFEs under the constant population size scenario using a folded SFS (cf. Figure C1, which uses an unfolded SFS). Estimates are less accurate, particularly for beneficial mutations.

**Figure C7:**
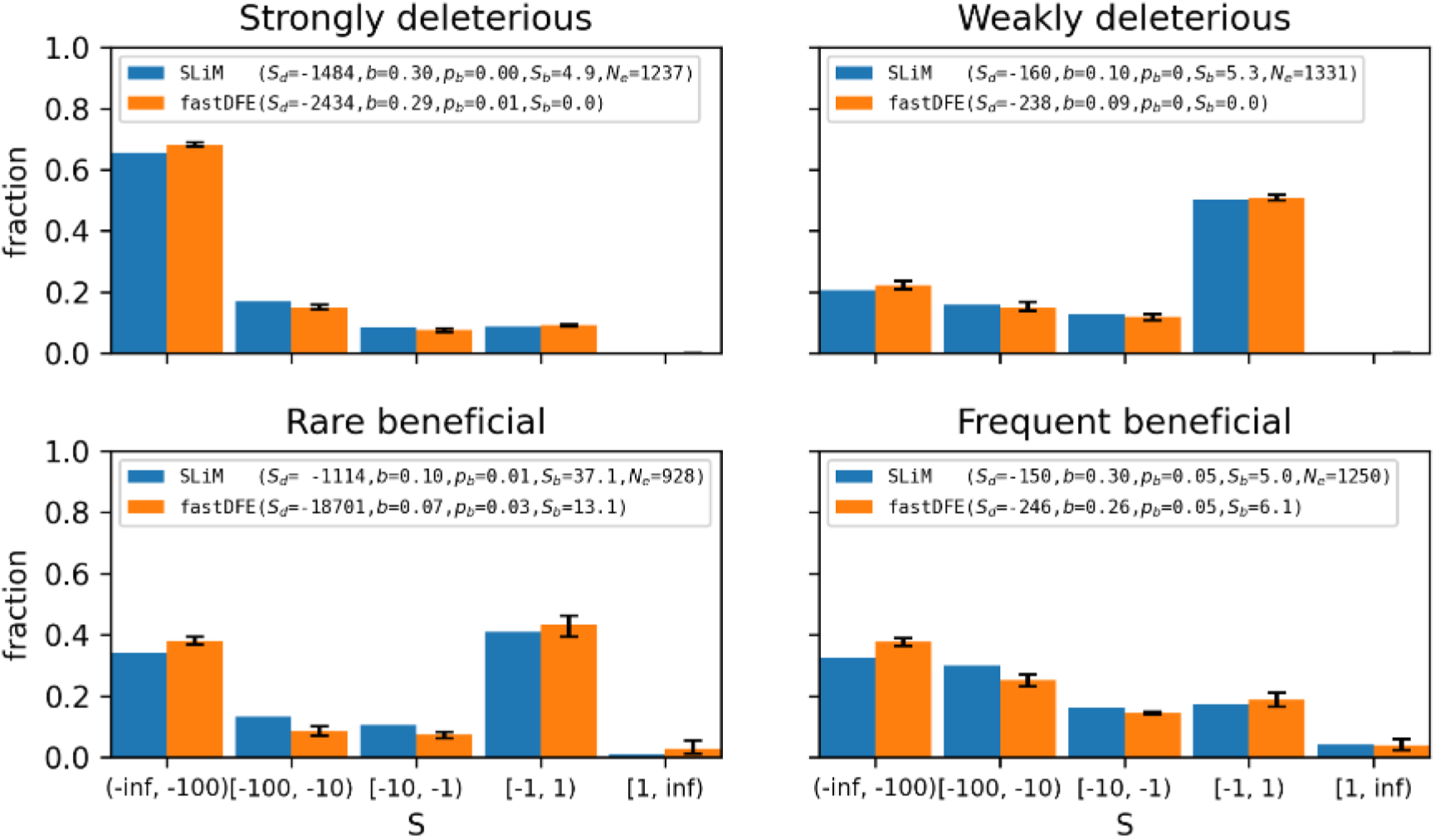
Ground truth and inferred DFEs under a population expansion scenario. The population size increases fourfold from 1000 to 4000 individuals 500 generations before the end of the simulation.

**Figure C8:**
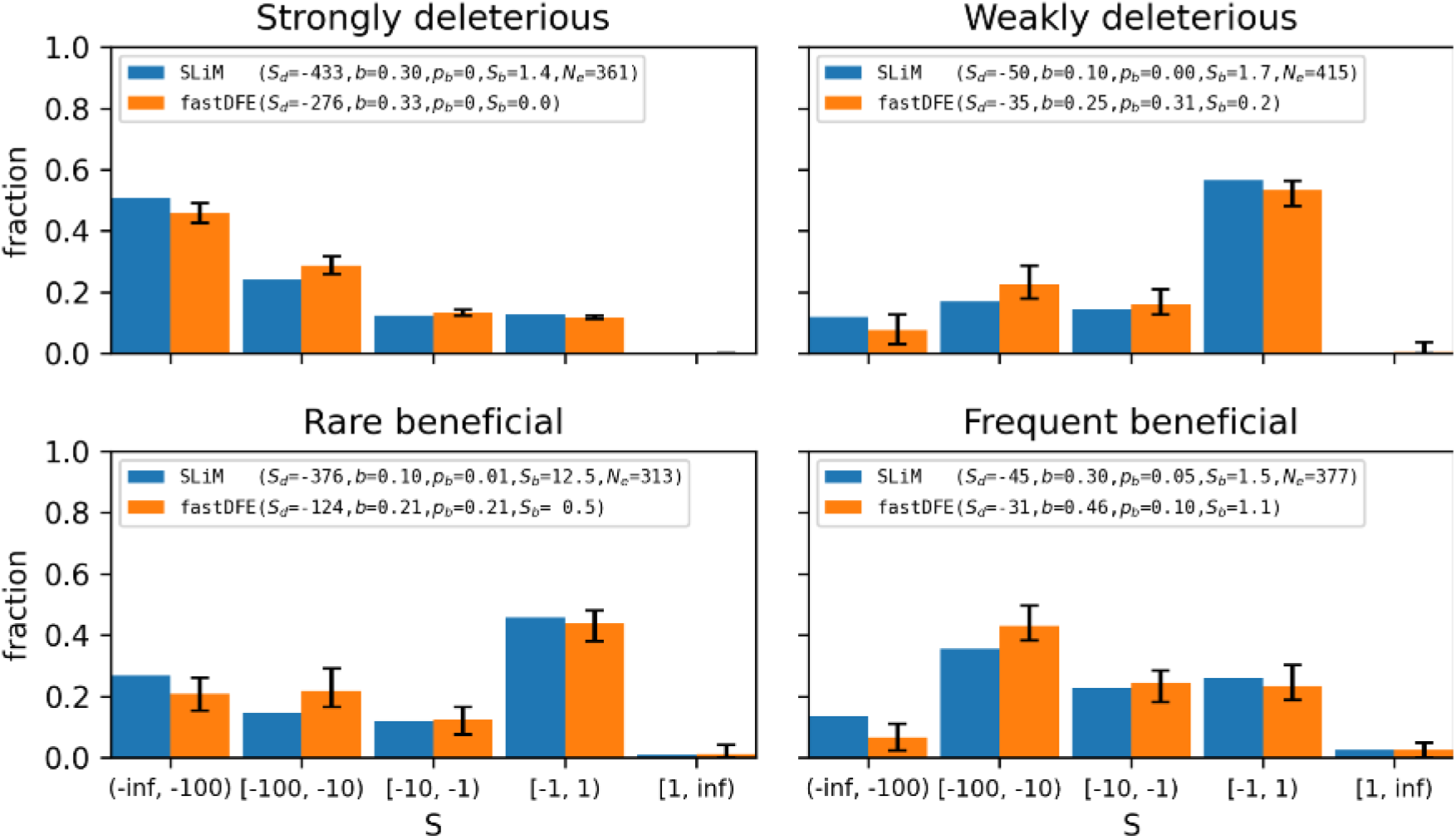
Ground truth and inferred DFEs under a population reduction scenario. The population size decreases fourfold from 1000 to 250 individuals 500 generations before the end of the simulation.

**Figure C9:**
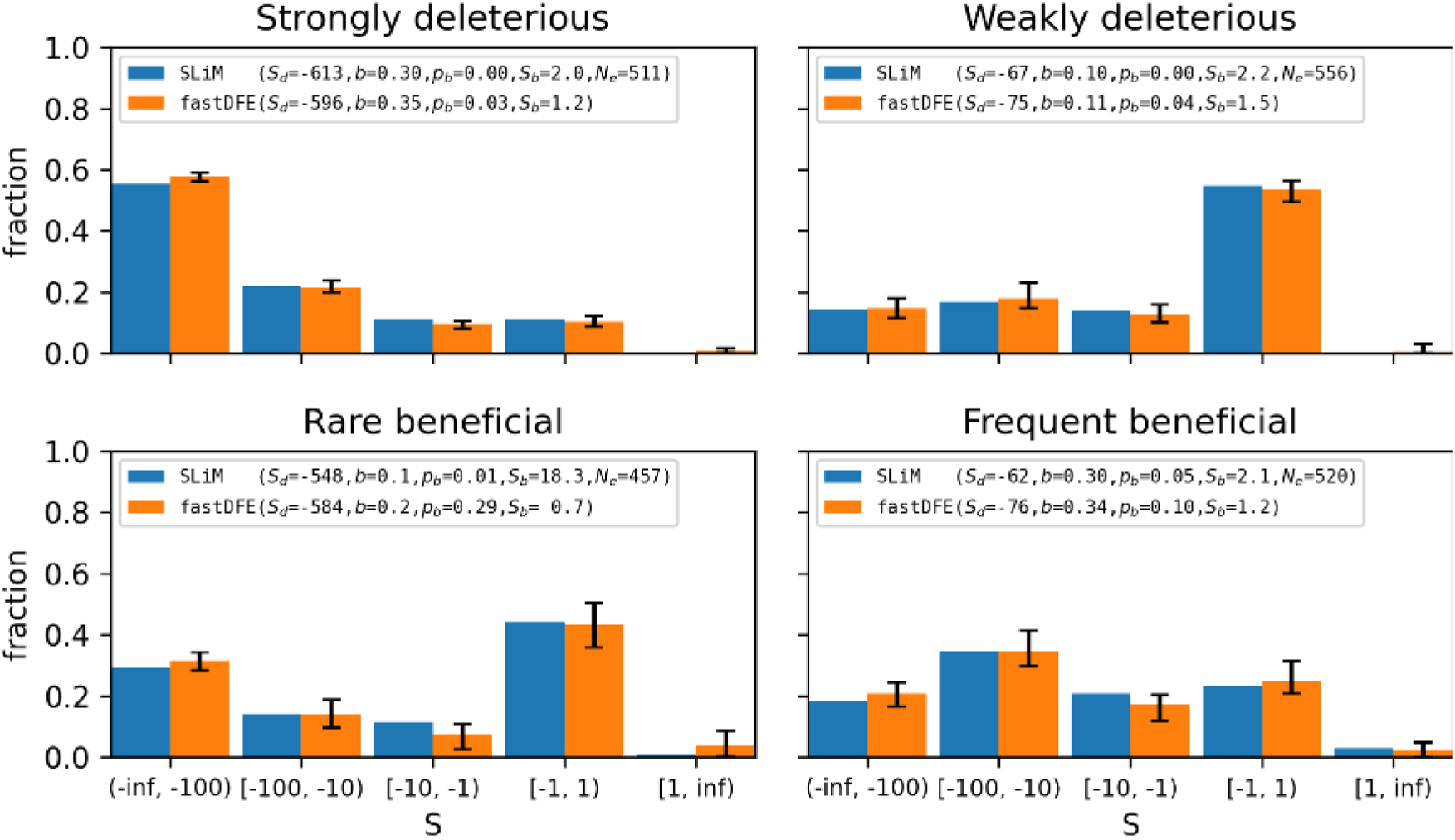
Ground truth and inferred DFEs under a bottleneck scenario. The population size is reduced from 1000 to 50 individuals 600 generations before the end of the simulation, then recovers to 1000 individuals 500 generations before the end.

**Figure C10:**
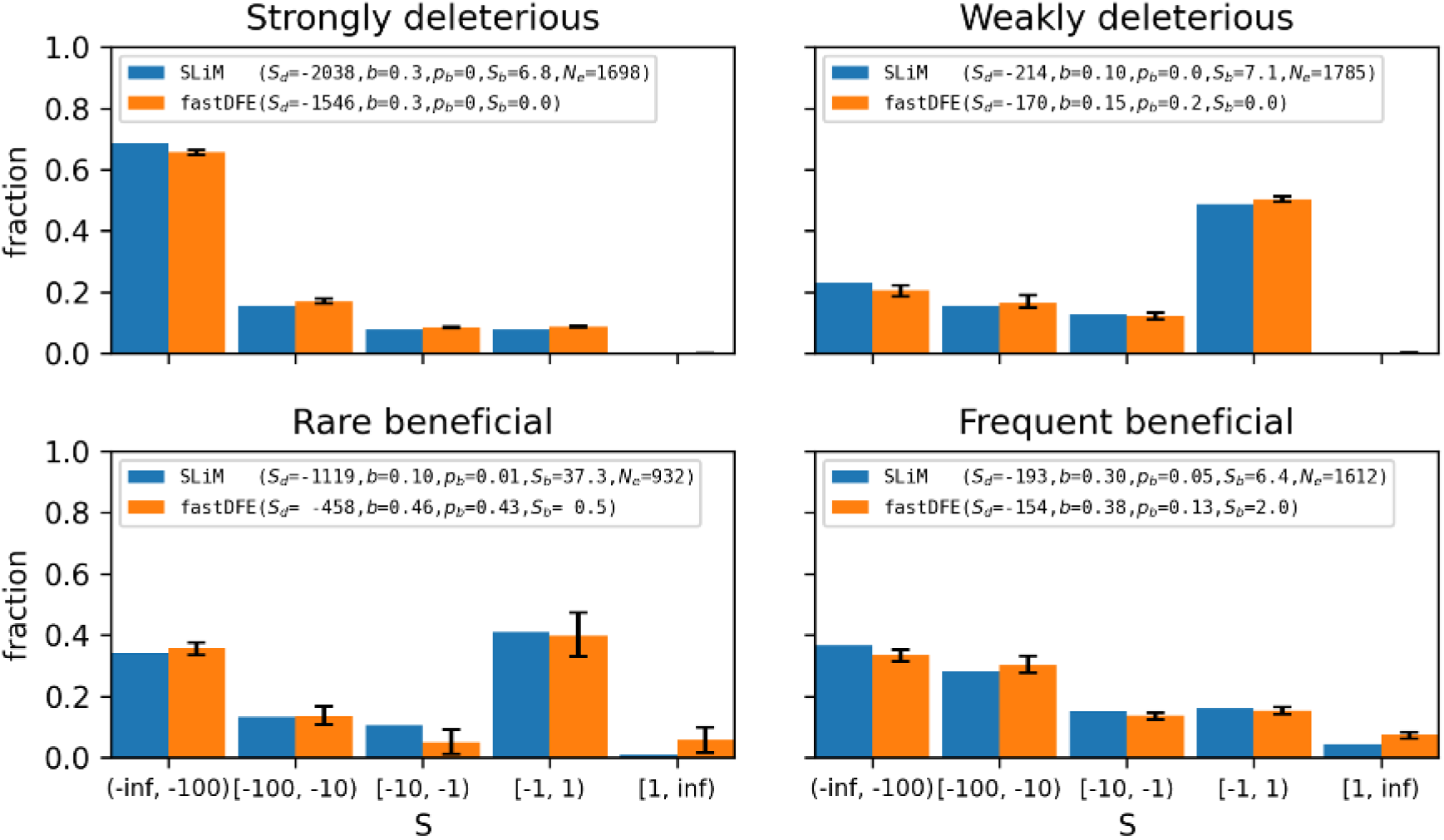
Ground truth and inferred DFEs under a population with sub-structure. Two subpopulations of 500 individuals each exchange migrants at a symmetric rate of *m* = 0.0001 per generation, resulting in an *F_ST_* of approximately 0.3 to 0.45 depending on the underlying DFE.

**Figure C11:**
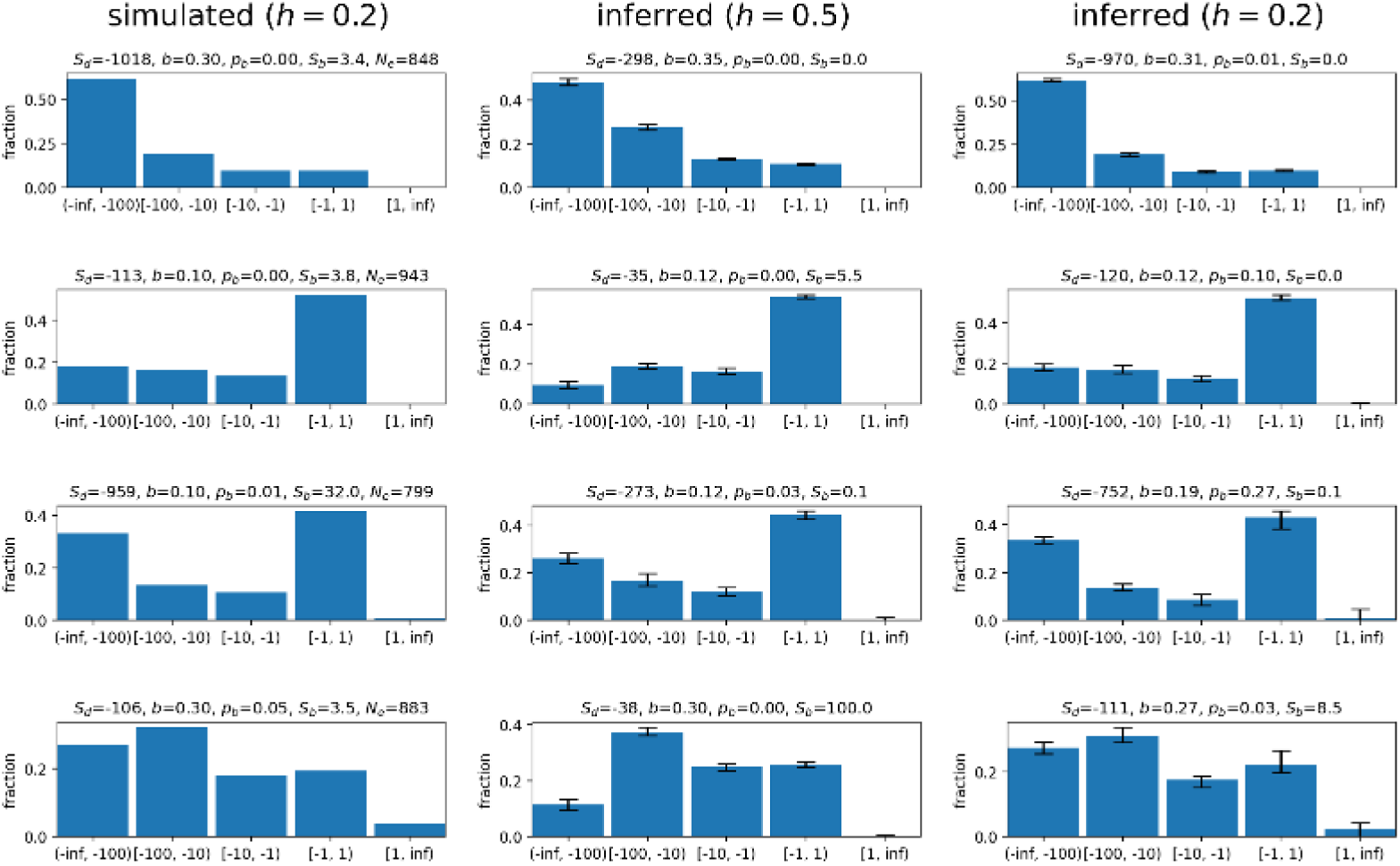
Left: Simulated DFEs with *h* = 0.2, corresponding to partially recessive mutations. **Middle:** Inferred DFEs assuming additivity (*h* = 0.5). **Right:** DFEs inferred using the correct dominance coefficient (*h* = 0.2). Assuming additivity for partially recessive mutations leads to underestimation of selection strength.

**Figure C12:**
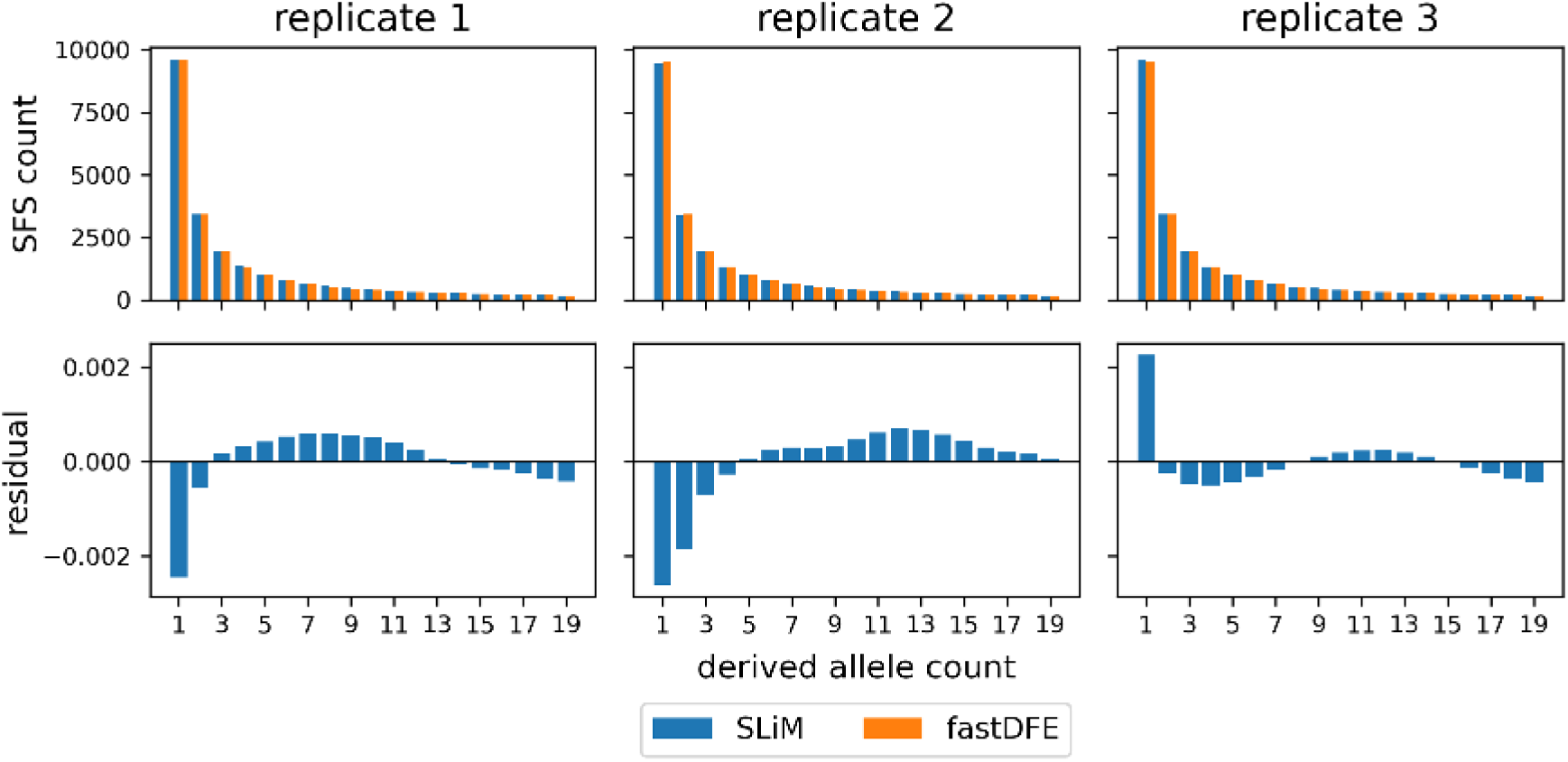
Three replicates of simulated spectra under the constant population size scenario. The underlying DFE is purely deleterious, with parameters *s_d_* = −0.3 and *b* = 0.3 (cf. strongly deleterious DFE in Figure C1). **Top:** SLiM and fastDFE spectra as counts. **Bottom:** the residual, defined as the SLiM minus the fastDFE proportion in each bin after normalizing both spectra to sum to one over their polymorphic bins. Deviations remain below 0.25% in every bin, and their sign varies across replicates and across frequency bins, reflecting drift.

**Figure C13:**
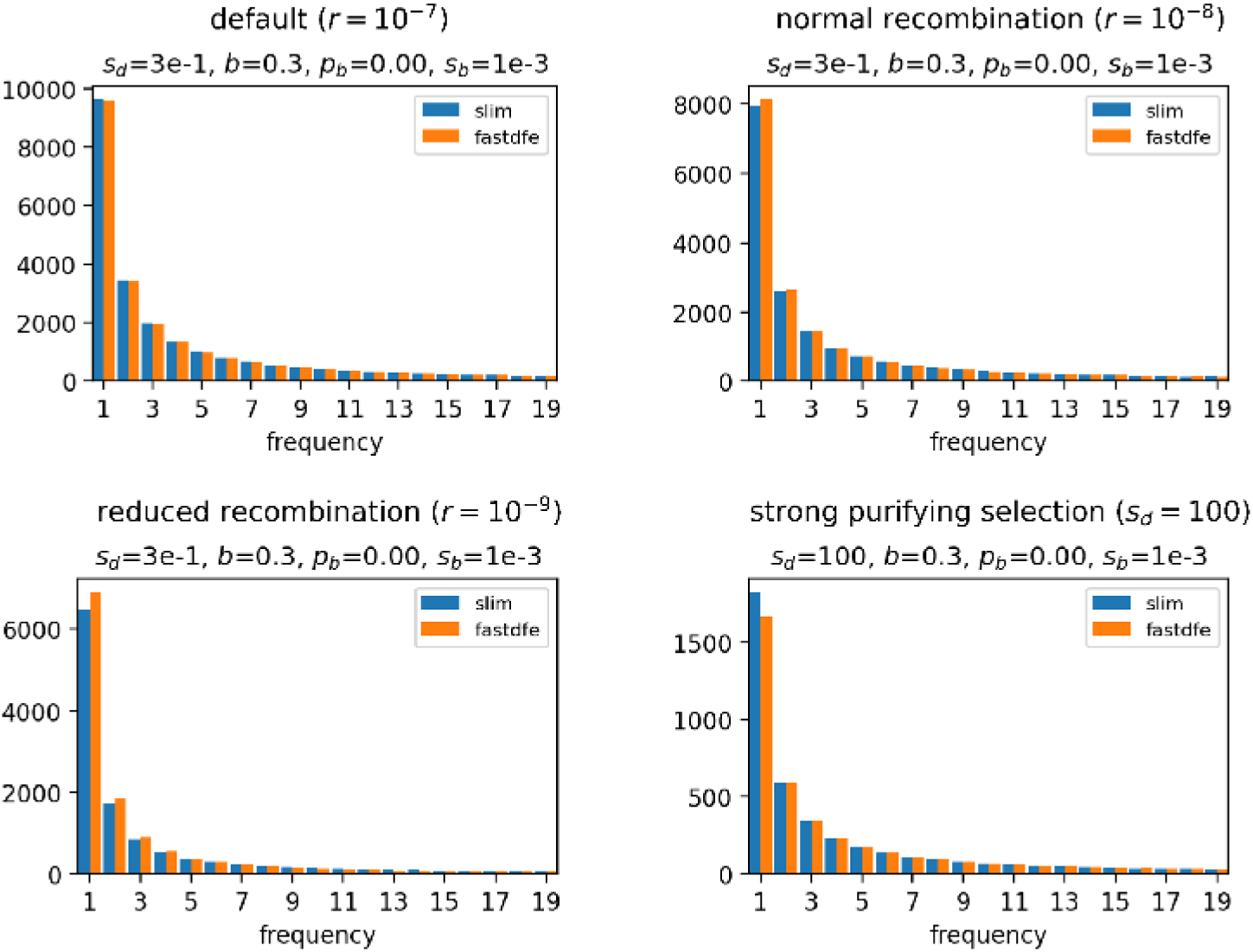
SFS comparison under varying recombination rates and strengths of negative selection. fastDFE assumes site independence and therefore does not account for background selection. Lower recombination and stronger deleterious effects increase discrepancies. Under a human-like recombination rate (*r* = 10^−8^), linked selection has little effect on the inferred spectra. The simulations assume mutations and recombination events are uniformly distributed along the genome, rather than clustered into functional regions as in real genomes.

**Figure C14:**
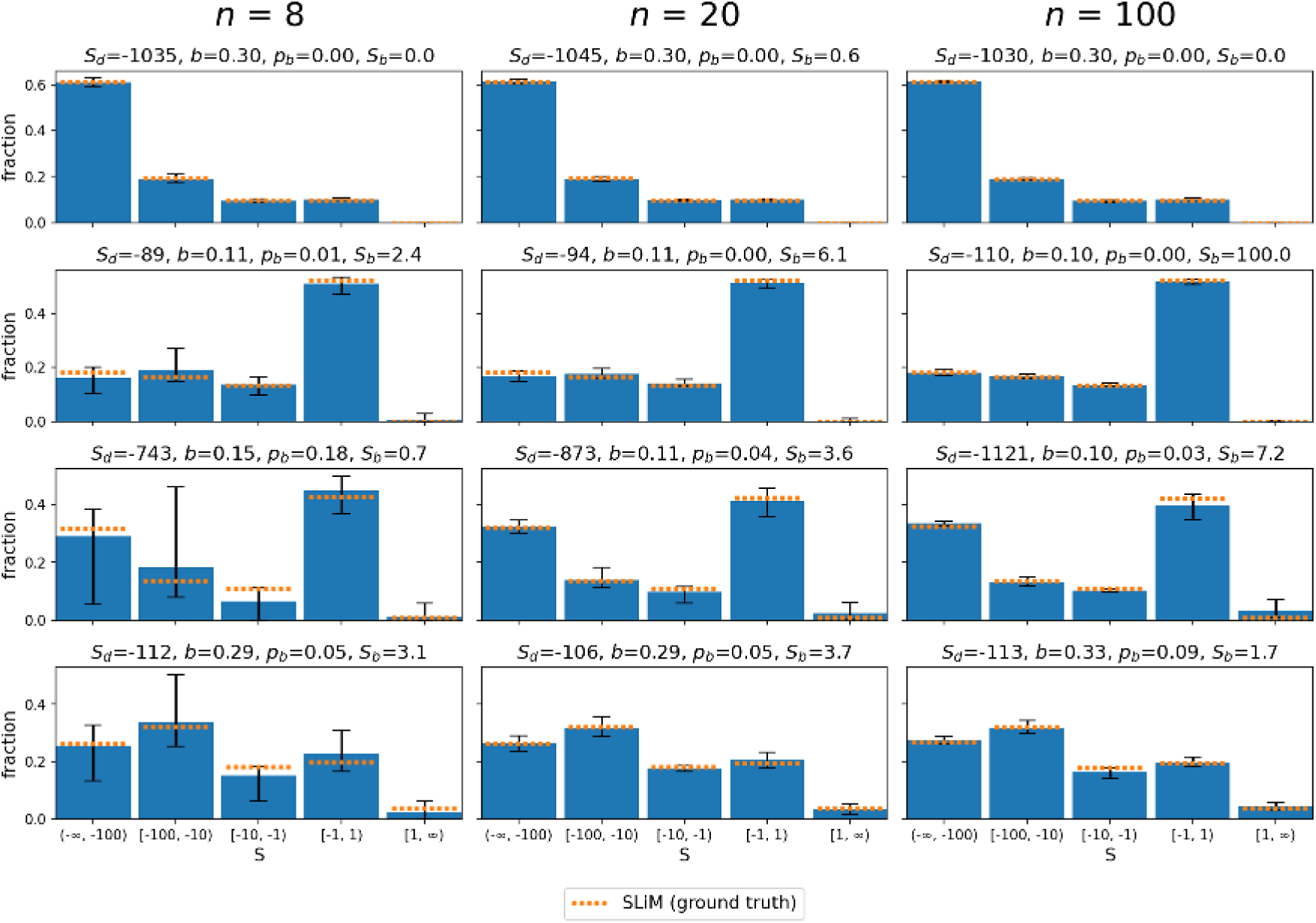
Inferred DFEs across varying SFS sample sizes under the constant population size scenario. Orange lines mark the simulated DFE, converted to population-scaled coefficients using the *N_e_* estimated from each simulation’s *neutral* spectrum. Rows correspond to the four simulated DFEs introduced above, and bars show the inferred fraction in each interval with 90% bootstrap confidence intervals. The deleterious classes are recovered accurately at every sample size, whereas the beneficial parameters are poorly determined at *n* = 8 and improve as the sample size increases.

## Notes

https://github.com/Sendrowski/PrimateDFE

### Competing Interest Statement

The authors have declared no competing interest.

### Summary of Updates

Accepted for publication in GENETICS after a second round of review. A control restricting the spectra to the ortholog groups shared by all reference genomes was added, with a new Results subsection and three new appendix figures. The Results were also reorganised into subsections.

