## Supplementary Tables X for "Comparison of the Distribution of Fitness Effects Across Primates" (from another version of this preprint: version 1, 2026-03-25)

**Supplementary Tables X:** Effect of regression framework and DFE parameterization on estimates.
Intercepts and slopes associated with generation time (`gen_time`) and `log10(Ne)` are reported. For each Grafen PGLS model, we additionally report ρ, the branch length scaling parameter, inferred from the data.
Code availability: https://github.com/tbata/PLS_DFE

**DFE model: discrete bins (allowing for a proportion of beneficial mutations)**

| Regression Model | AIC | R^2 | Intercept | p(Intercept) | log10Ne | p(log10Ne) | gen_time | p(gen_time) |
|---|---|---|---|---|---|---|---|---|
| OLS | -129.38 | 0.825 | 0.194 | 0.0406 | 0.106 | 8e-07 | -0.003 | 0.0281 |
| PGLS Brownian | -129.35 | 0.822 | 0.016 | 0.831 | 0.157 | 4.46e-11 | -0.005 | 0.00758 |
| PGLS Grafen (rho=0.37) | -128.96 | 0.825 | 0.011 | 0.902 | 0.147 | 3.85e-09 | -0.003 | 0.071 |

**DFE model: discrete bins (deleterious mutations only)**

| Regression Model | AIC | R^2 | Intercept | p(Intercept) | log10Ne | p(log10Ne) | gen_time | p(gen_time) |
|---|---|---|---|---|---|---|---|---|
| OLS | -140.31 | 0.784 | 0.152 | 0.0516 | 0.103 | 4.96e-08 | 0.001 | 0.549 |
| PGLS Brownian | -133.77 | 0.767 | 0.047 | 0.511 | 0.134 | 2.63e-10 | -0.001 | 0.526 |
| PGLS Grafen (rho=0.10) | -138.8 | 0.783 | 0.096 | 0.229 | 0.115 | 2.43e-08 | 0.001 | 0.45 |

**DFE model: gamma + exponential model (allowing for a proportion of beneficial mutations)**

| Regression Model | AIC | R^2 | Intercept | p(Intercept) | log10Ne | p(log10Ne) | gen_time | p(gen_time) |
|---|---|---|---|---|---|---|---|---|
| OLS | -132.95 | 0.803 | 0.231 | 0.0113 | 0.098 | 1.12e-06 | -0.002 | 0.0884 |
| PGLS Brownian | -134.47 | 0.791 | 0.094 | 0.194 | 0.144 | 4.57e-11 | -0.006 | 0.00289 |
| PGLS Grafen (rho=0.33) | -132.43 | 0.803 | 0.098 | 0.26 | 0.128 | 2.13e-08 | -0.002 | 0.129 |

**DFE model: gamma model (deleterious mutations only)**

| Regression Model | AIC | R^2 | Intercept | p(Intercept) | log10Ne | p(log10Ne) | gen_time | p(gen_time) |
|---|---|---|---|---|---|---|---|---|
| OLS | -147.08 | 0.844 | 0.199 | 0.00598 | 0.097 | 1.65e-08 | -0.001 | 0.309 |
| PGLS Brownian | -144.8 | 0.835 | 0.115 | 0.0611 | 0.124 | 2.53e-11 | -0.003 | 0.0724 |
| PGLS Grafen (rho=0.23) | -147.25 | 0.844 | 0.129 | 0.0655 | 0.113 | 2.35e-09 | -0.001 | 0.364 |
